# Evolution of chemosensory and detoxification gene families across herbivorous Drosophilidae

**DOI:** 10.1101/2023.03.16.532987

**Authors:** Julianne N. Pelaez, Andrew D. Gloss, Benjamin Goldman-Huertas, Bernard Kim, Richard T. Lapoint, Giovani Pimentel-Solorio, Kirsten I. Verster, Jessica M. Aguilar, Anna C. Nelson Dittrich, Malvika Singhal, Hiromu C. Suzuki, Teruyuki Matsunaga, Ellie E. Armstrong, Joseph L.M. Charboneau, Simon C. Groen, David H. Hembry, Christopher J. Ochoa, Timothy K. O’Connor, Stefan Prost, Sophie Zaaijer, Paul D. Nabity, Jiarui Wang, Esteban Rodas, Irene Liang, Noah K. Whiteman

## Abstract

Herbivorous insects are exceptionally diverse, accounting for a quarter of all known eukaryotic species, but the genetic basis of adaptations that enabled this dietary transition remains poorly understood. Many studies have suggested that expansions and contractions of chemosensory and detoxification gene families – genes directly mediating interactions with plant chemical defenses – underlie successful plant colonization. However, this hypothesis has been challenging to test because the origins of herbivory in many lineages are ancient (>150 million years ago [mya]), obscuring genomic evolutionary patterns. Here, we characterized chemosensory and detoxification gene family evolution across *Scaptomyza,* a genus nested within *Drosophila* that includes a recently derived (<15 mya) herbivore lineage of mustard (Brassicales) specialists and carnation (Caryophyllaceae) specialists, and several non-herbivorous species. Comparative genomic analyses revealed that herbivorous *Scaptomyza* have among the smallest chemosensory and detoxification gene repertoires across 12 drosophilid species surveyed. Rates of gene turnover averaged across the herbivore clade were significantly higher than background rates in over half of the surveyed gene families. However, gene turnover was more limited along the ancestral herbivore branch, with only gustatory receptors and odorant binding proteins experiencing strong losses. The genes most significantly impacted by gene loss, duplication, or changes in selective constraint were those involved in detecting compounds associated with feeding on plants (bitter or electrophilic phytotoxins) or their ancestral diet (yeast and fruit volatiles). These results provide insight into the molecular and evolutionary mechanisms of plant-feeding adaptations and highlight strong gene candidates that have also been linked to other dietary transitions in *Drosophila*.

## Introduction

The origin of land plants over 500 million years ago presented a new niche for early insects to colonize (Southwood 1972). The intimate relationship between plants and insects has since generated one of the most ecologically and evolutionarily dominant groups in Earth’s history: the herbivorous insects. Herbivorous insects account for over a quarter of all known eukaryotic species and help form the basis of terrestrial food webs (Strong *et al*. 1984; Farrell 1998; Bernays 1998). It has long been hypothesized that the diversity of herbivorous insects emerged as a result of co-diversification processes with their host plants (Mitter *et al*. 1988; Farrell 1998; Marvaldi *et al*. 2002).

As proposed by Ehrlich and Raven (1964), herbivores can diversify by specializing on plants bearing the same chemical defenses through adaptations to chemical defenses (in one or a few plant families), that is, until new plant defenses evolve, allowing plants to escape and diversify under this release from herbivore pressure in this ‘coevolutionary arms-race’ (Ehrlich and Raven 1964). This theory of coevolution, commonly referred to as ‘escape and radiate’ (Thompson 1988, 1994), has inspired much of the research on insect-plant interactions throughout the last several decades, (Heidel-Fischer and Vogel 2015; Simon *et al*. 2015; Vertacnik and Linnen 2017). As a result, we have learned that genes that mediate interactions with plant secondary compounds, such as chemosensory and detoxification genes, likely underlie adaptive mechanisms for plant colonization (Ehrlich and Raven 1964; Berenbaum 1983; Li *et al*. 2003a; Futuyma and Agrawal 2009; Edger *et al*. 2015). Despite significant progress towards understanding the genetic basis of herbivore-plant interactions, the evolutionary processes shaping these large, complex and rapidly evolving gene families are still not fully understood.

A principal source of the functional genetic variation that underlies dietary novelty in herbivorous arthropods arises from extensive gene family evolution. For example, the spider mite (*Tetranychus urticae*) and diamondback moth (*Plutella xylostella*) genomes harbor expanded gene families encoding enzymes involved in the detoxification of plant secondary compounds they encounter (Grbić *et al*. 2011; You *et al*. 2013; Dermauw *et al*. 2013). Similarly, genes encoding gustatory receptors (GRs) involved in host finding have experienced extensive lineage-specific duplications in butterflies (Briscoe *et al*. 2013). Despite the identification of gene-family expansions and contractions in herbivorous insects, (McBride 2007; Edger *et al*. 2015; Johnson *et al*. 2018), isolating herbivory as the cause remains controversial, as these changes may have resulted from subsequent specialization occurring over a hundred million years. For example, the most diverse extant herbivore lineages – butterflies and moths (Lepidoptera), as well as leaf beetles, weevils and close relatives (Phytophaga) – arose in the late Paleozoic and early Mesozoic, respectively (Wiens *et al*. 2015; Kawahara *et al*. 2019). Parsing herbivore-specific effects from those resulting from specialization is particularly challenging given the prominent role of specialization in driving herbivore diversification rates. This is strongly supported by associations between host shifts and speciation events and by the higher species richness found among specialist herbivores compared to generalist herbivores (Futuyma and Agrawal 2009; Forister *et al*. 2015).

While it is unclear whether specialization on specific plant taxa evolves during or after the evolution of herbivory (Bernays 1998), many phytophagous insects nonetheless exhibit phylogenetic conservatism: associating with the same plant taxa for many millions of years (Futuyma and Agrawal 2009). Comparative genomic studies examining younger herbivore lineages would thus allow for a more refined analysis to identify herbivore-associated changes from those arising in response to specialization or other evolutionary forces (Gardiner *et al*. 2008; Yassin *et al*. 2016). Furthermore, most herbivore lineages lack the functional genetic tools necessary to examine the implications of these copy number changes. Here, we addressed these limitations by studying the evolution of herbivory within Drosophilidae using a comparative genomics approach.

Although larvae of most Drosophilidae species retain the ancestral habit of feeding on decaying plant tissue and associated microbes, herbivory has evolved several times in the lineage (Okada and Sasakawa 1956; Jones 1998; Whiteman *et al*. 2011; Maunsell *et al*. 2017; Durkin *et al*. 2021). A major clade of herbivorous species in the family are members of the genus *Scaptomyza*, a monophyletic lineage of ∼272 species and 21 subgenera (O’Grady and Desalle 2008). *Scaptomyza* spp. are nested within the paraphyletic subgenus *Drosophila,* which also includes Hawaiian *Drosophila* and the virilis-repleta radiation (Fig. 1a) (O’Grady and Desalle 2008; Lapoint *et al*. 2013; Katoh *et al*. 2017; O’Grady and DeSalle 2018; Church and Extavour 2022). Herbivorous *Scaptomyza* are found across the Holarctic, and *S. flava* in particular has been introduced into New Zealand, where it is a pest (Martin 2004). DNA barcoding revealed that the clade of herbivores may be a cryptic radiation, with the divergence of eight species in North America within the last ∼10 million years (Fig. 1b) (Lapoint *et al*. 2013; Peláez *et al*. 2022). This is similar in species richness to the *D. melanogaster* subgroup worldwide (nine species within ∼12 million years) (David *et al*. 2007).

**Figure 1.**
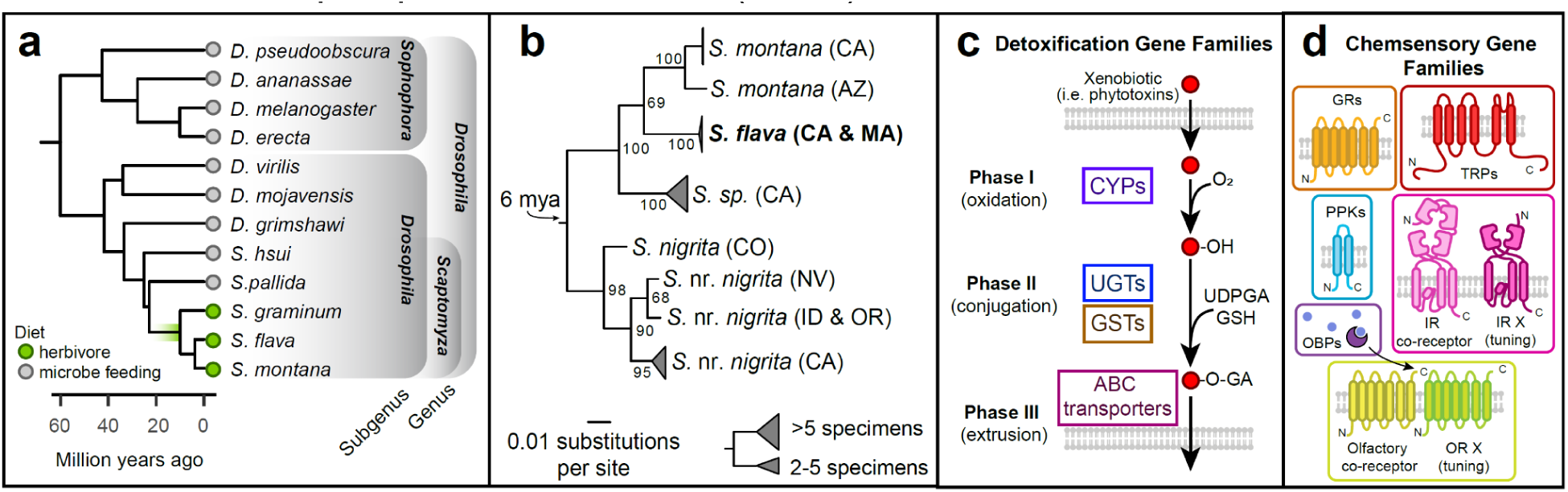
Gene families at the interface of plant-herbivore interactions. (a) Phylogenetic placement of herbivorous *Scaptomyza* within the paraphyletic genus *Drosophila* (Matsunaga *et al*. 2022). (b) Maximum likelihood nucleotide phylogeny built using *COI* sequences from North American *Scaptomyza* collected feeding on mustard plants (Brassicales spp.). Individuals with < 1% pairwise nucleotide divergence were collapsed into clades. Collection localities indicated by two-letter state abbreviations. For the current study, the genome of an *S. flava* line from MA, USA (bolded) was sequenced and assembled. Sequences and complete phylogeny are included in Supporting Dataset 4. Divergence time estimated from Whiteman et al. 2012. (c) Detoxification of xenobiotics/plant secondary metabolites (phytotoxins) involves three phases: oxidation/reduction by cytochrome P450s (CYPs), enzymatic conjugation by UDP glucuronosyltransferases (UGTs) or glutathione S-transferases (GSTs), and excretion/transport out of the cells by ABC transporters. (d) Insects detect environmental compounds mostly with six major chemosensory gene families: gustatory receptors (GRs), odorant binding proteins (OBPs), olfactory receptors (ORs), degenerin/epithelial sodium channels (pickpockets PPKs), ionotropic receptors (IRs), and transient receptor potential channels (TRPs).

The herbivorous *Scaptomyza* species have become models for the evolution of herbivory, and are an ideal group to test hypotheses about gene family evolution for several reasons. Most specialize on plants in the Brassicales (mustards and their relatives) and can be reared on the genetic model plant, *Arabidopsis thaliana* (Whiteman *et al*. 2011, 2012). Genetic dissection of adaptive traits in this lineage (Gloss *et al*. 2014; Goldman-Huertas *et al*. 2015; Peláez *et al*. 2022) has been enabled by the rich knowledge of gene function in *D. melanogaster*, a growing number of high-quality genomes across the drosophilid phylogeny (Drosophila 12 Genomes Consortium *et al*. 2007; Kim *et al*. 2021), strong phylogenetic frameworks (O’Grady and DeSalle 2018; Finet *et al*. 2021), and the ability to test hypotheses using genetic tools from both *Drosophila* and *Arabidopsis* (Groen and Whiteman 2016).

A critical advantage of studying *Scaptomyza* is that the genus encompasses species exhibiting varying degrees of specialization on different plant families: *S. graminum,* which specializes on plants of the Caryophyllaceae family (pinks or carnations), and *S. flava* as well as *S. montana,* which are largely specialists on Brassicales. The Brassicales harbor non-toxic glucosinolates that are the precursors to toxic mustard oils. While *S. flava* feeds on numerous genera of Brassicales, *S. montana* is more specialized, showing a strong preference for plants with indole glucosinolates (Gloss *et al*. 2017). *S. flava* has also been found to attack common pea plants (*Pisum sativum,* Fabaceae) and has further expanded its host range to some Caryophyllaceae in New Zealand (Martin 2004, 2012). Genetic changes shared by these three species are more likely to be associated with the initial transition to herbivory, rather than their respective subsequent specializations. Comparisons with closely related non-herbivores, such as *S. pallida* and *S. hsui*, each from different subgenera (*Parascaptomyza* and *Hemiscaptomyza*, respectively), also allow us to differentiate herbivore-specific changes from those that arose earlier along the *Scaptomyza* lineage. Lastly, non-herbivore specialists outside *Scaptomyza* can be used to further polarize patterns of specialization from those linked to herbivory. For instance, genome assemblies are available for *D. erecta*, a specialist on screwpine fruit (*Pandanus* spp.), and *D. mojavensis* that has populations specializing on rotting prickly pear cactus (*Opuntia littoralis*) (Rio and Others 1983; Pfeiler *et al*. 2005).

Given the salience of gene expansions and contractions associated with herbivorous insects, we hypothesized that genes involved in sensing or detoxifying plant chemical defenses would experience the greatest copy number changes. Specifically, in accordance with the neural limitation theory, we expected herbivores to lose numerous chemosensory genes, to streamline neural processing in the face of choosing between many host plants (Bernays 2001) and also their ancestral diet. We also expected to find duplicated genes with signatures of rapid protein evolution, which could have enabled early herbivores to gain new chemosensory or detoxification functions through neofunctionalization or subfunctionalization (Ohno 1970; Lynch and Conery 2000).

To address these hypotheses and predictions, we sequenced and assembled a high-quality genome sequence of *S. flava* to analyze alongside publicly available genome sequences of two herbivorous species (*S. montana* and *S. graminum*), two non-herbivores (*S. pallida* and *S. hsui*), and seven non-herbivorous *Drosophila*, across a phylogenetic gradient from the herbivores (Fig. 1a). Following the curation of the major chemosensory and detoxification gene families (those in Fig. 1c-d), we then used maximum likelihood (ML) methods to test whether rates of gene gain and loss in herbivore branches differed from background branches. We next generated codon-based, ML models to identify genes with signatures of changing selective regimes along the branch at the base of the herbivore clade. We also evaluated whether our results may be explained by demographic events in recent history. We found that the most dramatic changes occurred in chemosensory genes involved in sensing yeast fermentation products/fruit volatiles and bitter/toxic plant chemical defenses.

## Materials and Methods

### *S. flava* genome sequencing, assembly, and annotation

Full details of these methods can be found in the Supplemental Methods.

#### PacBio and Dovetail HiC libraries and sequencing

Sequence data for our main *S. flava* assembly (sfla_v2) were generated from a partially inbred laboratory colony. The colony was founded from >150 larvae collected near Dover, NH, USA, and subsequently maintained for several years in the laboratory. 300 male flies were flash frozen and stored at −80°C. SMRTbell libraries (∼20kb) for PacBio Sequel were constructed using SMRTbell Template Prep Kit 1.0 (PacBio, Menlo Park, CA, USA) using the manufacturer’s recommended protocol. Sequencing was performed on two PacBio Sequel SMRT cells. A Dovetail HiC library was prepared in a similar manner as described previously (Lieberman-Aiden *et al*. 2009). DNA was sheared to ∼350 bp mean fragment size, and sequencing libraries were generated using NEBNext Ultra enzymes and Illumina-compatible adapters. The libraries were sequenced on an Illumina HiSeqX to produce 380 million 2x150 bp paired-end reads, approximately 30x sequence coverage.

#### Illumina library and sequencing

Additional Illumina sequence data was used to polish the PacBio assembly. Sequences were generated from a laboratory population initially collected in Belmont, MA, USA in 2008 that was inbred through 10 generations of single-pair matings. Paired-end 180 bp and 300 bp insert libraries and 3 kbp and 5 kbp mate-pair libraries from female flies were sequenced with 100 bp read length on an Illumina Hiseq 2000 at the University of Arizona. Reads were quality filtered and Illumina TruSeq3 adapters were removed using Trimmomatic v0.35 (Bolger *et al*. 2014) with the following parameters: “LEADING:10 TRAILING:10 SLIDINGWINDOW:4:15 MINLEN:99”.

#### Genome assembly and scaffolding

The *S. flava* genome was assembled using the long-read hybrid assembly pipeline described in Kim et. al (2021), which has been shown to produce highly complete genome assemblies for drosophilid flies resulting in very few coding sequence indels (Kim *et al*. 2021). We generated an initial draft assembly with Flye 2.9 (Kolmogorov *et al*. 2019), and identified and removed duplicated haplotypes (haplotigs) with purge_dups (Guan *et al*. 2020). We polished the draft assembly using the PacBio reads with one round of Racon (Vaser *et al*. 2017), then polished further using Illumina reads with one round of Pilon (Walker *et al*. 2014), only fixing base-level errors. The fully polished assembly was scanned for contaminant sequences using NCBI BLAST (Johnson *et al*. 2008) and BlobTools (Laetsch and Blaxter). Repetitive sequences in the assembly were identified with RepeatModeler2 (Flynn *et al*. 2020). To integrate the long-range assembly information generated with HiC, we scaffolded the assembly with an initial assembly generated by Dovetail (details included in the Supplemental Methods). Specifically, we soft-masked both genomes using the repeat library generated in the previous step, using RepeatMasker (Smit *et al*. 2013). Then, we created a whole-genome alignment with Progressive Cactus (Armstrong *et al*. 2020) and used the RagOut reference-based scaffolder (Kolmogorov *et al*. 2014) to scaffold the new genome. The genome assembly of *S. flava* was scaffolded into 1,252 scaffolds covering 315.4 Mbp (N50 = 32.966 Mb) with a maximum length of 85.98 Mbp and with 460 gaps.

#### Comparative annotation of *Scaptomyza* genomes

Gene annotations previously created for an Illumina-only *S. flava* assembly (sfla_v1) were transferred to the assemblies of *S. graminum, S. hsui, S. pallida, S. montana*, and to the newest *S. flava* assembly (sfla_v2), using whole-genome Progressive Cactus alignments and the Comparative Annotation Toolkit (CAT, (Fiddes *et al*. 2018)). Details of these methods are in the Supplemental Methods.

### Gene model curation & orthology inference

An iterative curation strategy was used to identify the complement of chemosensory and detoxification genes in four published *Drosophila* genomes (*D. melanogaster, D. virilis, D. mojavensis,* and *D. grimshawi) (Drosophila 12 Genomes Consortium et al. 2007)*, four published *Scaptomyza* genomes (*S. pallida, S. hsui, S. graminum,* and *S. montana*) (Kim *et al*. 2021), and in our *S. flava* assembly (sfla_v2). Genome assembly versions are listed in Table S1, along with which species were included in each analysis. Gene curation included six chemosensory gene families (gustatory receptors, GRs; ionotropic receptors, IRs; olfactory receptors, ORs; odorant binding proteins, OBPs; pickpocket proteins, PPKs; transient receptor potential channels, TRPs) and three detoxification gene families (Cytochrome P450s, CYPs; Glutathione *S*-transferases, GSTs; UDP-glycosyltransferases, UGTs). First, all protein sequences in each family from *D. melanogaster* were queried against the assembled genomes and annotated proteomes of *D. virilis, D. mojavensis,* and *D. grimshawi*, using BLASTP (e-value cutoff < 1e-3) (Altschul *et al*. 1997). The resulting collection of genes was then queried against the automated annotations for *Scaptomyza* species (e-value cutoff < 1e-3). We iteratively ran BLAST searches using the identified genes from each species as queries against their genome assemblies until no new genes were identified. To validate putatively lost genes, we performed an additional TBLASTN search (e-value cutoff < 1e-4). Genes were considered truly absent if this yielded no hits. Gene models significantly deviating in length from *D. melanogaster* orthologs were manually inspected, and corrected using aligned homologous sequences to the annotated genes in other species. Additional validation steps we implemented are described in the Supplementary Methods. Nucleotide sequences were aligned in Geneious v.10.2.6 using MUSCLE (Edgar 2004) and manually inspected. All gene coordinates are provided in Supporting Dataset 1.

To verify correct orthology assignments, ML gene trees were constructed from sequences from *D. grimshawi* and all *Scaptomyza*, using RAxML with default settings (Stamatakis 2006). Genes were binned into orthologous groups if they formed a clade with > 70 bootstrap support. For poorly supported clades (bootstrap support < 70), orthology groups were assigned on the basis of previous orthology assignments (Low *et al*. 2007; Almeida *et al*. 2014; Good *et al*. 2014). Gene trees were midpoint rooted and visualized in iTOL v6 (Letunic and Bork 2021).

### Gene family size evolution

To identify chemosensory or detoxification gene families with rapid expansions and/or contractions among the herbivore lineages, we used the program Computational Analysis of gene Family Evolution (CAFE) v4.2.1 (Han *et al*. 2013), which models a birth-death process of gene gain and loss across a species tree. In addition to the nine species mentioned above, we included three species of the subgenus *Sophophora* (*D. pseudoobscura*, *D. ananassae*, and *D. erecta*) to improve background rate estimates. We imported gene models for these species from published orthology annotations (Good et al. 2014; Low et al. 2007; Almeida et al. 2014). As input, we provided a matrix of gene counts for each orthologous gene cluster (Supporting Dataset 2). Orthology groups were merged until there was at least one homologous gene reconstructed at the base of the phylogeny as the analysis assumes at least one ancestral gene per group. We used a published time-calibrated phylogeny of drosophilids (Matsunaga *et al*. 2022), which included all our species of interest, with the exception of *S. montana*, which was grafted onto this tree using a reported divergence time estimate from *S. flava* (Peláez *et al*. 2022).

First, we generated models of the average turnover rate (λ) (the ML estimate of cumulative rate of gains and losses per gene per unit time). For each gene family, we compared a null model with a single λ rate estimated for all branches to a model with two rates: one rate estimated for a foreground branch, another for the remaining background branches, where the foreground branch was the internal (ancestral) branch leading to all herbivores, a terminal branch leading to one of the 12 species, or the entire clade of herbivores. All models were run in triplicate, and the iteration with the highest ML probability was retained. To test whether a two-rate model was significantly better than the single-rate model, we performed likelihood ratio tests (LRTs). *P*-values were adjusted at a false discovery rate of 1% using the qvalue package in R (Storey 2008). Using the same methods, we also generated models estimating gains and losses (λ, µ) separately, rather than as a single parameter.

To test whether expansion and contraction rates of chemosensory and detoxification gene families deviated from genome-wide rates, we performed the same analyses on 200 randomly chosen orthology groups. We identified these gene sets using the OrthoVenn2 server (Wang *et al*. 2015; Xu *et al*. 2019), a web-based orthology assignment tool based on OrthoMCL (Li *et al*. 2003b). We uploaded genome-wide proteomes for the five *Scaptomyza* species, and used protein sequences for the remaining seven species available through OrthoVenn2 (Ensembl database, release January 2019). Default parameters were used with an e-value cutoff of 0.05 and an inflation value of 1.5. Clusters were randomly chosen that summed to ∼200 genes per species.

### Molecular evolutionary analysis

To identify genes that experienced changes in selective pressure along the ancestral branch at the base of the herbivorous species, we used codon-based models of evolution (*codeml* program) in the Phylogenetic Analysis by Maximum Likelihood (PAML) package (Yang 2007). These models estimate the non-synonymous (dN) to synonymous (dS) substitution rate ratio (ω = dN/dS), wherein protein-coding genes experiencing positive directional selection may accumulate a significant number of amino acid substitutions resulting in dN/dS>1, those evolving neutrally dN/dS ≈ 1, and those experiencing negative or purifying selection dN/dS<1.

For each group of orthologous genes, nucleotide sequences from nine taxa (*S. flava, S. montana, S. graminum, S. pallida, S hsui, D. grimshawi, D. mojavensis, D. virilis*, and *D. melanogaster*) were aligned in Geneious v.10.2.6 using MAFFT v7.450 translation alignment with default settings (Katoh *et al*. 2002; Katoh and Standley 2013). A species tree was generated for these analyses by summarizing the 100 ML-generated gene trees published in (Kim *et al*. 2021), using ASTRAL v.5.5.9 (Zhang *et al*. 2018) with default settings. For genes with multiple paralogs, we estimated gene trees using FastTreeMP v2.1 with a GTR+gamma nucleotide substitution model and 100 bootstrap replicates (Price *et al*. 2010).

We employed branch, branch-site, and clade models within PAML. The branch model allows dN/dS to vary across branches, assuming the same rates across sites (Yang 1998). To assess changes in selection pressures across branches, we compared a “two-ratio” branch model, where dN/dS is estimated separately for a specified foreground branch and background branches (model = 2, NSites = 0), against a null model (“M0”), which estimates a single dN/dS rate over all branches (model = 0, NSsites = 0). This test specifically asks whether foreground and background dN/dS rates differ, which could be attributed to positive selection, reduced purifying selection, or higher purifying selection. To test whether the foreground branch has experienced positive selection (dN/dS>1), we compared the two-ratio model to a “constrained two-ratio” model, which includes the same parameters but ω is fixed to 1 (model = 2, NSites = 0, fix_omega = 1).

Branch models may be unable to detect positive selection if it acts only at a few sites with the remaining sites likely to remain under purifying selection. Therefore, we also used branch-site models, which more realistically model protein evolution, allowing dN/dS to vary across both codons and branches (Yang and Nielsen 2002; Zhang *et al*. 2005). We compared the “branch-site model A” (model = 2, NSsites = 2, fix_omega = 0) against a null model in which ω_2_ is fixed to 1 (model = 2, NSsites = 2, fix_omega = 1). This comparison offers a direct test for positive selection.

Finally, we used the “Clade model C” (CmC) to estimate changing selection pressures when genes exhibited numerous paralogs within species (Weadick and Chang 2012). Based on branch-site models, CmC models allow dN/dS to vary in a proportion of sites between two gene clades, and can detect more subtle divergent selection, particularly after duplication events (Bielawski and Yang 2004; Weadick and Chang 2012). We tested for divergent selection by comparing the CmC model (model = 3, NSsites = 2, fixed = 0) against the null model “M2a_rel” (model = 0, NSsites = 22, fixed = 0), which has the same number of site classes as the CmC model but does not vary across branches. To test for positive selection on the ancestral herbivore branch, we compared the CmC model against a null model in which the ω of the divergent site class is constrained to 1 (model = 3, NSsites = 2, fixed = 1) (Van Nynatten *et al*. 2015). For clades with foreground divergent sites significantly deviating from the null expectation, we then performed branch and branch-site tests to test which set of paralogs experienced divergent selection.

For all PAML analyses, model comparisons were made using LRTs with a χ^2^ distribution and one degree of freedom. *P*-values were adjusted at a false discovery rate of 5% using the qvalue package in R (Storey 2008).

## Results

### *S. flava* genome assembly and annotation

To complement existing genome assemblies of non-herbivorous and herbivorous drosophilids, we sequenced, assembled, and annotated the genome of *Scaptomyza flava*, a leaf-mining specialist on mustards. Our main assembly (sfla_v2) contained 781 scaffolds with an N50 of 31.83 Mb and an assembly size of 331.7 Mb (Table S2). Using gene models from other *Drosophila* species and transcriptome sequences across *S. flava* life stages, we generated annotations for the *S. flava* assembly sfla_v1, which were carried over to our main assembly sfla_v2, amounting to 12,365 predicted genes. We performed the same carry-over procedures on four other *Scaptomyza* species (*S. pallida, S. hsui, S. montana,* and *S. graminum*) to facilitate annotation of their chemosensory and detoxification gene families. The completeness of the assemblies was assessed by comparing the assembly against the BUSCO dipteran database. The genome assembly of *S. flava* is near complete, as are the assemblies of the four other *Scaptomyza* species with values comparable to the published assemblies of the seven other *Drosophila* species included in subsequent analyses (*D. melanogaster, D. erecta, D. ananassae, D. pseudoobscura, D. mojavensis, D. virilis,* and *D. grimshawi*) (Fig. S1a). Approximately 98.8% of complete BUSCO gene models (98% single-copy and 0.8% duplicated) were identified from the sfla_v2 assembly, 0.4% were found fragmented, and 0.8% were not found. BUSCO scores were also high for the genome-wide automated annotations in all *Scaptomyza* species (Fig. S1b). The completeness of the genome assemblies and automated annotations provided confidence that we would be able to reliably assess the evolution of our gene families of interest.

### Contractions and expansions of chemosensory and detoxification gene families

We analyzed the evolution of chemosensory and detoxification gene families to determine if rates of gain, loss, and/or turnover (cumulative gains and losses) were significantly different between non-herbivorous species and herbivorous *Scaptomyza* species. We used CAFE, a program that makes maximum likelihood estimates of gene copy-number evolutionary rates, (Han *et al*. 2013) and included nine non-herbivorous and three herbivorous species (all species in Fig. 1a), all of which use a range of different feeding substrates with varying degrees of specialization. While we tested whether rates were significantly higher along terminal herbivore branches and across the entire herbivore clade, compared to the remainder of the tree, we were particularly interested in significantly higher copy number changes along the ancestral herbivore branch. We curated six chemosensory gene families (gustatory receptors, GRs; ionotropic receptors, IRs; olfactory receptors, ORs; odorant binding proteins, OBPs; pickpocket proteins, PPKs; transient receptor potential channels, TRPs) and three detoxification gene families (cytochrome P450s, CYPs; glutathione *S*-transferases, GSTs; UDP-glycosyltransferases, UGTs) (gene coordinates: Supporting Dataset 1, RAXML gene trees: Fig. S2-10).

Overall, herbivorous species had smaller repertoires of detoxification and chemosensory genes than non-herbivorous species. (Fig. 2b, Table S3). The rate of gene turnover was significantly higher along the ancestral herbivore branch than the background branches when all chemosensory genes were considered (λ_anc_herb_=0.005; λ_bkgrd_=0.002; q-value=0.005) but not for all detoxification genes (λ_anc_herb_=0.005; λ_bkgrd_=0.002; q-value>0.05) (Fig. 2c, Table S4). When individual gene families were analyzed, the ancestral herbivore branch only experienced significantly higher rates of turnover among OBPs. We also estimated rates of gene duplication and loss separately, finding that along the ancestral herbivore branch the gene duplication rate was not significantly higher for any gene family compared to background rates (Fig. 2d, Table S5).

**Figure 2.**
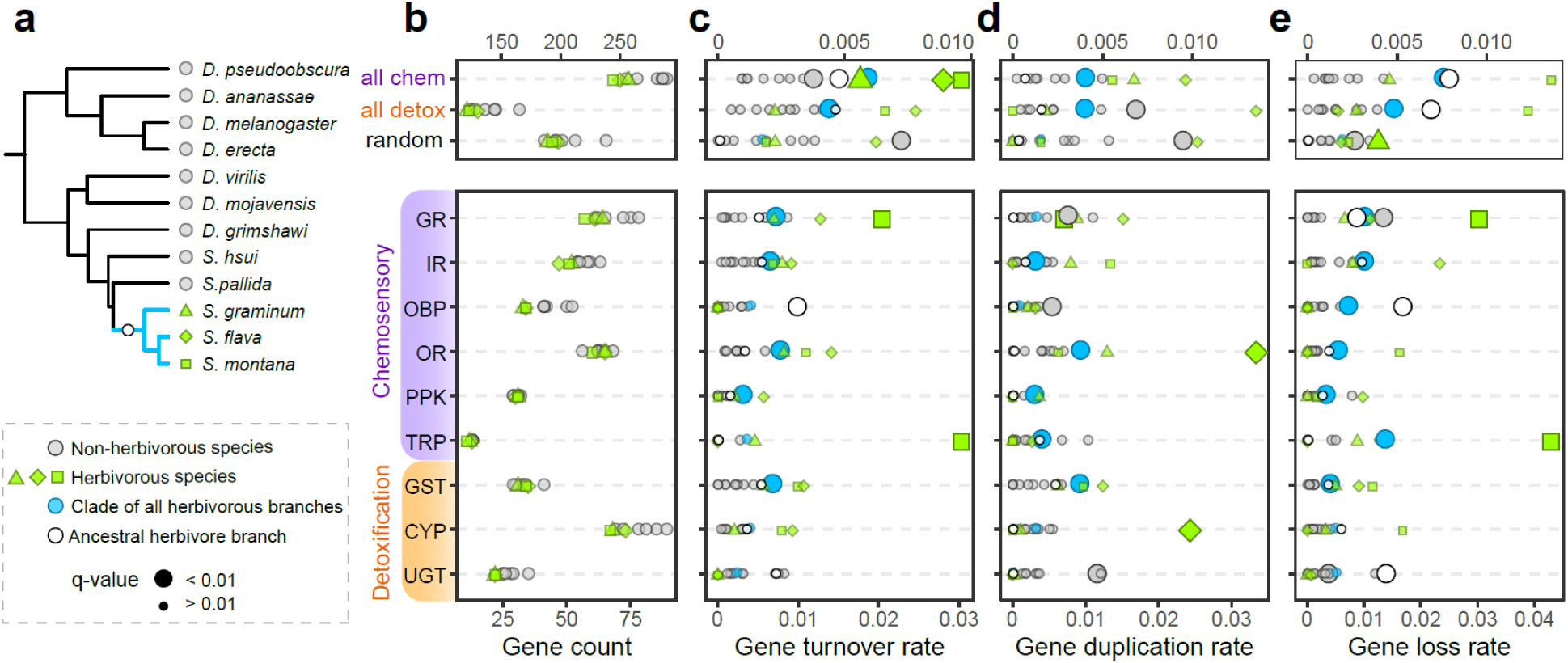
The clade of *Scaptomyza* herbivores (but not their ancestral lineage) exhibit elevated gene turnover within many chemosensory and detoxification gene families. (a) Phylogeny of drosophilids included in CAFE analyses: three herbivorous and nine non-herbivorous *Drosophila*. (b) Gene counts per gene family. Evolution of gene family size was estimated by maximum likelihood in CAFE, where the rates of (c) gene turnover, λ, (cumulative gains and losses), (d) gene duplication, λ, and (e) gene loss, µ, were estimated for each species. Points represent the rate of a foreground branch or clade from models in which it was allowed to evolve at a rate separate from background branches. Each model was compared to a null model (single rate estimated for the entire phylogeny) using a likelihood ratio test at an analysis-wide false discovery rate of 1% (*q* < 0.01). Foreground branches with significantly higher rates than the rest of the phylogeny are indicated by larger shapes. Full details can be found in Tables S3-5. All chem = all chemosensory genes. all detox = all detoxification genes. Random = random set of 200 orthology groups.

The gene loss rate, however, was significantly higher for all chemosensory genes (λ_anc_herb_=0.008; λ_bkgrd_=0.002; q-value<0.001) and detoxification genes (λ_anc_herb_=0.007; λ_bkgrd_=0.002; q-value=0.003) (Fig. 2e, Table S5). Specifically, higher rates of gene loss along the ancestral herbivore branch were found among GRs, OBPs, and UGTs. The CAFE analysis also allowed us to identify specific genes that experienced expansions and contractions (Table S6). Significant contractions along the ancestral herbivore branch included OBPs (*Obp18a, Obp58b,* and *Obp58c*) and *Or22a*, an ester-sensitive, yeast volatile receptor. The only expansion was of *Ir67a*, which was duplicated in the ancestral herbivorous lineage.

While we found limited copy number changes along the ancestral herbivore branch, rates of gene turnover, duplication, and loss estimated across the entire herbivore clade were significantly higher than background rates for numerous gene families (gene turnover: GRs, IRs, ORs, PPKs, GSTs (Fig. 2c); duplication: IRs, ORs, PPKs, TRPs and GSTs (Fig. 2d); loss: GRs, IRs, OBPs, ORs, PPKs, TRPs and GSTs (Fig. 2e)). We confirmed that it is unlikely that these elevated rates among the herbivorous species could be attributed to the longer branch length of the ancestral branch preceding the herbivore clade. We performed another set of CAFE analyses to compare rates between two clades of similar age: the clade of *D. melanogaster* and *D. erecta* (28 million years old) versus the clade of *S. flava* and *S. graminum* (23 million years old). This confirmed that the former pair showed no gene families evolving at higher rates, whereas the clade of *S. flava* and *S. graminum* still showed elevated rates of turnover, loss and gain (Fig. S11).

Based on the RAxML trees for each gene family (Fig. S2-S10), there were a large number of herbivore-specific losses, in addition to a few gains (summarized in Figure 3; Table S7 for gene counts). The majority of copy number changes shared by all herbivores were concentrated among chemosensory gene families (22 out of 27). Herbivores lost several gustatory receptors that are required for bitter reception or are expressed in bitter gustatory neurons in *D. melanogaster* (multiple paralogs of isoform A of *Gr39a* [*Gr39aA*], *Gr59a, Gr59d*) (Kwon *et al*. 2011; Dweck and Carlson 2020). Herbivorous lineages also lost *Gr68a*, which is involved in the detection of an anti-aphrodisiac (Bray and Amrein 2003), and *Gr39aE* which has no known ortholog in *D. melanogaster*. The only OR lost was the aforementioned *Or22a*, which has been previously reported (Goldman-Huertas *et al*. 2015). Among the OBPs, almost all of the herbivore-specific losses (*Obp46a, Obp50cd, Obp58b, Obp58c, Obp58d, Obp93a*) were among the ‘plus-C OBPs,’ which are characterized as having more than six cysteine residues, and only two losses (*Obp18a* and *Obp56b*) were among the ‘classic OBPs’ with six cysteines (Fig. S12). Strikingly, non-herbivorous *Scaptomyza* have 11 Plus-C OBPs, while the herbivores possess only five. Almost all ionotropic receptors that were lost in the herbivores belonged to the ‘divergent IR’ class (*Ir7f, Ir51e, Ir56e, Ir94abc, Ir94f*), which are expressed in gustatory neurons. The only ‘antennal IR’ lost in all herbivores was *Ir60a,* and the only duplication was the divergent IR *Ir67a*. A single loss was found among PPKs (*ppk8*), whose function is unknown. Expression localization of *D. melanogaster* chemosensory gene orthologs that were lost or duplicated in all herbivores is presented in Fig. S13.

**Figure 3.**
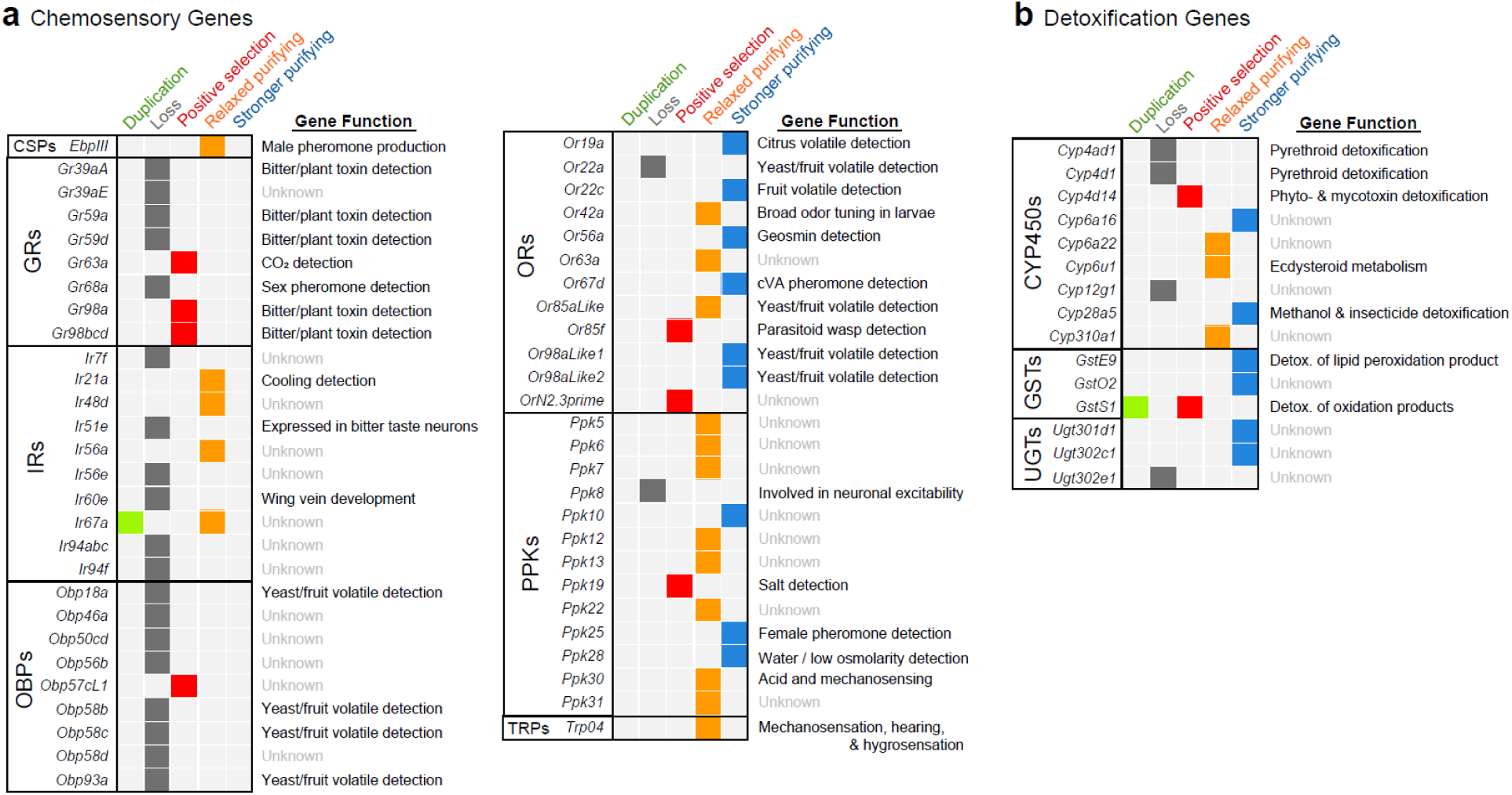
Genes involved in the detection of bitter compounds and yeast/fruit odorants disproportionately experienced gene loss or divergent selection in the ancestral *Scaptomyza* herbivore lineage. Summary of (a) chemosensory and (b) detoxification genes that were duplicated, lost or experienced divergent selection (positive selection or relaxed or stronger purifying selection). Gene duplications and losses are based on gene trees generated through RAXML, and genes that experienced changing selection pressures were identified with PAML. Gene counts from each species are shown in Table S7. Gene functions based on published data for *D. melanogaster* orthologs (sources for can be found in Table S10).

Only five herbivore-specific changes in copy number were found among the detoxification gene families: the duplication of *GstS1*, which has a role in oxidative stress responses and in flight muscle structure (Singh *et al*. 2001); the loss of *Ugt302e1*, whose function is currently unknown, and three cytochrome P450s (*Cyp4ad1, Cyp4d1,* and *Cyp12g1*). While there is no ortholog of *Cyp12g1* in *D. melanogaster*, both *Cyp4ad1* and *Cyp4d1* are upregulated in response to the pyrethroid deltamethrin (Liu *et al*. 2020).

### Genes under divergent selection in the herbivore lineage

We next used a ML approach through PAML to identify chemosensory and detoxification genes with signatures of changes in selection regime in the branch leading to the herbivorous species. We focused on this ancestral herbivore branch, rather than those leading to individual herbivore species, to identify changes that occurred as a result of or coincident with herbivory, rather than subsequent specialization on mustard or carnation plants. We used codon-based tests for positive selection, estimating dN/dS values under branch, branch-site, and clade models. Clade models offer more sensitivity in detecting divergent selection among a clade of genes, particularly those with multiple recent paralogs undergoing complex evolution (Weadick and Chang 2012). Thus, unsurprisingly, a large number of genes (128 genes) initially showed evidence of divergent selection through the use of clade models (*P* < 0.05; false discovery rate <5%). In these cases, we additionally tested branch and branch-site models on branches leading to different paralogs to identify which paralog experienced significant changing selection pressure. We report significant results from the initial branch and branch-site tests, and significant results from clade model follow-up tests. In total, 44 genes were found to have experienced a significant change in selection pressure along the basal herbivore branch (Tables 1 and S8). Results on all genes can be found in Supporting Dataset 3.

**Table 1.** Summary of selection analyses on chemosensory and detoxification genes. PAML analyses under branch and branch-site models. Branch model: M0 vs 2-ratios; branch model (positive selection): 2-ratios vs 2-ratios (ω1 = 1); branch-site model: Model A vs Model A (ω2 = 1). Omega values are reported only for the alternative model. Detailed results can be found in Table S8.

| Gene | Model | dN/dS |  | LRT<br>2(ΔlnL) | q-value<br>(FDR 5%) |
| --- | --- | --- | --- | --- | --- |
| Chemosensory Genes |  |  |  |  |  |
| Csp2 | branch | ω0=0.06 | ω1=0.24 | 7.14 | 0.05 |
| Gr63a | branch-site | ω0=0.03 (81%) | ω1=1 (7%) | 16.52 | <0.01 |
|  |  | ω2a=1 (11%) | ω2b=1 (1%) |  |  |
| (SmonGr98a1, SflaGr98a1,<br>SgraGr98a2) | branch | ω0=0.29 | ω1=0.87 | 7.74 | 0.04 |
|  | branch-site | ω0=0.22 (71%) | ω1=1 (26%) | 11.4 | 0.03 |
|  |  | ω2a=32.44 (2%) | ω2b=32.44 (1%) |  |  |
| Gr98bcd | branch-site | ω0=0.16 (73%) | ω1=1 (27%) | 10.2 | 0.05 |
|  |  | ω2a=179.55 (0.004%) | ω2b=179.55 (0.001%) |  |  |
| Ir21a | branch | ω0=0.11 | ω1=0.26 | 9.2 | 0.02 |
| Ir48d | branch | ω0=0.18 | ω1=0.38 | 7.96 | 0.04 |
| (Sflalr56a, Smonlr56a) | branch | ω0=0.33 | ω1=0.98 | 18.8 | <0.001 |
| (Sgralr56a1, Sgralr56a2, Sgralr56a4, Sgralr56a5) | branch | $\omega_0=0.33$ | $\omega_1=3.66$ | 7.98 | 0.03 |
| | branch (positive) | $\omega_0=0.34$ | $\omega_1=1$ | 1.56 | 0.31 |
| Ir67a | branch | $\omega_0=0.3$ | $\omega_1=0.6$ | 10.6 | 0.01 |
| Obp57cL1 | branch-site | $\omega_0=0.21$ (59%) | $\omega_1=1$ (36%) | 15.68 | <0.01 |
| | | $\omega_{2a}=148.51$ (3%) | $\omega_{2b}=148.51$ (2%) | | |
| Or19a | branch | $\omega_0=0.3$ | $\omega_1=0.09$ | 13.7 | <0.01 |
| Or22c | branch | $\omega_0=0.18$ | $\omega_1=0.05$ | 7.24 | 0.05 |
| (SflaOr42a1, SmonOr42a1, SgraOr42a1) | branch | $\omega_0=0.13$ | $\omega_1=0.45$ | 11.46 | <0.01 |
| Or56a | branch | $\omega_0=0.11$ | $\omega_1=0.03$ | 8.98 | 0.02 |
| Or63a | branch | $\omega_0=0.2$ | $\omega_1=0.76$ | 29.5 | <0.0001 |
| Or67d | branch | $\omega_0=0.18$ | $\omega_1=0.03$ | 15.18 | <0.01 |
| Or85aLike | branch | $\omega_0=0.13$ | $\omega_1=0.29$ | 6.98 | 0.05 |
| Or85f | branch-site | $\omega_0=0.13$ (88%) | $\omega_1=1$ (11%) | 12.54 | 0.03 |
| | | $\omega_{2a}=46.61$ (1%) | $\omega_{2b}=46.61$ (0.001%) | | |
| Or98aLike1 | branch | $\omega_0=0.19$ | $\omega_1=0.05$ | 7.58 | 0.04 |
| Or98aLike2 | branch | $\omega_0=0.2$ | $\omega_1=0$ | 35.92 | <0.0001 |
| OrN2.3prime | branch-site | $\omega_0=0.16$ (73%) | $\omega_1=1$ (20%) | 15.6 | <0.01 |
| | | $\omega_{2a}=15.3$ (5%) | $\omega_{2b}=15.3$ (1%) | | |
| Ppk5 | branch | $\omega_0=0.11$ | $\omega_1=0.36$ | 15.16 | <0.01 |
| Ppk6 | branch | $\omega_0=0.09$ | $\omega_1=0.25$ | 10.35 | 0.01 |
| Ppk7 | branch | $\omega_0=0.15$ | $\omega_1=0.27$ | 7.75 | 0.04 |
| Ppk10 | branch | $\omega_0=0.1$ | $\omega_1=0.01$ | 12.12 | <0.01 |
| Ppk12 | branch | $\omega_0=0.18$ | $\omega_1=0.38$ | 10.07 | 0.02 |
| Ppk13 | branch | $\omega_0=0.05$ | $\omega_1=0.19$ | 16.12 | <0.01 |
| Ppk19 | branch-site | $\omega_0=0.11$ (75%) | $\omega_1=1$ (23%) | 13.98 | 0.01 |
| | | $\omega_{2a}=15.75$ (2%) | $\omega_{2b}=15.75$ (1%) | | |
| Ppk22 | branch | $\omega_0=0.16$ | $\omega_1=0.37$ | 7.7 | 0.04 |
| Ppk25 | branch | $\omega_0=0.21$ | $\omega_1=0.04$ | 16.82 | <0.01 |
| Ppk28 | branch | $\omega_0=0.1$ | $\omega_1=0.03$ | 11.72 | <0.01 |
| Ppk30 | branch | $\omega_0=0.2$ | $\omega_1=0.44$ | 12.2 | <0.01 |
| Ppk31 | branch | $\omega_0=0.12$ | $\omega_1=0.39$ | 17.23 | <0.01 |
| Trp04 | branch | $\omega_0=0.03$ | $\omega_1=0.09$ | 10.66 | 0.01 |
| **Detoxification Genes** |  |  |  |  |  |
| Cyp28a5 | branch | $\omega_0=0.13$ | $\omega_1=0.05$ | 9.95 | 0.02 |
| Cyp310a1 | branch | $\omega_0=0.19$ | $\omega_1=0.57$ | 18.43 | <0.001 |
| Cyp4d14 | branch-site | $\omega_0=0.06$ (84%) | $\omega_1=1$ (15%) | 11.44 | 0.03 |
| | | $\omega_{2a}=809.22$ (1%) | $\omega_{2b}=809.22$ (0.001%) | | |
| Cyp6a16 | branch | $\omega_0=0.21$ | $\omega_1=0.1$ | 7.57 | 0.04 |
| Cyp6a22 | branch | $\omega_0=0.07$ | $\omega_1=0.26$ | 21.17 | <0.001 |
| Cyp6u1 | branch | $\omega_0=0.14$ | $\omega_1=0.61$ | 20.06 | <0.001 |
| GstE9 | branch | $\omega_0=0.15$ | $\omega_1=0.01$ | 9.35 | 0.02 |
| GstO2 | branch | $\omega_0=0.12$ | $\omega_1=0$ | 10.3 | 0.01 |
| (SflaGstS1b, SmonGstS1b, SgraGstS1b) | branch | $\omega_0=0.09$ | $\omega_1=3.27$ | 50.43 | <0.0001 |
| | branch-site | $\omega_0=0.08$ (84%) | $\omega_1=1$ (2%) | 14.04 | 0.01 |
| | | $\omega_{2a}=38.47$ (13%) | $\omega_{2b}=38.47$ (0.004%) | | |
| Ugt301D1 | branch | $\omega_0=0.09$ | $\omega_1=0.03$ | 7.22 | 0.05 |
| Ugt302C1 | branch | $\omega_0=0.11$ | $\omega_1=0.04$ | 7.42 | 0.05 |

Nine genes were identified as having experienced positive selection (dN/dS >1) in the basal herbivore lineage: *Gr63a, Gr98a, Gr98bcd, GstS1b, Obp57cL1, Or85f, OrN2.3prime, Cyp4d14*, and *Ppk19* (Table 1). The proportion of positively selected sites among these genes was generally low, with a few exceptions. One such exception was *GstS1b*, which exhibited 13% positively selected sites (dN/dS = 38.47), almost all of which had exceptionally strong support (i.e., high posterior probabilities, *P*>0.95). As noted earlier, *GstS1* was duplicated in the ancestor of herbivorous *Scaptomyza*. The single ortholog in *D. melanogaster* is involved in oxidative stress response, conjugation of the lipid peroxidation product 4-HNE, and modulating methylmercury toxicity (Singh *et al*. 2001; Whitworth *et al*. 2005; Vorojeikina *et al*. 2017). Despite expression of both *GstS1a* and *GstS1b* in the *S. flava* larval gut (Gloss *et al*. 2019), positive selection on only *GstS1b* indicates potential neofunctionalization of this copy. Notably, mustard-derived toxins (isothiocyanates) induce the formation of peroxidized lipids (Gago-Dominguez *et al*. 2007).

Two genes (*Gr98a* and *Gr98bcd*) that experienced positive selection along the ancestral herbivore lineage are involved in the detection of noxious, ‘bitter’ compounds. The *D. melanogaster* ortholog of *Gr98a* is involved in detecting histamine, which is high in fermented foods (Aryal & Lee 2022) and *Gr98b* detects L-canavanine, an extremely bitter plant-derived toxic amino acid (Shim *et al*. 2015).

Among genes that did not show a strong indication of positive selection but were found to be evolving at significantly different rates between herbivores and background branches, many (20/35) had higher foreground dN/dS values (Table 1), indicating relaxation from selective constraints among the herbivores, whereas the remaining experienced stronger purifying selection. Many ORs in *D. melanogaster* have been de-orphanized, providing a wealth of information about the breadth and sensitivity of individual ORs (e.g. (Hallem *et al*. 2004)). Though the properties of *D. melanogaster*’s ORs cannot be used to precisely determine those of other species, it offers a starting point to generate hypotheses about how ORs are evolving in closely related species. We found that almost all ORs experiencing relaxed purifying selection (*Or42a1, Or85aLike*) or stronger purifying selection (*Or98aLike1*, *Or98aLike2*) detect esters and alcohols, which are produced in high abundance by yeasts and fruit, and are attractive to microbe-feeders (Becher *et al*. 2012), but not herbivores (Goldman-Huertas *et al*. 2015; Matsunaga *et al*. 2022).

Detoxification genes that experienced relaxed purifying selection in the ancestral herbivore lineage were *Cyp310a, Cyp6a22,* and *Cyp6u1*, and those that experienced stronger purifying selection were *GstE9, GstO2, Cyp28a5, Cyp6a16, Ugt301d1,* and *Ugt302c1* (Table 1). The majority of these have been found to be upregulated in response to toxin consumption, although some have putative functions in development, hormone metabolism, cold tolerance, and olfaction (Fig. 3).

### Effects of demographic history

Demographic processes, such as population bottlenecks, can weaken the efficacy of natural selection, leading to an accelerated fixation of slightly deleterious gene gains and losses (Gardiner *et al*. 2008). To investigate the possibility that our results on gene family evolution were confounded by demographic events, we examined three lines of evidence from our analyses.

First, the randomly selected set of orthologous clusters (∼200 genes) exhibited gain, loss, and turnover rates that were not significantly higher among herbivores than in other *Drosophila* (λ_herb_=0.002; λ_non-herb_=0.003; q-value<0.02, Fig. 2). These results suggest demographic processes were not the underlying cause of elevated rates of turnover among chemosensory and detoxification genes because these demographic processes would have generated similar patterns genome-wide.

Second, we inferred the level of nucleotide diversity (π) in a *S. flava* population (collected in 2013 from Belmont, MA, USA), using pooled whole genome sequencing (methods described in the Supplemental Methods). Autosomal nucleotide diversity, which is proportional to the coalescent effective population size, was similar between a single population of *S. flava* (π = 0.0056) and North American populations of *D. melanogaster* (the DGRP; π = 0.0060 (Mackay *et al*. 2012)). The relationship between physical distance and linkage disequilibrium was also similar between the two species and decayed quickly (Peláez *et al*. 2022), consistent with sharing similarly large coalescent effective population sizes (Sved 1971).

Third, we estimated the proportion of the genome composed of repetitive elements, which is sensitive to demographic shifts (Bourgeois and Boissinot 2019), and found that repeat content in *S. flava* is within the range observed across *Drosophila* species (Table S9).

## Discussion

In this study, we sought to determine the extent to which the evolution of herbivory affected patterns of molecular evolution across chemosensory and detoxification gene families in the Drosophilidae. We focused on a clade within the genus *Scaptomyza* – nested within the paraphyletic genus *Drosophila* – that evolved to feed as both leafminers (larvae) and leaf puncture-feeding adults (females) on living plants <15 mya (Fig. 1a). A great deal is known regarding how these gene families, especially chemosensory ones, evolve across *Drosophila* species in the context of diet shifts and diet specialization (McBride 2007; Gardiner *et al*. 2008; Rane *et al*. 2019; Reisenman *et al*. 2023). However, we have far less insight into how these gene families evolve in response to a truly herbivorous niche shift (i.e., larval development is completed by feeding on living plants).

Here, we identified, through iterative curation and manual inspection, the full complement of several chemosensory and detoxification gene families for five *Scaptomyza* species – three herbivorous species within the subgenus *Scaptomyza* (*S. flava*, *S. montana,* and *S. graminum*) and two non-herbivorous species from two other subgenera (*S. pallida* in *Parascaptomyza* and *S. hsui* in *Hemiscaptomyza*) – and more distantly related *Drosophila* species. By including species across a range of diets and phylogenetic distances, we were able to disentangle herbivore-specific changes from those shared *Scaptomyza*-wide and those involved in more recent host plant-specific specialization (*S. flava* and *S. montana* on Brassicales and *S. graminum* on Caryophyllaceae), although other selective and neutral processes may have effects.

Despite consistent reductions in gene family sizes across all three herbivorous species (Fig. 2b, Table S3), we did not find significant gene turnover, duplications or losses for most chemosensory and detoxification gene families along the ancestral herbivore branch (Fig. 2c-e), contrary to the long-held view that the transition to herbivory involves extensive expansions and contractions of these gene families (McBride 2007; Edger *et al*. 2015; Johnson *et al*. 2018). The only exceptions were GRs and OBPs that were lost at a significantly higher rate (Fig. 2e), which is consistent with the neural limitation hypothesis (Bernays 2001). The loss of chemosensory genes, by limiting sensory inputs, facilitates rapid and accurate decision-making in herbivores that are faced with many host plant options. The lack of excessive copy number changes among the remaining gene families suggests that the initial transition to herbivory requires a more limited set of changes and subsequent high turnover may be related to further specialization. While our phylogenetic analyses could not elucidate precisely whether the identified genetic changes along the ancestral branch occurred before, during or after herbivory evolved, several of the candidate genes have been implicated in other dietary transitions (discussed below), providing strong support for their involvement in the evolution of herbivory as well.

### Evolutionary patterns across gustatory receptors involved in detecting bitter compounds

Numerous studies have now shown that expansions of bitter GRs are typical of generalist species, to enable detection of a wide variety of plant-derived bitter compounds, while oligophagous and monophagous herbivores derived from generalist ancestors tend to lose some of these bitter GRs. This pattern has been found in butterflies, moths, aphids, flies, beetles, sawflies, and mites (Smadja *et al*. 2009; Xu *et al*. 2016; Suzuki *et al*. 2018; Crava *et al*. 2020; Vertacnik *et al*. 2021). Our results suggest that the ancestral herbivore lineage experienced a loss in bitter detection as a result of losing many bitter GRs (paralogs of *Gr39aA*, *Gr59a, Gr59d*) and experiencing rapid evolution of other bitter GRs (*Gr98a* and *Gr98bcd*). These evolutionary genetic changes likely reduced the ability of these flies to detect bitter compounds, which would otherwise limit their intake of their host plants. A reduction in bitter sensitivity has been found in *D. suzukii* (Dweck *et al*. 2021), which has evolved herbivory (feeding on living, ripe fruit) on a similar timescale as herbivorous *Scaptomyza* species (Suvorov *et al*. 2022). However, although *D. suzukii* is herbivorous (attacks ripe fruit), it is polyphagous on many plant families (Poyet *et al*. 2015), suggesting that the loss of bitter reception is a trait involved in the transition to herbivory, independent of specialization.

With the exception of *Gr39aA*, these candidate bitter GRs do not encode “commonly expressed receptors” – a set of bitter GRs expressed in all bitter gustatory neurons (Weiss *et al*. 2011; Dweck and Carlson 2020). This suggests that, despite a diet containing bitter, toxic compounds, *Scaptomyza* species are still able to detect bitter compounds generally, which is also true of specialist feeders on toxic hosts: *D. sechellia*, *D. erecta,* and *D. suzukii* (Dweck and Carlson 2020; Dweck *et al*. 2021). This ability is likely important for being able to differentiate toxin levels between individual leaves or host plants and distinguishing old from young or healthy from damaged plants. In *S. flava*, feeding on plants bearing aliphatic and indolic glucosinolates causes them to develop more slowly (Gloss *et al*. 2019), so differentiating between different types of bitter host-derived compounds could impact their fitness.

The lost copies of *Gr39aA, Gr59a*, and *Gr59d* are particularly interesting candidates to focus on for future study in relation to the evolution of toxin specialization, considering all three GRs have been implicated in dietary shifts in other species. *Gr59a* and *Gr59d* both underwent expansions in *D. suzukii* (Hickner *et al*. 2016), while both of these genes were lost in *D. sechellia* and *D. erecta* (McBride *et al*. 2007). *D. suzukii* lost some bitter sensitivity, but through transcriptional down-regulation of bitter GRs (Dweck *et al*. 2021). The pattern of expansion of *Gr59a* and *Gr59d* in a generalist (*D. suzukii*) and contraction in specialists (*D. sechellia*, and *D. erecta*) seems to strongly suggest involvement in mediating host breadth. Furthermore, signals of positive selection have been detected in *Gr59a* in *D. yakuba mayottensis,* which has convergently evolved specialization on noni fruit along with *D. sechellia* (Ferreira *et al*. 2020). Similarly, losses of *Gr39aA* have occurred across independently evolved specialist lineages in both *D. sechellia* and *D. erecta* (McBride *et al*. 2007). *Gr39a* encodes several different isoforms, but only *Gr39aA* is expressed in all bitter gustatory neurons, and is involved in the detection of many bitter compounds in *D. melanogaster* (Dweck and Carlson 2020). Because *D. melanogaster* only bears a single copy of the A exon, it remains to be studied whether these additional *Gr39aA* copies are expressed in species bearing multiple copies. Notably, all three of these GRs exhibit numerous tandem duplications. The remarkably high rate of turnover for *Gr59d* and *Gr39aA* across the sampled species (Table S6) suggests that there may be transposable elements in the vicinity driving these duplications, which could be selected upon during dietary shifts to generate a dosage effect, where more copies may enable stronger bitter detection. Altogether, our results on bitter GR evolution indicate a strong likelihood for reduced bitter detection in the ancestral herbivorous *Scaptomyza* species.

### Concerted losses of and positive selection on genes involved in olfaction

Although we did not find excessively high levels of OR gene turnover at the base of the *Scaptomyza* herbivore clade, the ORs that were lost shared related and ecologically important functions, specifically in detecting yeast and fruit volatiles. Goldman-Huertas et al. (2015) showed that the evolution of herbivory in *S. flava* was associated with OR gene losses that reduced attraction towards their ancestral diet of yeast feeding. This was mediated through a behavioral loss of attraction to yeast volatiles, putatively ancestral feeding attractants, and with the step-wise loss of olfactory receptors (*Or22a*, *Or42b*, and *Or85d*) tuned to detect them (Goldman-Huertas *et al*. 2015). With full annotations from four other *Scaptomyza* species, we obtained results consistent with this finding, with the addition of two other yeast volatile-detecting ORs (*Or42a* and *Or85aLike*) that experienced relaxed purifying selection (Table 1, Fig. S5). In keeping with past studies on *Or22a* in various species, the loss of *Or22a* – which was the only OR loss shared exclusively by all herbivores in this study – strongly indicates an outsized role for *Or22a* in association with dietary shifts. *Or22a* and *Or22b* (sometimes found as a single copy in some lineages of *Drosophila*) have been subjected to repeated, independent bouts of natural selection, resulting in: increased sensitivity in *D. sechellia* to esters specific to noni fruit (*Morinda citrifolia*) (Dekker *et al*. 2006); increased sensitivity to *Pandanus* fruit volatiles in *D. erecta* (Linz *et al*. 2013); attraction to marula fruit (*Sclerocarya birrea*) in some populations of *D. melanogaster*; and finally, loss of sensitivity towards fermentation odors and increased sensitivity to certain leaf volatiles in the ripe fruit specialist *D. suzukii* (Keesey *et al*. 2015). Collectively, this indicates the importance of *Or22a* for driving attraction to fermenting plant tissues in drosophilids, which is no longer a niche used by ovipositing females of herbivorous species.

Previous divergence-based genomic analyses that included only *S. flava* and the 12 original *Drosophila* species with genome annotations found evidence of positive selection on *Or63a, Or67b* paralogs, *Or88a*, and *Or98a* (Goldman-Huertas *et al*. 2015), whereas here, we only found evidence of positive selection on *Or85f* and *OrN2.3prime* along the ancestral herbivore branch. This highlights the advantage of including additional closely related herbivorous species and non-herbivorous *Scaptomyza* species (i.e., *S. pallida* is ∼23 million years diverged from the herbivores versus *D. grimshawi* at *∼*33 million years (Matsunaga *et al*. 2022)). In particular, there is now strong evidence that the triplication and strong positive selection on *Or67b* paralogs is specifically related to *S. flava*’s specialization on plants of the order Brassicales, as these ORs are specifically tuned to the volatile isothiocyanates (Matsunaga *et al*. 2022). Similarly, we found that previous analyses that focused on *S. flava* without the inclusion of close *Scaptomyza* species identified the rapid evolution and repeated duplication of *GstE5-8* (5 copies) as significant for the evolution of herbivory in this lineage (Gloss *et al*. 2019), but our analyses reveal that this expansion is restricted to the mustard specialists.

The most striking herbivore-specific loss of genes occurred within the OBPs. OBPs have been classified into three groups: classic OBPs that have six cysteine residues, minus-C with less than six, and plus-C with more than six (Zhou *et al*. 2006). The majority of OBPs genes lost in herbivores (six out of eight) were among the Plus-C class (Fig. S12). Some of the plus-C OBPs lost in herbivores (*Obp58b, Obp58c,* and *Or85a*) are expressed only in the antennae and/or head of *D. melanogaster*, but other lost OBPs have been found additionally in the legs or body of adults (*Obp46a, Obp50c, Obp93a*) (Zhou *et al*. 2004; Larter *et al*. 2016). While it is unclear if any of these OBPs play a role in olfactory perception and what the functional significance of bearing additional cysteine residues is, we speculate that the Plus-C OBPs may be more vulnerable to damage by plant toxins, like the largely Brassicales-specific isothiocyanates, which are highly electrophilic and attack nucleophilic sulfhydryl moieties of the cysteine residues (Kawakishi and Kaneko 1987).

### Diverse functions of sensory genes associated with herbivory

In this study, we hypothesized that the majority of genetic changes associated with the evolution of herbivory would involve genes that interact with host plant compounds. It was thus surprising to find significant changes in genes involved in detecting various other stimuli: carbon dioxide (positive selection on *Gr63a*), salinity (positive selection on *Ppk19*), water/low osmolarity (stronger purifying selection on *Ppk28*), cooling (relaxed purifying selection on *Ir21a*), and pheromones (loss of *Gr68a,* strong purifying selection on *Ppk25*) (Liu *et al*. 2003; Bray and Amrein 2003; Kwon *et al*. 2007; Cameron *et al*. 2010; Ni *et al*. 2016). This suggests that while the toxins in the diet play a significant role in driving the evolution of these gene families, the leaf-mining lifestyle imposes other sensory changes. The larvae of herbivorous species navigate through the mesophyll of the leaf – a fluid-filled cavity, where salt levels, temperature gradients, and risk of carbon dioxide poisoning may be significantly different than within decaying organic material.

### Duplication and positive selection on a detoxification gene

Numerous studies have indicated that gene duplications, followed by functional divergence, can spur biological novelties – new traits or adaptations to new niches (reviewed in (Carscadden *et al*.) 2022). Examples abound across the diversity of life and across complex traits, from trichromatic vision in old world monkeys to snake venom phospholipase genes (Dulai *et al*. 1999; Lynch 2007). Contrary to our initial hypotheses and predictions that we might find numerous instances of gene duplications across chemosensory and detoxification gene families, our phylogenetic gene trees of each family revealed that duplications were more prevalent within individual herbivore species or among just the mustard specialists, with only two gene duplications shared by the three surveyed herbivores.

This suggests that gene duplications may spur further specialization but may not play a prominent role in initial transitions to herbivory. We cannot speculate on the role of the *Ir67a* duplication because the function is unknown in other species. However, the duplication and rapid evolution of *GstS1b* (Table 3) may be indicative of its involvement in adaptation to herbivory, as its *D. melanogaster* ortholog encodes an enzyme involved in detoxifying oxidation products, lipid peroxidation products, and organometallic compounds (Singh *et al*. 2001; Whitworth *et al*. 2005; Saisawang *et al*. 2012; Vorojeikina *et al*. 2017). Both *GstS1* copies are expressed in *S. flava* larvae, but there is not strong up-regulation of either copy in response to isothiocyanates, the defense compound in mustard plants, reinforcing that this duplication is not specific to mustard feeding. In *D. melanogaster*, *GstS1* has high expression in flight muscle and in the adult central nervous system, with previous studies suggesting both structural and detoxifying functions (Clayton *et al*. 1998; Whitworth *et al*. 2005). Functional genetic testing in herbivorous drosophilids should reveal the fate and possible functional divergence of these two paralogs: whether expression patterns differ across tissues, whether the loss of either or both paralogs affects flight muscle structure, and the specificity of each paralog towards diet-derived xenobiotics.

Finally, the facultatively herbivorous *S. flavella*, endemic to New Zealand and in a different subgenus than the obligate herbivores, lays eggs on rotting leaves of sea celery (*Apium prostratum*) – the females lack a cutting ovipositor (Martin 2014). Their neonate larvae cannot mine the leaves of living celery plants, but the second instars move into these living leaves and form gregarious mines similar to those formed by the larvae of *S. flava*. Thus, a facultative, larva-first transition from microbe-feeding/saprophagy rather than an adult-first transition (cutting ovipositor-enabled) seems likely. This suggests that phenotypic plasticity and facultatively herbivorous larvae may pave the way to herbivory and host plant specialization in drosophilids and that exaptation, rather than extensive genome evolution, during the first stages of this dietary shift may be more salient.

## Conclusions

Genomic comparisons of older herbivorous lineages to distantly related non-herbivores, or across herbivorous lineages, have uncovered striking expansions and losses of genes involved in chemosensation and detoxification in arthropods (e.g., (Goldman-Huertas *et al*. 2015; Rane *et al*. 2016; Calla *et al*. 2017; Schoville *et al*. 2018; Johnson *et al*. 2018). Yet, the lack of dense sampling among closely related taxa has, in many cases, precluded pinpointing of the timing of these changes relative to the evolution of herbivory. Here, using a comparative genomic approach across a growing number of drosophilid genome assemblies, we found that the evolution of herbivory involved some accelerated protein evolution and copy number changes across detoxification and chemosensory gene families. These patterns of evolution lend support to the hypothesis that the chemical composition of plant tissues drives herbivore genome evolution, an idea at the core of early theories on species interactions that motivated the development of co-evolutionary theory (Fraenkel 1959; Ehrlich and Raven 1964; Swain 1977; Berenbaum and Zangerl 2008). Similar comparative approaches in other young herbivore lineages may reveal the extent to which the genomic changes tied to herbivory in *Scaptomyza* species reflect general strategies underpinning the evolution of herbivory.

## Data Availability

The *S. flava* Illumina-only genome assembly (sfla_v1) is available as GenBank assembly accession GCA_003952975.1, and the PacBio/HiC/Illumina assembly (sfla_v2) will be made available at the time of publication. Source data, scripts, and analysis output files are accessible as Supporting Datasets 1-4 in the Dryad repository (doi:10.6078/D1841H); for further details, see “List of Supporting Datasets” in the Supplementary Materials.

## Acknowledgements

We wish to acknowledge Kamalakar Chatla and Dovetail Genomics for assistance in assembling the *Scaptomyza flava* genome (sfla_v2).

