## Supplemental Methods & Figures for "Evolution of chemosensory and detoxification gene families across herbivorous Drosophilidae"

**Supplementary Material for**
**“Evolution of chemosensory and detoxification gene families across herbivorous Drosophilidae”**

### 1. SUPPLEMENTARY METHODS

***S. flava* genome sequencing and assembly (sfla_v2).**

Dovetail HiC library preparation and sequencing. *S. flava* flies used for the genome assembly were collected from a partially inbred laboratory colony. The colony was founded from >150 larvae collected near Dover, NH, USA, and subsequently maintained for several years in the laboratory. 300 male flies were flash frozen in liquid nitrogen and stored at -80°C. A Dovetail HiC library was prepared in a similar manner as described previously (Lieberman-Aiden *et al.* 2009). Briefly, for each library, chromatin was fixed in place with formaldehyde in the nucleus and then extracted. Fixed chromatin was digested with DpnII, the 5’ overhangs filled in with biotinylated nucleotides, and free blunt ends were ligated. After ligation, crosslinks were reversed and the DNA purified from protein. Purified DNA was treated to remove biotin that was not internal to ligated fragments. DNA was then sheared to ~350 bp mean fragment size, and sequencing libraries were generated using NEBNext Ultra enzymes and Illumina-compatible adapters. Biotin-containing fragments were isolated using streptavidin beads before PCR enrichment of each library. The libraries were sequenced on an Illumina HiSeqX to produce 380 million 2x150 bp paired end reads, to approximately 30x sequence coverage.

PacBio library and sequencing. DNA samples were quantified using Qubit 2.0 Fluorometer (Life Technologies, Carlsbad, CA, USA). SMRTbell libraries (~20kb) for PacBio Sequel were constructed using SMRTbell Template Prep Kit 1.0 (PacBio, Menlo Park, CA, USA) using the manufacturer's recommended protocol. The pooled library was bound to polymerase using the Sequel Binding Kit 2.0 (PacBio) and loaded onto PacBio Sequel using the MagBead Kit V2 (PacBio). Sequencing was performed on two PacBio Sequel SMRT cells, using Instrument Control Software Version 5.0.0.6235, Primary analysis software Version 5.0.0.6236 and SMRT Link Version 5.0.0.6792.

Illumina library and sequencing. Illumina sequence data was generated from the “OHI-9 line” of *S. flava*, derived from a laboratory population initially collected in Belmont, MA, USA in 2008 that was inbred through 10 generations of single pair sibling mating on *A. thaliana* Col-0 plants. Paired-end 180 bp and 300 bp insert libraries and 3 kbp and 5 kbp mate pair libraries from OHI-9 female flies were sequenced with 100 bp read length on an Illumina Hiseq 2000 at the University of Arizona. Reads were quality filtered and Illumina TruSeq3 adapters were removed using Trimmomatic v0.35 (Bolger *et al.* 2014) with the following parameters: “LEADING:10 TRAILING:10 SLIDINGWINDOW:4:15 MINLEN:99”.

Falcon and HiRise initial assembly. An initial draft assembly of sfla_v2 was generated by Dovetail using the following methods, but was not used because of a high rate of errors detected and an inflated genome size, which we suspected was caused by high heterozygosity in the assembly. We report these methods, however, because this assembly was used for scaffolding our final assembly (details below). The genome assembly was performed using the FALCON 1.8.8 pipeline from Pacific Bioscience. First, 70-fold whole-genome, single-molecule, real-time sequencing (SMRT) data of *S. flava* was used as input to the traditional FALCON pipeline using a length cut-off that correspond to 50x coverage of data during the initial error-correcting stage. This resulted in 0.511 million error corrected reads with an N50 read length equal to 19.6 kb. Second, the error-corrected reads were processed by the overlap portion of the FALCON pipeline. The aligned reads were assembled in the third stage of FALCON into 3,561 primary contigs containing 443.7 Mbp with an NG50 contig length of 543.7 kbp. Finally, the assembly was polished through PacBio's Arrow algorithm from SMRT Link 5.0.1, using the original raw-reads. The input de novo assembly, and Dovetail HiC library reads were used as input data for HiRise, a software pipeline designed specifically for using proximity ligation data to scaffold genome assemblies (Putnam *et al.* 2016). Dovetail HiC library sequences were aligned to the draft input assembly using bwa (http://github.com/lh3/bwa). The separations of Dovetail HiC read pairs mapped within draft scaffolds were analyzed by HiRise to produce a likelihood model for genomic distance between read pairs, and the model was used to identify and break putative misjoins, to score prospective joins, and make joins above a threshold.

Reassembly. As mentioned above, while this initial de novo assembly exhibited high scaffold contiguity, we discovered that it unfortunately contained a significant number of frameshift mutations in coding sequences that limited our ability to correctly annotate the genome. A BUSCO (v5.2.2, (Manni *et al.* 2021)) analysis only showed 83.4% complete BUSCOs, notably lower than typical long-read drosophilid genome assemblies (e.g., Kim et al. 2021). A manual examination of preliminary annotation records showed that approximately 15% of randomly checked genes contained one or more deletions resulting in fragmented exons. We attempted to further polish the scaffolds with Illumina data but that resulted in only 97.4% complete BUSCOs and many of the manually checked gene annotation records contained indels.

We thus decided to reassemble the genome, using the same data described here, with the long-read hybrid assembly pipeline described in Kim et. al (2021), which has been shown to produce highly complete genome assemblies for drosophilid flies with very few coding sequence indels (Kim *et al.* 2021). Briefly, we generated an initial draft assembly with Flye 2.9 (Kolmogorov *et al.* 2019), identified and removed duplicated haplotypes (haplotigs) with purge_dups (Guan *et al.* 2020), polished the draft assembly using the PacBio reads with one round of Racon (Vaser *et al.* 2017), then polished further using Illumina reads with one round of Pilon (Walker *et al.* 2014), only fixing base-level errors. The fully polished assembly was scanned for contaminant sequences using NCBI BLAST (Johnson *et al.* 2008) and BlobTools (Laetsch and Blaxter). Repetitive sequences in the assembly were identified with RepeatModeler2 (Flynn *et al.* 2020). This produced a fairly contiguous 323.9 Mbp assembly contained in 1,430 contigs, with an N50 of 575,920 bp and L50 of 148. BUSCO completeness was significantly improved (98.8%) and manual spot checks of another preliminary annotation showed no indel issues.

Scaffolding. To fully integrate the long-range assembly information we had previously generated with HiC, we scaffolded the less contiguous but more accurate new assembly with the error-prone Dovetail version, reasoning that base-level errors in the previous version were unlikely to impact reference-based scaffolding. Specifically, we soft-masked both genomes using the repeat library generated in the previous step, using RepeatMasker (Smit *et al.* 2013). Then, we created a whole-genome alignment with Progressive Cactus (Armstrong *et al.* 2020) and used the RagOut reference-based scaffolder (Kolmogorov *et al.* 2014) to scaffold the new genome, using the old assembly as the reference. The final genome assembly of *S. flava* (sfla_v2) was scaffolded into 1,252 scaffolds covering 315.4 Mbp (N50 = 32.966 Mb) with a maximum length of 85.98 Mbp and with 460 gaps.

Comparative annotation of *Scaptomyza* genomes. Gene annotations previously created for an Illumina-only *S. flava* assembly (sfla_v1) (described below) were transferred to the assemblies of *S. graminum, S. hsui, S. pallida, S. montana*, and this newer *S. flava* assembly, using whole-genome Progressive Cactus alignments and the Comparative Annotation Toolkit (CAT, (Fiddes *et al.* 2018)). A *D. grimshawi* genome (Kim *et al.* 2021) was used as an outgroup. All genomes were repeat-masked using repeat libraries generated with RepeatModeler2 and soft-masked using RepeatMasker. A published phylogeny for these species (Suvorov *et al.* 2022) was used as a guide tree for the alignment. RNA-seq data generated from female *S. flava* gut, proboscis, and maxillary palp organs was downsampled to 100X coverage using BBTools (Bushnell 2014) and aligned to the *S. flava* genome using STAR (Dobin *et al.* 2013) to assist CAT annotation. RNA-seq data are currently not available for any of the other species. Briefly, CAT uses the transMap mode to project annotations from a reference (i.e, the original *S. flava* genome) onto the target genomes, evaluating the projections and correcting them with Augustus (Stanke *et al.* 2008) and utilizing RNA-seq evidence if it exists.

##### *S. flava* Illumina-only assembly (sfla_v1).

Prior to the generation of our main assembly sfla_v2, an *S. flava* assembly generated from Illumina sequencing was utilized by various projects on this species (e.g. (Peláez *et al.* 2020), (Gloss *et al.* 2022), Verster et al. 2019). The sequence data was also used in polishing steps in the long read assembly, for generating automated annotations that were then also copied over to the newer assembly, sfla_v2, and for estimating repeat coverage.

Assembly and annotation. Read pairs that survived quality filtering were subsampled to an estimated ~90x coverage of the *S. flava* genome and assembled using ALLPATHS-LG (Gnerre *et al.* 2011) on the XSEDE high performance computing system. Contigs were extended and ambiguous regions were resolved iteratively using GapCloser (Luo *et al.* 2012). Prior to annotation, repeat regions were masked using RepeatMasker (Smit *et al.* 2010) with the *Drosophila* repeat library. Protein-coding genes were annotated using MAKER2 (Holt and Yandell 2011), with the *S. flava* transcriptome (Whiteman *et al.* 2012) and predicted gene sequences from 12 *Drosophila* species (FlyBase release 2013_06) provided to inform gene models. We recovered 17,997 genes in our liberal gene set (annotations predicted by Augustus). The proportion of core dipteran genes recovered in our assembly was determined with BUSCO v5.4.2 (Simão *et al.* 2015).

Repetitive element content. Repeat identification was carried out using both homology-based and ab-initio approaches. We used the Drosophila RepBase repeat database for the homology-based annotation (http://www.girinst.org/repbase; update 20150807) within RepeatMasker version open-4.0.6 (Smit *et al.* 2010). The RepeatMasker option -gccalc was used to infer GC content for each contig separately to improve the repeat annotation. Ab-initio repeat finding was carried out using RepeatModeler version 1.73 (http://repeatmasker.org/RepeatModeler.html).

**Gene model validation.**

To validate lost genes of interest (GOIs), we performed an additional TBLASTN search for each GOI in *S. flava*, as well as genes assumed to be proximal to GOIs (PGOIs) based on their location in *D. grimshawi* or *D. virilis*. These searches used the predicted orthologs in *D. grimshawi* or *D. virilis* as queries. GOIs were considered truly absent if TBLASTN searches yielded no hits. In some cases, homology between conserved protein domains resulted in weakly supported hits to the *S. flava* genome and/or transcriptome, in which case the aligned region was extracted, translated, and BLASTed against the NCBI nr database. The gene was considered lost if the output showed stronger homology to genes other than the GOI. To avoid confirmation bias, we also did this with the *S. flava* PGOIs to ensure their identity matched the expected ortholog in *D. grimshawi*. The absence of the GOIs, coupled with the presence of 95% of the PGOIs strongly support that the GOIs are truly lost and are not an artifact of missing scaffolds in the genome assembly. We implemented a similar BLAST-based search of both genomes to validate genes that underwent lineage-specific expansions in *S. flava*.

Finally, to guard against errors in our curation of the published *Drosophila* genomes, we compared our gene curations to those from published studies (Low *et al.* 2007; Almeida *et al.* 2014; Good *et al.* 2014) and those inferred in OrthoDB (Zdobnov *et al.* 2017). We performed comprehensive TBLASTN searches against the relevant genome assemblies to search for the full complement of orthologous genes, re-curated gene models if necessary, and manually inspected aligned gene models. In a few cases, we discarded genes that had high similarity (>99% nucleotide identity) and perfect synteny to another scaffold in the assembly because these are likely artifactual duplicates.

**Population genomics.**

DNA pooled from 45 wild-collected *S. flava* larvae was sequenced to yield 100 bp paired-end reads on an Illumina HiSeq 2000. Methods on sample collection, sequencing, and read mapping can be found in Pelaez, Gloss et al. 2022. Nucleotide diversity (π) was calculated across four-fold degenerate sites using the script Variance-sliding.pl from Popoolation v.1.2.2 (Kofler *et al.* 2011) for repeat-masked autosomal scaffolds (N = 819) greater than 20 kb in length. Parameters were: a ploidy level of 90, minimum allele count of two, minimum quality score of 20, minimum coverage of four, and a maximum coverage of 100. Scaffolds were considered autosomal if more than half the predicted proteins had a best BLASTP score to an autosome in *D. melanogaster*.

#### 2. SUPPLEMENTARY TABLES

##### Table S1. Species included in analyses and their assembly versions.

| Species | Assembly version | CAFE | PAML |
|---|---|---|---|
| *Scaptomyza flava* | sfla_v1 | | |
| *Scaptomyza flava* | sfla_v2 | ✓ | ✓ |
| *Scaptomyza montana* | iso-CA-L1 | ✓ | ✓ |
| *Scaptomyza graminum* | TMU-2019 | ✓ | ✓ |
| *Scaptomyza pallida* | iso-CA-L1 | ✓ | ✓ |
| *Scaptomyza hsui* | iso-CA-L1 | ✓ | ✓ |
| *Drosophila grimshawi* | dgri_caf1 | ✓ | ✓ |
| *Drosophila mojavensis* | dmoj_caf1 | ✓ | ✓ |
| *Drosophila virilis* | dvir_caf1 | ✓ | ✓ |
| *Drosophila melanogaster* | Release_6_plus_ISO1_MT | ✓ | ✓ |
| *Drosophila ananassae* | dana_caf1 | ✓ | |
| *Drosophila erecta* | dere_caf1 | ✓ | |
| *Drosophila pseudoobscura* | Pse_3.0 | ✓ | |

##### Table S2. *Scaptomyza* genome assembly statistics.

| Assembly | Sequencing | Number of scaffolds | Longest scaffold (Mbp) | Assembly length (Mbp) | N50 (Mbp) | L50 | Automated annotations |
|---|---|---|---|---|---|---|---|
| *S. flava* (sfla_v2) | PacBio + HiC + Illumina | 781 | 92.65 | 331.7 | 31.83 | 3 | 12,365 |
| *S. flava* (sfla_v1) | Illumina | 6619 | 2.85 | 216.1 | 0.112 | 404 | 17,997 |
| *S. montana* | Nanopore | 735 | 22.67 | 229.1 | 2.44 | 18 | 11,924 |
| *S. graminum* | Nanopore | 352 | 21.82 | 137.8 | 17.41 | 4 | 11,747 |
| *S. pallida* | Nanopore | 273 | 10.12 | 201.7 | 4.28 | 15 | 11,701 |
| *S. hsui* | Nanopore | 313 | 20.51 | 223.5 | 5.29 | 10 | 11,680 |

**Table S3. Chemosensory and detoxification gene family sizes across drosophilid genomes.** Gene coordinates for *Scaptomyza* species and gene ID numbers for *Drosophila* species are included in Supplemental Dataset 1.

| Gene family | *Dmel* | *Dpse* | *Dere* | *Dana* | *Dvir* | *Dmoj* | *Dgri* | *Spal* | *Shsu* | *Sgra* | *Smon* | *Sfla* |
|---|---|---|---|---|---|---|---|---|---|---|---|---|
| CYP450 | 89 | 79 | 88 | 93 | 81 | 78 | 85 | 72 | 69 | 68 | 67 | 73 |
| GST | 41 | 36 | 42 | 46 | 34 | 33 | 32 | 29 | 30 | 31 | 34 | 35 |
| UGT | 35 | 29 | 30 | 37 | 29 | 25 | 28 | 26 | 25 | 22 | 22 | 22 |
| GR | 65 | 58 | 56 | 67 | 62 | 61 | 75 | 78 | 72 | 64 | 57 | 61 |

| | | | | | | | | | | | | |
|---|---|---|---|---|---|---|---|---|---|---|---|---|
| IR | 58 | 58 | 63 | 62 | 54 | 54 | 59 | 55 | 63 | 52 | 51 | 47 |
| OBP | 52 | 45 | 50 | 50 | 41 | 42 | 50 | 41 | 41 | 33 | 34 | 34 |
| OR | 62 | 65 | 59 | 66 | 56 | 62 | 63 | 68 | 65 | 65 | 60 | 65 |
| PPK | 31 | 30 | 31 | 31 | 29 | 32 | 29 | 31 | 31 | 31 | 31 | 30 |
| TRP | 13 | 16 | 13 | 13 | 13 | 13 | 13 | 13 | 13 | 12 | 11 | 13 |
| Detoxification (all) | 165 | 144 | 160 | 176 | 144 | 136 | 145 | 127 | 124 | 121 | 123 | 130 |
| Chemosensation (all) | 281 | 272 | 272 | 289 | 255 | 264 | 289 | 286 | 285 | 257 | 244 | 250 |
| Random Gene Set | 186 | 183 | 184 | 206 | 212 | 201 | 238 | 196 | 195 | 189 | 193 | 198 |

**Table S4. Summarized CAFE model output for branch-specific estimates of rates of gene turnover, gene gain and loss across *Drosophila* and *Scaptomyza*.** Gene turnover rates were taken from the two-rate model. Models were run in triplicate, and only the model with the highest log likelihood is shown. Foreground branches: *Dpse = D. pseudoobscura, Dana = D. ananassae*, *Dere = D. erecta*, *Dmel = D. melanogaster*, *Dmoj = D. mojavensis*, *Dvir = D. virilis*, *Dgri = D. grimshawi*, *Shsu = S. hsui*, *Spal = S. pallida*, *Sgra = S. graminum*, *Smon = S. montana*, *Sfla = S. flava*, AncH = ancestral branch at the base of all three herbivorous *Scaptomyza*, CladeH = all branches in the herbivorous *Scaptomyza* clade. CHE = all chemosensory gene families, DET = all detoxification gene families, RAN = random set of gene families. Other gene family abbreviations as in the main text.

| | | | Gene turnover rates | | | | | | |
|---|---|---|---|---|---|---|---|---|---|
| family | branch | run | Background (lambda0) | Foreground (lambda1) | lnL (2 rate model) | lnL (1 rate model) | LRT | P value | FDR q |
| CHE | *Dpse* | r3 | 0.002478 | 0.000972 | -1600.45 | -1609.39 | 17.88256 | 2.35E-05 | 0.000282 |
| CHE | *Dana* | r3 | 0.002353 | 0.001792 | -1608.66 | -1609.39 | 1.467542 | 0.225734 | 0.144761 |
| CHE | *Dere* | r3 | 0.002292 | 0.00284 | -1609.16 | -1609.39 | 0.474728 | 0.49082 | 0.222641 |
| CHE | *Dmel* | r1 | 0.002273 | 0.003367 | -1608.48 | -1609.39 | 1.83063 | 0.176053 | 0.125751 |
| CHE | *Dmoj* | r2 | 0.002421 | 0.001126 | -1604.55 | -1609.39 | 9.686002 | 0.001857 | 0.005064 |
| CHE | *Dvir* | r3 | 0.002442 | 0.00093 | -1602.1 | -1609.39 | 14.58796 | 0.000134 | 0.000803 |
| CHE | *Dgri* | r2 | 0.002173 | 0.003767 | -1603.98 | -1609.39 | 10.82308 | 0.001002 | 0.003336 |
| CHE | *Shsu* | r1 | 0.00239 | 0.001161 | -1606.12 | -1609.39 | 6.54671 | 0.010508 | 0.017513 |
| CHE | *Spal* | r3 | 0.002302 | 0.00241 | -1609.37 | -1609.39 | 0.0416 | 0.838384 | 0.318374 |
| CHE | *Sgra* | r1 | 0.002236 | 0.005629 | -1604.03 | -1609.39 | 10.72623 | 0.001056 | 0.003336 |
| CHE | *Smon* | r2 | 0.00225 | 0.009616 | -1601.93 | -1609.39 | 14.93801 | 0.000111 | 0.000763 |
| CHE | *Sfla* | r1 | 0.002246 | 0.008902 | -1603.35 | -1609.39 | 12.08469 | 0.000508 | 0.002076 |
| CHE | AncH | r3 | 0.002227 | 0.004753 | -1604.7 | -1609.39 | 9.3928 | 0.002178 | 0.005446 |
| CHE | CladeH | r2 | 0.001947 | 0.005917 | -1578.97 | -1609.39 | 60.85218 | 6.15E-15 | 3.69E-13 |
| CYP | *Dpse* | r3 | 0.002666 | 0.00068 | -490.567 | -495.022 | 8.910192 | 0.002836 | 0.006544 |
| CYP | *Dana* | r1 | 0.002378 | 0.003565 | -494.272 | -495.022 | 1.500368 | 0.220615 | 0.143879 |
| CYP | *Dere* | r1 | 0.002453 | 0.002966 | -494.956 | -495.022 | 0.13093 | 0.71747 | 0.283212 |
| CYP | *Dmel* | r3 | 0.002516 | 0.001152 | -494.447 | -495.022 | 1.15034 | 0.283478 | 0.160566 |
| CYP | *Dmoj* | r1 | 0.002406 | 0.00316 | -494.658 | -495.022 | 0.727708 | 0.393627 | 0.196814 |
| CYP | *Dvir* | r3 | 0.002628 | 0.00097 | -492.464 | -495.022 | 5.114698 | 0.023724 | 0.031632 |
| CYP | *Dgri* | r2 | 0.002414 | 0.003067 | -494.749 | -495.022 | 0.545016 | 0.460361 | 0.218468 |

| | | | | | | | | | |
|---|---|---|---|---|---|---|---|---|---|
| CYP | *Shsu* | r2 | 0.002442 | 0.002898 | -494.919 | -495.022 | 0.20483 | 0.65085 | 0.262087 |
| CYP | *Spal* | r3 | 0.002599 | 0.000406 | -491.58 | -495.022 | 6.883502 | 0.008699 | 0.016312 |
| CYP | *Sgra* | r1 | 0.002481 | 0.002069 | -494.992 | -495.022 | 0.060234 | 0.806126 | 0.308074 |
| CYP | *Smon* | r2 | 0.002429 | 0.007994 | -493.801 | -495.022 | 2.440962 | 0.118204 | 0.098503 |
| CYP | *Sfla* | r1 | 0.00241 | 0.009308 | -493.064 | -495.022 | 3.914968 | 0.047858 | 0.053175 |
| CYP | AncH | r2 | 0.002438 | 0.003529 | -494.76 | -495.022 | 0.522666 | 0.469707 | 0.218468 |
| CYP | CladeH | r2 | 0.002324 | 0.003995 | -493.258 | -495.022 | 3.52772 | 0.060351 | 0.062432 |
| DET | *Dpse* | r1 | 0.002678 | 0.000875 | -884.941 | -891.126 | 12.36903 | 0.000437 | 0.002076 |
| DET | *Dana* | r1 | 0.002348 | 0.004122 | -888.35 | -891.126 | 5.5518 | 0.018462 | 0.025175 |
| DET | *Dere* | r3 | 0.002444 | 0.003797 | -890.451 | -891.126 | 1.349266 | 0.245406 | 0.153379 |
| DET | *Dmel* | r2 | 0.00247 | 0.003034 | -891.001 | -891.126 | 0.249896 | 0.617148 | 0.253623 |
| DET | *Dmoj* | r1 | 0.002484 | 0.002546 | -891.121 | -891.126 | 0.009398 | 0.922771 | 0.33711 |
| DET | *Dvir* | r2 | 0.00259 | 0.00152 | -889.437 | -891.126 | 3.376808 | 0.06612 | 0.06612 |
| DET | *Dgri* | r3 | 0.002456 | 0.002848 | -890.943 | -891.126 | 0.366094 | 0.545141 | 0.231975 |
| DET | *Shsu* | r2 | 0.002521 | 0.002028 | -890.883 | -891.126 | 0.485032 | 0.486152 | 0.222641 |
| DET | *Spal* | r2 | 0.002605 | 0.000523 | -886.119 | -891.126 | 10.01271 | 0.001555 | 0.004664 |
| DET | *Sgra* | r1 | 0.002495 | 0.002253 | -891.108 | -891.126 | 0.034448 | 0.852757 | 0.320482 |
| DET | *Smon* | r3 | 0.002457 | 0.006605 | -889.837 | -891.126 | 2.576302 | 0.108475 | 0.094326 |
| DET | *Sfla* | r1 | 0.002442 | 0.007808 | -888.847 | -891.126 | 4.558102 | 0.032763 | 0.040118 |
| DET | AncH | r2 | 0.002425 | 0.004617 | -889.468 | -891.126 | 3.314468 | 0.068673 | 0.067547 |
| DET | CladeH | r1 | 0.00231 | 0.004378 | -886.542 | -891.126 | 9.167798 | 0.002463 | 0.005911 |
| GR | *Dpse* | r2 | 0.00365 | 0.001074 | -422.036 | -425.447 | 6.822304 | 0.009003 | 0.016368 |
| GR | *Dana* | r1 | 0.00352 | 0.002069 | -424.821 | -425.447 | 1.252098 | 0.263152 | 0.160566 |
| GR | *Dere* | r2 | 0.00323 | 0.008628 | -423.337 | -425.447 | 4.219042 | 0.039973 | 0.047027 |
| GR | *Dmel* | r2 | 0.003302 | 0.005856 | -424.883 | -425.447 | 1.127724 | 0.288261 | 0.161642 |
| GR | *Dmoj* | r3 | 0.003637 | 0.00077 | -422.093 | -425.447 | 6.708198 | 0.009597 | 0.016936 |
| GR | *Dvir* | r1 | 0.00363 | 0.00083 | -421.87 | -425.447 | 7.154402 | 0.007478 | 0.014956 |
| GR | *Dgri* | r3 | 0.003168 | 0.005916 | -423.415 | -425.447 | 4.063412 | 0.043822 | 0.04961 |
| GR | *Shsu* | r3 | 0.003615 | 0.000499 | -421.858 | -425.447 | 7.17794 | 0.007381 | 0.014956 |
| GR | *Spal* | r2 | 0.003431 | 0.003 | -425.398 | -425.447 | 0.097086 | 0.755355 | 0.292395 |
| GR | *Sgra* | r2 | 0.003295 | 0.007017 | -424.447 | -425.447 | 1.99944 | 0.157357 | 0.117109 |
| GR | *Smon* | r3 | 0.003246 | 0.020437 | -421.056 | -425.447 | 8.780728 | 0.003044 | 0.006765 |
| GR | *Sfla* | r1 | 0.003296 | 0.012771 | -423.591 | -425.447 | 3.71067 | 0.054066 | 0.058981 |
| GR | AncH | r1 | 0.003338 | 0.005029 | -425.13 | -425.447 | 0.634228 | 0.425809 | 0.206036 |
| GR | CladeH | r2 | 0.002967 | 0.007189 | -419.996 | -425.447 | 10.90081 | 0.000961 | 0.003336 |
| GST | *Dpse* | r2 | 0.002387 | 0.000862 | -204.641 | -205.874 | 2.466218 | 0.116318 | 0.098342 |
| GST | *Dana* | r2 | 0.00221 | 0.002244 | -205.874 | -205.874 | 0.000628 | 0.980007 | 0.353337 |
| GST | *Dere* | r2 | 0.002178 | 0.003217 | -205.778 | -205.874 | 0.192124 | 0.661155 | 0.264462 |
| GST | *Dmel* | r1 | 0.002133 | 0.004424 | -205.416 | -205.874 | 0.915696 | 0.338608 | 0.183031 |
| GST | *Dmoj* | r3 | 0.002336 | 0.000964 | -204.997 | -205.874 | 1.7546 | 0.185299 | 0.127793 |
| GST | *Dvir* | r1 | 0.002193 | 0.00241 | -205.857 | -205.874 | 0.033526 | 0.854719 | 0.320482 |
| GST | *Dgri* | r3 | 0.002222 | 0.002114 | -205.87 | -205.874 | 0.008382 | 0.927053 | 0.33711 |
| GST | *Shsu* | r1 | 0.002337 | 1.03E-09 | -204.178 | -205.874 | 3.39199 | 0.065514 | 0.06612 |
| GST | *Spal* | r1 | 0.002322 | 6.31E-09 | -204.577 | -205.874 | 2.593832 | 0.107281 | 0.094326 |
| GST | *Sgra* | r2 | 0.002146 | 0.005712 | -205.255 | -205.874 | 1.237456 | 0.265962 | 0.160566 |
| GST | *Smon* | r2 | 0.002152 | 0.00995 | -204.976 | -205.874 | 1.79562 | 0.180243 | 0.125751 |
| GST | *Sfla* | r1 | 0.002137 | 0.010696 | -204.52 | -205.874 | 2.708376 | 0.099823 | 0.091022 |

| | | | | | | | | | |
|---|---|---|---|---|---|---|---|---|---|
| GST | AncH | r2 | 0.00212 | 0.005307 | -205.064 | -205.874 | 1.619792 | 0.203121 | 0.136935 |
| GST | CladeH | r1 | 0.001781 | 0.006787 | -199.785 | -205.874 | 12.17858 | 0.000483 | 0.002076 |
| IR | *Dpse* | r2 | 0.00266 | 0.000429 | -353.771 | -358.705 | 9.867476 | 0.001682 | 0.004806 |
| IR | *Dana* | r1 | 0.002474 | 0.001666 | -358.383 | -358.705 | 0.643252 | 0.422536 | 0.206036 |
| IR | *Dere* | r1 | 0.002425 | 0.001987 | -358.673 | -358.705 | 0.064434 | 0.79962 | 0.307546 |
| IR | *Dmel* | r3 | 0.002344 | 0.004422 | -358.107 | -358.705 | 1.195368 | 0.274249 | 0.160566 |
| IR | *Dmoj* | r2 | 0.00249 | 0.00156 | -358.191 | -358.705 | 1.028058 | 0.310615 | 0.171407 |
| IR | *Dvir* | r2 | 0.002552 | 0.000932 | -356.89 | -358.705 | 3.629712 | 0.056757 | 0.059744 |
| IR | *Dgri* | r3 | 0.002307 | 0.003498 | -358.058 | -358.705 | 1.29303 | 0.25549 | 0.158035 |
| IR | *Shsu* | r3 | 0.002355 | 0.003116 | -358.49 | -358.705 | 0.429824 | 0.512075 | 0.222641 |
| IR | *Spal* | r1 | 0.002253 | 0.005128 | -356.568 | -358.705 | 4.274262 | 0.038694 | 0.046433 |
| IR | *Sgra* | r2 | 0.002282 | 0.008019 | -355.744 | -358.705 | 5.922692 | 0.014947 | 0.021874 |
| IR | *Smon* | r1 | 0.002372 | 0.006858 | -357.982 | -358.705 | 1.445788 | 0.229205 | 0.144761 |
| IR | *Sfla* | r1 | 0.002362 | 0.009157 | -357.472 | -358.705 | 2.465488 | 0.116372 | 0.098342 |
| IR | AncH | r1 | 0.002312 | 0.005378 | -357.353 | -358.705 | 2.703564 | 0.100124 | 0.091022 |
| IR | CladeH | r3 | 0.001989 | 0.00646 | -351.612 | -358.705 | 14.18631 | 0.000166 | 0.000903 |
| OBP | *Dpse* | r2 | 0.001751 | 0.001398 | -232.248 | -232.36 | 0.224346 | 0.635748 | 0.257736 |
| OBP | *Dana* | r3 | 0.00161 | 0.00287 | -231.594 | -232.36 | 1.53267 | 0.215712 | 0.143808 |
| OBP | *Dere* | r3 | 0.001643 | 0.003724 | -231.609 | -232.36 | 1.503024 | 0.220207 | 0.143879 |
| OBP | *Dmel* | r3 | 0.001668 | 0.002906 | -232.058 | -232.36 | 0.603462 | 0.43726 | 0.209885 |
| OBP | *Dmoj* | r1 | 0.001837 | 0.000381 | -230.319 | -232.36 | 4.082154 | 0.043338 | 0.04961 |
| OBP | *Dvir* | r1 | 0.001797 | 0.000781 | -231.522 | -232.36 | 1.676526 | 0.195387 | 0.133218 |
| OBP | *Dgri* | r3 | 0.001586 | 0.002966 | -231.306 | -232.36 | 2.107716 | 0.146558 | 0.114661 |
| OBP | *Shsu* | r1 | 0.001836 | 6.84E-11 | -228.832 | -232.36 | 7.056802 | 0.007896 | 0.015284 |
| OBP | *Spal* | r1 | 0.00182 | 1.09E-11 | -229.249 | -232.36 | 6.222672 | 0.012612 | 0.019914 |
| OBP | *Sgra* | r2 | 0.001742 | 9.38E-10 | -231.462 | -232.36 | 1.796278 | 0.180163 | 0.125751 |
| OBP | *Smon* | r2 | 0.001723 | 6.45E-10 | -232.014 | -232.36 | 0.6927 | 0.405248 | 0.199302 |
| OBP | *Sfla* | r1 | 0.001722 | 8.74E-09 | -232.014 | -232.36 | 0.69271 | 0.405244 | 0.199302 |
| OBP | AncH | r2 | 0.001477 | 0.009788 | -224.919 | -232.36 | 14.88138 | 0.000114 | 0.000763 |
| OBP | CladeH | r1 | 0.001515 | 0.004071 | -229.524 | -232.36 | 5.671962 | 0.017238 | 0.024053 |
| OR | *Dpse* | r3 | 0.003389 | 0.000824 | -428.347 | -433.12 | 9.544764 | 0.002005 | 0.005231 |
| OR | *Dana* | r2 | 0.003154 | 0.00233 | -432.895 | -433.12 | 0.449084 | 0.50277 | 0.222641 |
| OR | *Dere* | r1 | 0.003154 | 0.000828 | -432.069 | -433.12 | 2.101566 | 0.147148 | 0.114661 |
| OR | *Dmel* | r3 | 0.003096 | 0.0029 | -433.114 | -433.12 | 0.01128 | 0.915418 | 0.33711 |
| OR | *Dmoj* | r3 | 0.003163 | 0.002376 | -432.876 | -433.12 | 0.48685 | 0.485336 | 0.222641 |
| OR | *Dvir* | r3 | 0.003283 | 0.001058 | -430.834 | -433.12 | 4.571372 | 0.032511 | 0.040118 |
| OR | *Dgri* | r2 | 0.002859 | 0.005858 | -430.617 | -433.12 | 5.006164 | 0.025257 | 0.032243 |
| OR | *Shsu* | r3 | 0.003152 | 0.002257 | -432.853 | -433.12 | 0.532964 | 0.465363 | 0.218468 |
| OR | *Spal* | r1 | 0.00314 | 0.002329 | -432.933 | -433.12 | 0.372782 | 0.541492 | 0.231975 |
| OR | *Sgra* | r3 | 0.002975 | 0.008222 | -431.283 | -433.12 | 3.67349 | 0.055284 | 0.059233 |
| OR | *Smon* | r1 | 0.003023 | 0.010984 | -431.561 | -433.12 | 3.117238 | 0.077468 | 0.073894 |
| OR | *Sfla* | r3 | 0.002983 | 0.014143 | -430.279 | -433.12 | 5.681066 | 0.017149 | 0.024053 |
| OR | AncH | r1 | 0.003082 | 0.003271 | -433.115 | -433.12 | 0.00924 | 0.923421 | 0.33711 |
| OR | CladeH | r1 | 0.002619 | 0.007735 | -424.739 | -433.12 | 16.76148 | 4.24E-05 | 0.000374 |
| PPK | *Dpse* | r2 | 0.000677 | 0.000397 | -78.7597 | -78.9114 | 0.30336 | 0.581784 | 0.240738 |
| PPK | *Dana* | r1 | 0.000699 | 3.13E-10 | -77.7974 | -78.9114 | 2.2279 | 0.135538 | 0.109896 |
| PPK | *Dere* | r3 | 0.000665 | 3.30E-10 | -78.5067 | -78.9114 | 0.809322 | 0.368321 | 0.19374 |

| | | | | | | | | | |
|---|---|---|---|---|---|---|---|---|---|
| PPK | *Dmel* | r2 | 0.000664 | 5.68E-10 | -78.5067 | -78.9114 | 0.809336 | 0.368317 | 0.19374 |
| PPK | *Dmoj* | r3 | 0.000663 | 0.000483 | -78.8625 | -78.9114 | 0.097726 | 0.754576 | 0.292395 |
| PPK | *Dvir* | r1 | 0.000607 | 0.001052 | -78.6886 | -78.9114 | 0.44547 | 0.504494 | 0.222641 |
| PPK | *Dgri* | r3 | 0.000607 | 0.001045 | -78.6945 | -78.9114 | 0.433694 | 0.510182 | 0.222641 |
| PPK | *Shsu* | r1 | 0.000695 | 3.77E-12 | -77.8823 | -78.9114 | 2.058058 | 0.151403 | 0.11499 |
| PPK | *Spal* | r1 | 0.000589 | 0.001515 | -78.313 | -78.9114 | 1.196796 | 0.273963 | 0.160566 |
| PPK | *Sgra* | r3 | 0.000612 | 0.002137 | -78.399 | -78.9114 | 1.024772 | 0.31139 | 0.171407 |
| PPK | *Smon* | r1 | 0.000651 | 1.59E-09 | -78.7938 | -78.9114 | 0.235064 | 0.627794 | 0.256242 |
| PPK | *Sfla* | r2 | 0.000604 | 0.005714 | -77.5962 | -78.9114 | 2.63024 | 0.104846 | 0.093892 |
| PPK | AncH | r3 | 0.000619 | 0.00145 | -78.6478 | -78.9114 | 0.527174 | 0.467797 | 0.218468 |
| PPK | CladeH | r3 | 0.000403 | 0.0031 | -73.2531 | -78.9114 | 11.31658 | 0.000768 | 0.002881 |
| RAN | *Dpse* | r1 | 0.002812 | 0.003372 | -1220.05 | -1220.45 | 0.788164 | 0.374656 | 0.19374 |
| RAN | *Dana* | r2 | 0.00279 | 0.003817 | -1219.5 | -1220.45 | 1.88542 | 0.169719 | 0.122689 |
| RAN | *Dere* | r2 | 0.002943 | 0.000615 | -1217.42 | -1220.45 | 6.058722 | 0.013838 | 0.020757 |
| RAN | *Dmel* | r1 | 0.002868 | 0.002867 | -1220.45 | -1220.45 | 0 | 1 | 0.357143 |
| RAN | *Dmoj* | r1 | 0.003006 | 0.001509 | -1217.22 | -1220.45 | 6.44811 | 0.011107 | 0.018011 |
| RAN | *Dvir* | r1 | 0.002968 | 0.0019 | -1218.89 | -1220.45 | 3.11471 | 0.077588 | 0.073894 |
| RAN | *Dgri* | r3 | 0.002591 | 0.007234 | -1209.44 | -1220.45 | 22.0014 | 2.72E-06 | 4.09E-05 |
| RAN | *Shsu* | r2 | 0.00307 | 1.52E-11 | -1200.26 | -1220.45 | 40.36223 | 2.11E-10 | 6.33E-09 |
| RAN | *Spal* | r1 | 0.003055 | 0.000304 | -1208.51 | -1220.45 | 23.8622 | 1.03E-06 | 2.07E-05 |
| RAN | *Sgra* | r2 | 0.002887 | 0.002263 | -1220.26 | -1220.45 | 0.378024 | 0.538663 | 0.231975 |
| RAN | *Smon* | r1 | 0.002878 | 0.001904 | -1220.28 | -1220.45 | 0.33883 | 0.560505 | 0.233544 |
| RAN | *Sfla* | r1 | 0.002826 | 0.006251 | -1218.92 | -1220.45 | 3.042752 | 0.081098 | 0.07603 |
| RAN | AncH | r2 | 0.002984 | 5.21E-05 | -1212.09 | -1220.45 | 16.70506 | 4.37E-05 | 0.000374 |
| RAN | CladeH | r1 | 0.00302 | 0.001742 | -1217.37 | -1220.45 | 6.15574 | 0.013099 | 0.020152 |
| TRP | *Dpse* | r1 | 0.000367 | 0.002743 | -32.2513 | -34.7644 | 5.026268 | 0.024966 | 0.032243 |
| TRP | *Dana* | r3 | 0.000698 | 5.03E-10 | -34.2983 | -34.7644 | 0.932136 | 0.334309 | 0.18235 |
| TRP | *Dere* | r2 | 0.000665 | 1.28E-09 | -34.5946 | -34.7644 | 0.339504 | 0.560116 | 0.233544 |
| TRP | *Dmel* | r3 | 0.000665 | 8.59E-10 | -34.5946 | -34.7644 | 0.339496 | 0.56012 | 0.233544 |
| TRP | *Dmoj* | r2 | 0.000711 | 8.31E-11 | -34.1897 | -34.7644 | 1.14944 | 0.283666 | 0.160566 |
| TRP | *Dvir* | r2 | 0.000712 | 2.88E-10 | -34.1897 | -34.7644 | 1.149438 | 0.283667 | 0.160566 |
| TRP | *Dgri* | r2 | 0.000711 | 3.69E-10 | -34.1896 | -34.7644 | 1.149638 | 0.283625 | 0.160566 |
| TRP | *Shsu* | r3 | 0.000694 | 9.49E-08 | -34.3329 | -34.7644 | 0.86308 | 0.352878 | 0.189042 |
| TRP | *Spal* | r3 | 0.000689 | 3.71E-10 | -34.3859 | -34.7644 | 0.75706 | 0.38425 | 0.19374 |
| TRP | *Sgra* | r1 | 0.00055 | 0.00463 | -33.6036 | -34.7644 | 2.321668 | 0.127583 | 0.104863 |
| TRP | *Smon* | r1 | 0.000432 | 0.030303 | -28.7414 | -34.7644 | 12.04592 | 0.000519 | 0.002076 |
| TRP | *Sfla* | r2 | 0.000651 | 7.12E-09 | -34.7139 | -34.7644 | 0.101076 | 0.750542 | 0.292395 |
| TRP | AncH | r2 | 0.000669 | 1.96E-10 | -34.543 | -34.7644 | 0.442844 | 0.505753 | 0.222641 |
| TRP | CladeH | r1 | 0.000353 | 0.003567 | -31.4631 | -34.7644 | 6.602602 | 0.010183 | 0.017457 |
| UGT | *Dpse* | r1 | 0.003064 | 0.001508 | -188.68 | -189.41 | 1.460858 | 0.226794 | 0.144761 |
| UGT | *Dana* | r1 | 0.00245 | 0.008234 | -185.585 | -189.41 | 7.650946 | 0.005674 | 0.012159 |
| UGT | *Dere* | r1 | 0.002755 | 0.007512 | -188.379 | -189.41 | 2.062396 | 0.150973 | 0.11499 |
| UGT | *Dmel* | r1 | 0.002737 | 0.007108 | -188.414 | -189.41 | 1.99233 | 0.158097 | 0.117109 |
| UGT | *Dmoj* | r3 | 0.00289 | 0.002922 | -189.41 | -189.41 | 0.00043 | 0.983456 | 0.353337 |
| UGT | *Dvir* | r1 | 0.002986 | 0.001987 | -189.18 | -189.41 | 0.461432 | 0.496955 | 0.222641 |
| UGT | *Dgri* | r1 | 0.002869 | 0.003139 | -189.395 | -189.41 | 0.031126 | 0.85996 | 0.320482 |
| UGT | *Shsu* | r2 | 0.002988 | 0.001627 | -189.013 | -189.41 | 0.79364 | 0.373002 | 0.19374 |

| | | | | | | | | | |
|---|---|---|---|---|---|---|---|---|---|
| UGT | *Spal* | r3 | 0.002992 | 0.001156 | -188.81 | -189.41 | 1.200398 | 0.273242 | 0.160566 |
| UGT | *Sgra* | r3 | 0.002961 | 1.80E-09 | -188.314 | -189.41 | 2.19204 | 0.138725 | 0.11098 |
| UGT | *Smon* | r1 | 0.002917 | 1.69E-09 | -189.029 | -189.41 | 0.761558 | 0.382841 | 0.19374 |
| UGT | *Sfla* | r2 | 0.002917 | 3.52E-10 | -189.029 | -189.41 | 0.761558 | 0.382841 | 0.19374 |
| UGT | AncH | r2 | 0.002772 | 0.007154 | -188.424 | -189.41 | 1.972274 | 0.160207 | 0.117224 |
| UGT | CladeH | r1 | 0.002942 | 0.002345 | -189.333 | -189.41 | 0.154956 | 0.693844 | 0.275699 |

**Table S5. Summarized CAFE model output for branch-specific estimates of rates of gene gain and loss across *Drosophila* and *Scaptomyza*.** Gene turnover rates were taken from the two-rate model. Models were run in triplicate, and only the model with the highest log likelihood is shown. Foreground branches: *Dpse = D. pseudoobscura, Dana = D. ananassae*, *Dere = D. erecta*, *Dmel = D. melanogaster*, *Dmoj = D. mojavensis*, *Dvir = D. virilis*, *Dgri = D. grimshawi*, *Shsu = S. hsui*, *Spal = S. pallida*, *Sgra = S. graminum*, *Smon = S. montana*, *Sfla = S. flava*, AncH = ancestral branch at the base of all three herbivorous *Scaptomyza*, CladeH = all branches in the herbivorous *Scaptomyza* clade. CHE = all chemosensory gene families, DET = all detoxification gene families, RAN = random set of gene families. Other gene family abbreviations as in the main text.

| | | | Gene duplication rates | | Gene loss rates | | | | | | |
|---|---|---|---|---|---|---|---|---|---|---|---|
| **family** | **branch** | **run** | **Background (lambda0)** | **Foreground (lambda1)** | **Background (mu0)** | **Foreground (mu1)** | **lnL (2 rate model)** | **lnL (1 rate model)** | **LRT** | **P value** | **FDR q** |
| CHE | *Dpse* | r1 | 0.002874 | 0.000984 | 0.002066 | 0.000972 | -1596.07 | -1596.07 | 0 | 1 | 0.42517 |
| CHE | *Dana* | r1 | 0.002668 | 0.00289 | 0.002009 | 0.000968 | -1603.19 | -1596.07 | -14.24 | 1 | 0.42517 |
| CHE | *Dere* | r1 | 0.002748 | 0.00136 | 0.001852 | 0.004241 | -1602.03 | -1596.07 | -11.92 | 1 | 0.42517 |
| CHE | *Dmel* | r1 | 0.00265 | 0.004088 | 0.00189 | 0.002783 | -1604.25 | -1596.07 | -16.36 | 1 | 0.42517 |
| CHE | *Dmoj* | r1 | 0.002865 | 0.001266 | 0.001994 | 0.001127 | -1600.48 | -1596.07 | -8.82 | 1 | 0.42517 |
| CHE | *Dvir* | r1 | 0.002929 | 0.000214 | 0.001994 | 0.001433 | -1593.15 | -1596.07 | 5.84 | 0.015666 | 0.019293 |
| CHE | *Dgri* | r1 | 0.002439 | 0.004972 | 0.001905 | 0.002314 | -1598.49 | -1596.07 | -4.84 | 1 | 0.42517 |
| CHE | *Shsu* | r1 | 0.002714 | 0.002357 | 0.002047 | 0.000355 | -1599.35 | -1596.07 | -6.56 | 1 | 0.42517 |
| CHE | *Spal* | r1 | 0.002626 | 0.003865 | 0.00196 | 0.001203 | -1603.08 | -1596.07 | -14.02 | 1 | 0.42517 |
| CHE | *Sgra* | r1 | 0.002616 | 0.006757 | 0.001853 | 0.004585 | -1599.64 | -1596.07 | -7.14 | 1 | 0.42517 |
| CHE | *Smon* | r1 | 0.002694 | 0.005552 | 0.001822 | 0.013604 | -1595.7 | -1596.07 | 0.74 | 0.389661 | 0.219026 |
| CHE | *Sfla* | r1 | 0.002633 | 0.009626 | 0.001877 | 0.007606 | -1599.29 | -1596.07 | -6.44 | 1 | 0.42517 |
| CHE | AncH | r1 | 0.002787 | 0.000652 | 0.001681 | 0.007868 | -1587.63 | -1596.07 | 16.88 | 3.98E-05 | 0.000284 |
| CHE | CladeH | r1 | 0.002516 | 0.004029 | 0.001403 | 0.007576 | -1564.66 | -1605.3 | 81.28 | 1.96E-19 | 1.40E-17 |
| CYP | *Dpse* | r1 | 0.003109 | 0.000738 | 0.002223 | 0.000802 | -489.252 | -493.465 | 8.426 | 0.003699 | 0.008523 |
| CYP | *Dana* | r1 | 0.002634 | 0.005448 | 0.002083 | 0.00165 | -491.313 | -493.465 | 4.304 | 0.038023 | 0.036212 |
| CYP | *Dere* | r1 | 0.00283 | 0.003535 | 0.002053 | 0.002274 | -493.4 | -493.465 | 0.13 | 0.718432 | 0.369184 |
| CYP | *Dmel* | r1 | 0.002847 | 0.002675 | 0.002122 | 1.63E-07 | -491.937 | -493.465 | 3.056 | 0.080439 | 0.062453 |
| CYP | *Dmoj* | r1 | 0.002827 | 0.003202 | 0.001952 | 0.003126 | -492.882 | -493.465 | 1.166 | 0.280225 | 0.173061 |
| CYP | *Dvir* | r1 | 0.002964 | 0.001602 | 0.002244 | 0.000438 | -490.588 | -493.465 | 5.754 | 0.016451 | 0.019865 |
| CYP | *Dgri* | r1 | 0.002639 | 0.005041 | 0.002149 | 0.000977 | -490.713 | -493.465 | 5.504 | 0.018973 | 0.020849 |
| CYP | *Shsu* | r1 | 0.002917 | 0.00175 | 0.001934 | 0.00403 | -491.766 | -493.465 | 3.398 | 0.065276 | 0.052388 |
| CYP | *Spal* | r2 | 0.002911 | 5.07E-11 | 0.002251 | 0.001019 | -489.448 | -493.465 | 8.034 | 0.004591 | 0.008911 |
| CYP | *Sgra* | r1 | 0.002881 | 0.001108 | 0.002053 | 0.003221 | -493.006 | -493.465 | 0.918 | 0.338001 | 0.199528 |
| CYP | *Smon* | r3 | 0.002855 | 3.40E-10 | 0.001969 | 0.016797 | -489.644 | -493.465 | 7.642 | 0.005702 | 0.010023 |

| | | | | | | | | | | | |
|---|---|---|---|---|---|---|---|---|---|---|---|
| CYP | *Sfla* | r3 | 0.002738 | 0.02432 | 0.002057 | 5.55E-09 | -489.453 | -493.465 | 8.024 | 0.004616 | 0.008911 |
| CYP | AncH | r2 | 0.002925 | 3.63E-09 | 0.001914 | 0.005813 | -490.428 | -493.465 | 6.074 | 0.013719 | 0.017816 |
| CYP | CladeH | r1 | 0.00281 | 0.003125 | 0.001802 | 0.00489 | -490.3 | -493.465 | 6.33 | 0.011871 | 0.016364 |
| DET | *Dpse* | r1 | 0.003216 | 0.000805 | 0.002165 | 0.000862 | -881.361 | -888.417 | 14.112 | 0.000172 | 0.00082 |
| DET | *Dana* | r1 | 0.002622 | 0.006844 | 0.002018 | 0.001545 | -880.845 | -888.417 | 15.144 | 9.96E-05 | 0.000547 |
| DET | *Dere* | r1 | 0.002901 | 0.004168 | 0.001957 | 0.003882 | -887.607 | -888.417 | 1.62 | 0.203092 | 0.135575 |
| DET | *Dmel* | r1 | 0.002913 | 0.004906 | 0.002043 | 2.20E-10 | -886.366 | -888.417 | 4.102 | 0.042833 | 0.037771 |
| DET | *Dmoj* | r1 | 0.002935 | 0.002231 | 0.00201 | 0.002739 | -887.604 | -888.417 | 1.626 | 0.202257 | 0.135575 |
| DET | *Dvir* | r1 | 0.003013 | 0.00225 | 0.002147 | 0.00077 | -886.213 | -888.417 | 4.408 | 0.035771 | 0.034528 |
| DET | *Dgri* | r1 | 0.002753 | 0.004086 | 0.002126 | 0.001585 | -887.04 | -888.417 | 2.754 | 0.097011 | 0.072941 |
| DET | *Shsu* | r1 | 0.00307 | 0.000966 | 0.001951 | 0.003014 | -885.259 | -888.417 | 6.316 | 0.011965 | 0.016364 |
| DET | *Spal* | r1 | 0.003025 | 0.000426 | 0.002155 | 0.000607 | -883.036 | -888.417 | 10.762 | 0.001036 | 0.003364 |
| DET | *Sgra* | r1 | 0.002887 | 0.001866 | 0.002076 | 0.002747 | -888.169 | -888.417 | 0.496 | 0.481263 | 0.258466 |
| DET | *Smon* | r3 | 0.002914 | 1.64E-10 | 0.002025 | 0.012327 | -884.604 | -888.417 | 7.626 | 0.005753 | 0.010023 |
| DET | *Sfla* | r1 | 0.002793 | 0.013552 | 0.00207 | 0.00171 | -884.939 | -888.417 | 6.956 | 0.008354 | 0.013745 |
| DET | AncH | r1 | 0.002999 | 0.001559 | 0.001833 | 0.006854 | -883.092 | -888.417 | 10.65 | 0.001101 | 0.003418 |
| DET | CladeH | r1 | 0.002837 | 0.003999 | 0.001765 | 0.004806 | -882.657 | -888.417 | 11.52 | 0.000689 | 0.002588 |
| GR | *Dpse* | r1 | 0.004913 | 0.000913 | 0.002423 | 0.0013 | -417.72 | -421.386 | 7.332 | 0.006774 | 0.01152 |
| GR | *Dana* | r2 | 0.004626 | 0.004675 | 0.002416 | 1.13E-10 | -418.765 | -421.386 | 5.242 | 0.022048 | 0.023861 |
| GR | *Dere* | r1 | 0.004608 | 0.002023 | 0.001984 | 0.01336 | -415.203 | -421.386 | 12.366 | 0.000437 | 0.001952 |
| GR | *Dmel* | r1 | 0.004491 | 0.010971 | 0.00221 | 0.00126 | -419.78 | -421.386 | 3.212 | 0.0731 | 0.057378 |
| GR | *Dmoj* | r1 | 0.005011 | 0.001069 | 0.002358 | 0.000684 | -417.34 | -421.386 | 8.092 | 0.004446 | 0.008911 |
| GR | *Dvir* | r1 | 0.004853 | 0.001152 | 0.002469 | 0.001092 | -418.611 | -421.386 | 5.55 | 0.018481 | 0.020626 |
| GR | *Dgri* | r1 | 0.004263 | 0.008228 | 0.00222 | 0.002366 | -419.072 | -421.386 | 4.628 | 0.031454 | 0.031205 |
| GR | *Shsu* | r3 | 0.004692 | 0.002521 | 0.002481 | 1.34E-10 | -418.019 | -421.386 | 6.734 | 0.009459 | 0.014376 |
| GR | *Spal* | r1 | 0.00445 | 0.007555 | 0.002369 | 1.81E-10 | -417.291 | -421.386 | 8.19 | 0.004212 | 0.008911 |
| GR | *Sgra* | r1 | 0.004574 | 0.008854 | 0.00208 | 0.006587 | -419.765 | -421.386 | 3.242 | 0.071773 | 0.056963 |
| GR | *Smon* | r1 | 0.004588 | 0.007088 | 0.002006 | 0.030252 | -414.305 | -421.386 | 14.162 | 0.000168 | 0.00082 |
| GR | *Sfla* | r1 | 0.004533 | 0.01516 | 0.002165 | 0.011094 | -419.468 | -421.386 | 3.836 | 0.050163 | 0.042154 |
| GR | AncH | r2 | 0.004755 | 3.54E-11 | 0.001968 | 0.008549 | -415.936 | -421.386 | 10.9 | 0.000962 | 0.003271 |
| GR | CladeH | r1 | 0.004427 | 0.003239 | 0.001469 | 0.009988 | -407.157 | -421.386 | 28.458 | 9.58E-08 | 1.71E-06 |
| GST | *Dpse* | r1 | 0.003302 | 2.51E-10 | 0.0016 | 0.000796 | -201.569 | -204.714 | 6.29 | 0.012142 | 0.016364 |
| GST | *Dana* | r1 | 0.00268 | 0.006183 | 0.001561 | 1.98E-10 | -201.864 | -204.714 | 5.7 | 0.016965 | 0.019865 |
| GST | *Dere* | r1 | 0.002872 | 0.006681 | 0.00143 | 0.001078 | -204.142 | -204.714 | 1.144 | 0.284809 | 0.173876 |
| GST | *Dmel* | r1 | 0.002546 | 0.004277 | 0.001689 | 0.004645 | -204.515 | -204.714 | 0.398 | 0.528124 | 0.281516 |
| GST | *Dmoj* | r1 | 0.002787 | 0.000901 | 0.001857 | 0.001027 | -204.122 | -204.714 | 1.184 | 0.276543 | 0.173061 |
| GST | *Dvir* | r1 | 0.00299 | 0.002768 | 0.001492 | 0.001148 | -204.674 | -204.714 | 0.08 | 0.777297 | 0.39658 |
| GST | *Dgri* | r1 | 0.00305 | 0.0021 | 0.001473 | 0.001266 | -204.565 | -204.714 | 0.298 | 0.585139 | 0.307321 |
| GST | *Shsu* | r1 | 0.003117 | 4.77E-10 | 0.001517 | 0.000352 | -202.602 | -204.714 | 4.224 | 0.039856 | 0.037459 |
| GST | *Spal* | r2 | 0.003086 | 1.66E-11 | 0.001427 | 0.000331 | -202.654 | -204.714 | 4.12 | 0.042379 | 0.037771 |
| GST | *Sgra* | r1 | 0.00288 | 0.006216 | 0.001435 | 0.00493 | -204.202 | -204.714 | 1.024 | 0.311572 | 0.187018 |
| GST | *Smon* | r1 | 0.002958 | 0.009684 | 0.001327 | 0.011536 | -203.316 | -204.714 | 2.796 | 0.0945 | 0.071808 |
| GST | *Sfla* | r1 | 0.002871 | 0.012382 | 0.001409 | 0.00909 | -203.431 | -204.714 | 2.566 | 0.109183 | 0.081238 |
| GST | AncH | r1 | 0.002854 | 0.00576 | 0.001425 | 0.003556 | -204.151 | -204.714 | 1.126 | 0.28863 | 0.174716 |
| GST | CladeH | r2 | 0.001989 | 0.009158 | 0.001556 | 0.003969 | -198.723 | -204.714 | 11.982 | 0.000537 | 0.002132 |
| IR | *Dpse* | r1 | 0.002763 | 0.00034 | 0.002559 | 0.000501 | -353.681 | -358.644 | 9.926 | 0.00163 | 0.004311 |
| IR | *Dana* | r1 | 0.00252 | 0.002107 | 0.002413 | 0.001362 | -358.258 | -358.644 | 0.772 | 0.379599 | 0.215929 |

| | | | | | | | | | | | |
|---|---|---|---|---|---|---|---|---|---|---|---|
| IR | *Dere* | r1 | 0.002501 | 0.003393 | 0.002353 | 0.000521 | -358.225 | -358.644 | 0.838 | 0.359969 | 0.207356 |
| IR | *Dmel* | r3 | 0.002522 | 4.99E-10 | 0.002165 | 0.007919 | -355.434 | -358.644 | 6.42 | 0.011284 | 0.01612 |
| IR | *Dmoj* | r1 | 0.002425 | 0.000627 | 0.00254 | 0.001851 | -357.924 | -358.644 | 1.44 | 0.230139 | 0.152209 |
| IR | *Dvir* | r1 | 0.002689 | 0.000552 | 0.002417 | 0.001262 | -356.557 | -358.644 | 4.174 | 0.041049 | 0.037771 |
| IR | *Dgri* | r1 | 0.002301 | 0.004452 | 0.002308 | 0.002339 | -357.493 | -358.644 | 2.302 | 0.129208 | 0.09449 |
| IR | *Shsu* | r1 | 0.002295 | 0.00546 | 0.002388 | 0.000807 | -355.832 | -358.644 | 5.624 | 0.017716 | 0.020132 |
| IR | *Spal* | r1 | 0.00238 | 0.004653 | 0.002123 | 0.005591 | -356.39 | -358.644 | 4.508 | 0.033737 | 0.03301 |
| IR | *Sgra* | r1 | 0.002376 | 0.00798 | 0.002185 | 0.008013 | -355.678 | -358.644 | 5.932 | 0.014868 | 0.018632 |
| IR | *Smon* | r1 | 0.002418 | 0.013422 | 0.002322 | 2.05E-11 | -356.66 | -358.644 | 3.968 | 0.046373 | 0.039908 |
| IR | *Sfla* | r2 | 0.002224 | 4.75E-06 | 0.002394 | 0.023344 | -355.52 | -358.644 | 6.248 | 0.012433 | 0.016446 |
| IR | AncH | r1 | 0.002485 | 0.00165 | 0.002121 | 0.009452 | -355.26 | -358.644 | 6.768 | 0.009281 | 0.014376 |
| IR | CladeH | r2 | 0.002057 | 0.00309 | 0.001892 | 0.009968 | -348.796 | -358.644 | 19.696 | 9.08E-06 | 9.26E-05 |
| OBP | *Dpse* | r1 | 0.001415 | 0.000551 | 0.00202 | 0.002606 | -230.713 | -231.392 | 1.358 | 0.243884 | 0.158062 |
| OBP | *Dana* | r1 | 0.001303 | 0.003082 | 0.00192 | 0.002637 | -230.718 | -231.392 | 1.348 | 0.245628 | 0.158062 |
| OBP | *Dere* | r1 | 0.001274 | 0.002071 | 0.001992 | 0.005761 | -230.408 | -231.392 | 1.968 | 0.16066 | 0.112507 |
| OBP | *Dmel* | r1 | 0.001249 | 0.003636 | 0.002072 | 0.002205 | -230.811 | -231.392 | 1.162 | 0.281051 | 0.173061 |
| OBP | *Dmoj* | r1 | 0.001251 | 7.31E-11 | 0.002203 | 0.002759 | -230.329 | -231.392 | 2.126 | 0.144818 | 0.103442 |
| OBP | *Dvir* | r3 | 0.001433 | 5.81E-11 | 0.002134 | 0.001109 | -229.638 | -231.392 | 3.508 | 0.061073 | 0.05033 |
| OBP | *Dgri* | r1 | 0.000863 | 0.005371 | 0.002295 | 3.35E-08 | -222.089 | -231.392 | 18.606 | 1.61E-05 | 0.000143 |
| OBP | *Shsu* | r3 | 0.001379 | 4.34E-05 | 0.002227 | 6.40E-11 | -227.926 | -231.392 | 6.932 | 0.008467 | 0.013745 |
| OBP | *Spal* | r1 | 0.001441 | 8.70E-11 | 0.002112 | 7.98E-05 | -228.414 | -231.392 | 5.956 | 0.014667 | 0.018632 |
| OBP | *Sgra* | r3 | 0.001588 | 0.002101 | 0.001985 | 1.97E-10 | -231.495 | -231.392 | -0.206 | 1 | 0.42517 |
| OBP | *Smon* | r2 | 0.001328 | 4.86E-05 | 0.002113 | 1.52E-05 | -231.053 | -231.392 | 0.678 | 0.410276 | 0.225427 |
| OBP | *Sfla* | r2 | 0.001323 | 0.003131 | 0.002107 | 2.80E-07 | -231.361 | -231.392 | 0.062 | 0.803362 | 0.406972 |
| OBP | AncH | r1 | 0.001467 | 8.01E-10 | 0.001485 | 0.016657 | -220.141 | -231.392 | 22.502 | 2.10E-06 | 2.50E-05 |
| OBP | CladeH | r1 | 0.001472 | 0.000906 | 0.001562 | 0.007163 | -226.333 | -231.392 | 10.118 | 0.001468 | 0.004195 |
| OR | *Dpse* | r1 | 0.004585 | 0.001615 | 0.00207 | 0.000419 | -423.588 | -427.648 | 8.12 | 0.004378 | 0.008911 |
| OR | *Dana* | r1 | 0.004285 | 0.004164 | 0.001951 | 0.000733 | -427.009 | -427.648 | 1.278 | 0.258271 | 0.164714 |
| OR | *Dere* | r3 | 0.004548 | 4.46E-10 | 0.001769 | 0.001693 | -425.282 | -427.648 | 4.732 | 0.029606 | 0.029785 |
| OR | *Dmel* | r1 | 0.004267 | 0.005079 | 0.00186 | 0.001402 | -427.58 | -427.648 | 0.136 | 0.71229 | 0.36868 |
| OR | *Dmoj* | r2 | 0.004316 | 0.004014 | 0.001857 | 0.001769 | -427.63 | -427.648 | 0.036 | 0.849515 | 0.424333 |
| OR | *Dvir* | r2 | 0.004611 | 1.28E-09 | 0.001945 | 0.001363 | -422.113 | -427.648 | 11.07 | 0.000877 | 0.003133 |
| OR | *Dgri* | r1 | 0.004046 | 0.006967 | 0.001601 | 0.00494 | -424.337 | -427.648 | 6.622 | 0.010073 | 0.014989 |
| OR | *Shsu* | r1 | 0.004311 | 0.003893 | 0.001932 | 0.000811 | -427.156 | -427.648 | 0.984 | 0.321213 | 0.191198 |
| OR | *Spal* | r1 | 0.004202 | 0.005567 | 0.00196 | 3.28E-08 | -425.232 | -427.648 | 4.832 | 0.027936 | 0.028506 |
| OR | *Sgra* | r2 | 0.004127 | 0.012972 | 0.001795 | 2.80E-10 | -424.784 | -427.648 | 5.728 | 0.016696 | 0.019865 |
| OR | *Smon* | r1 | 0.004375 | 0.006304 | 0.001677 | 0.016227 | -425.149 | -427.648 | 4.998 | 0.025377 | 0.026656 |
| OR | *Sfla* | r1 | 0.00411 | 0.033384 | 0.001816 | 4.60E-09 | -422.82 | -427.648 | 9.656 | 0.001887 | 0.004649 |
| OR | AncH | r1 | 0.004606 | 0.000138 | 0.001618 | 0.003677 | -425.944 | -427.648 | 3.408 | 0.064881 | 0.052388 |
| OR | CladeH | r1 | 0.003734 | 0.00927 | 0.001532 | 0.005383 | -420.036 | -427.648 | 15.224 | 9.55E-05 | 0.000547 |
| PPK | *Dpse* | r1 | 0.000484 | 0.000132 | 0.000847 | 0.000782 | -77.8901 | -78.3199 | 0.8596 | 0.353851 | 0.205488 |
| PPK | *Dana* | r2 | 0.000447 | 0.00298 | 0.001012 | 6.65E-10 | -79.794 | -78.3199 | -2.9482 | 1 | 0.42517 |
| PPK | *Dere* | r1 | 0.00048 | 0.002508 | 0.000799 | 9.39E-07 | -78.7137 | -78.3199 | -0.7876 | 1 | 0.42517 |
| PPK | *Dmel* | r3 | 0.000475 | 1.28E-09 | 0.000857 | 0.002395 | -78.6572 | -78.3199 | -0.6746 | 1 | 0.42517 |
| PPK | *Dmoj* | r1 | 0.000407 | 0.002656 | 0.000918 | 1.63E-10 | -77.936 | -78.3199 | 0.7678 | 0.380899 | 0.215929 |
| PPK | *Dvir* | r2 | 0.00047 | 2.78E-10 | 0.000752 | 0.002009 | -77.1715 | -78.3199 | 2.2968 | 0.129641 | 0.09449 |
| PPK | *Dgri* | r3 | 0.000426 | 4.18E-11 | 0.000768 | 0.001942 | -77.2567 | -78.3199 | 2.1264 | 0.14478 | 0.103442 |

| | | | | | | | | | | | |
|---|---|---|---|---|---|---|---|---|---|---|---|
| PPK | *Shsu* | r1 | 0.000475 | 3.11E-11 | 0.001158 | 0.007822 | -83.6806 | -78.3199 | -10.7214 | 1 | 0.42517 |
| PPK | *Spal* | r1 | 0.000392 | 0.001514 | 0.000786 | 0.001516 | -77.6213 | -78.3199 | 1.3972 | 0.237193 | 0.155434 |
| PPK | *Sgra* | r2 | 0.000378 | 0.003643 | 0.000843 | 2.25E-10 | -76.8932 | -78.3199 | 2.8534 | 0.091181 | 0.070031 |
| PPK | *Smon* | r3 | 0.000461 | 3.51E-09 | 0.000837 | 0.001685 | -78.3563 | -78.3199 | -0.0728 | 1 | 0.42517 |
| PPK | *Sfla* | r1 | 0.000465 | 1.04E-09 | 0.000738 | 0.009774 | -76.569 | -78.3199 | 3.5018 | 0.061302 | 0.05033 |
| PPK | AncH | r1 | 0.000477 | 1.99E-10 | 0.00076 | 0.002536 | -77.7372 | -78.3199 | 1.1654 | 0.280348 | 0.173061 |
| PPK | CladeH | r1 | 0.000202 | 0.002972 | 0.000602 | 0.003237 | -72.208 | -78.3199 | 12.2238 | 0.000472 | 0.001983 |
| RAN | *Dpse* | r1 | 0.003544 | 0.003111 | 0.002114 | 0.003456 | -1215.13 | -1217.19 | 4.12 | 0.042379 | 0.037771 |
| RAN | *Dana* | r1 | 0.003326 | 0.005339 | 0.00233 | 0.00126 | -1213.76 | -1217.19 | 6.86 | 0.008815 | 0.013992 |
| RAN | *Dere* | r1 | 0.003591 | 0.000347 | 0.002319 | 0.000732 | -1213.28 | -1217.19 | 7.82 | 0.005167 | 0.009713 |
| RAN | *Dmel* | r1 | 0.003486 | 0.003434 | 0.002248 | 0.002767 | -1217.11 | -1217.19 | 0.16 | 0.689157 | 0.35931 |
| RAN | *Dmoj* | r1 | 0.003598 | 0.001485 | 0.002469 | 0.00118 | -1213.37 | -1217.19 | 7.64 | 0.005709 | 0.010023 |
| RAN | *Dvir* | r1 | 0.003419 | 0.002851 | 0.002547 | 0.000769 | -1214.38 | -1217.19 | 5.62 | 0.017757 | 0.020132 |
| RAN | *Dgri* | r1 | 0.002958 | 0.00947 | 0.002225 | 0.002625 | -1198.52 | -1217.19 | 37.34 | 9.92E-10 | 3.54E-08 |
| RAN | *Shsu* | r2 | 0.002957 | 3.75E-07 | 0.002635 | 6.47E-11 | -1199.93 | -1217.19 | 34.52 | 4.22E-09 | 1.00E-07 |
| RAN | *Spal* | r1 | 0.003668 | 0.000496 | 0.002443 | 8.48E-05 | -1205.43 | -1217.19 | 23.52 | 1.24E-06 | 1.77E-05 |
| RAN | *Sgra* | r1 | 0.003543 | 5.17E-08 | 0.002229 | 0.00396 | -1212.28 | -1217.19 | 9.82 | 0.001726 | 0.004404 |
| RAN | *Smon* | r1 | 0.003508 | 0.001572 | 0.002267 | 0.002299 | -1216.84 | -1217.19 | 0.7 | 0.402784 | 0.223025 |
| RAN | *Sfla* | r1 | 0.003415 | 0.010289 | 0.002258 | 0.001883 | -1214.74 | -1217.19 | 4.9 | 0.026857 | 0.027802 |
| RAN | AncH | r2 | 0.003576 | 0.000305 | 0.002358 | 6.68E-11 | -1208.72 | -1217.19 | 16.94 | 3.86E-05 | 0.000284 |
| RAN | CladeH | r1 | 0.003681 | 0.001555 | 0.00233 | 0.001918 | -1213.1 | -1217.19 | 8.18 | 0.004235 | 0.008911 |
| TRP | *Dpse* | r2 | 0.00016 | 0.010327 | 0.000739 | 0.012642 | -35.1344 | -34.7644 | -0.74 | 1 | 0.42517 |
| TRP | *Dana* | r3 | 0.00063 | 0.006731 | 0.000705 | 5.81E-08 | -36.6751 | -34.7644 | -3.8214 | 1 | 0.42517 |
| TRP | *Dere* | r1 | 0.00064 | 0.002005 | 0.000676 | 1.13E-05 | -34.8587 | -34.7644 | -0.1886 | 1 | 0.42517 |
| TRP | *Dmel* | r3 | 0.000667 | 6.01E-05 | 0.000728 | 9.73E-10 | -34.6143 | -34.7644 | 0.3002 | 0.583757 | 0.307321 |
| TRP | *Dmoj* | r2 | 0.000715 | 3.33E-09 | 0.000687 | 0.004211 | -35.9979 | -34.7644 | -2.467 | 1 | 0.42517 |
| TRP | *Dvir* | r3 | 0.000705 | 0.000336 | 0.000715 | 4.45E-09 | -34.3337 | -34.7644 | 0.8614 | 0.353347 | 0.205488 |
| TRP | *Dgri* | r1 | 0.000806 | 0.001654 | 0.000627 | 4.32E-12 | -34.9504 | -34.7644 | -0.372 | 1 | 0.42517 |
| TRP | *Shsu* | r1 | 0.000682 | 0.00043 | 0.000769 | 0.004947 | -36.0792 | -34.7644 | -2.6296 | 1 | 0.42517 |
| TRP | *Spal* | r2 | 0.000689 | 4.52E-05 | 0.000692 | 3.45E-09 | -34.3988 | -34.7644 | 0.7312 | 0.392494 | 0.219026 |
| TRP | *Sgra* | r3 | 0.000662 | 8.91E-10 | 0.00045 | 0.008801 | -32.8352 | -34.7644 | 3.8584 | 0.049498 | 0.04209 |
| TRP | *Smon* | r3 | 0.000643 | 3.56E-06 | 0.000217 | 0.042931 | -26.99 | -34.7644 | 15.5488 | 8.04E-05 | 0.000522 |
| TRP | *Sfla* | r3 | 0.00065 | 0.002696 | 0.000653 | 1.01E-08 | -34.8189 | -34.7644 | -0.109 | 1 | 0.42517 |
| TRP | AncH | r1 | 0.000629 | 0.003588 | 0.001109 | 5.93E-11 | -35.6017 | -34.7644 | -1.6746 | 1 | 0.42517 |
| TRP | CladeH | r2 | 0.000664 | 0.003953 | 5.23E-11 | 0.013653 | -29.7643 | -34.7644 | 10.0002 | 0.001565 | 0.0043 |
| UGT | *Dpse* | r1 | 0.003449 | 0.001732 | 0.002672 | 0.001088 | -188.223 | -189.09 | 1.734 | 0.187901 | 0.127824 |
| UGT | *Dana* | r1 | 0.002349 | 0.011559 | 0.002526 | 0.003671 | -183.831 | -189.09 | 10.518 | 0.001182 | 0.003518 |
| UGT | *Dere* | r1 | 0.003196 | 0.003119 | 0.002284 | 0.011921 | -187.051 | -189.09 | 4.078 | 0.043445 | 0.037844 |
| UGT | *Dmel* | r2 | 0.002975 | 0.012055 | 0.002499 | 1.16E-09 | -186.492 | -189.09 | 5.196 | 0.022639 | 0.024135 |
| UGT | *Dmoj* | r1 | 0.003422 | 0.001276 | 0.002364 | 0.003843 | -188.15 | -189.09 | 1.88 | 0.170334 | 0.116988 |
| UGT | *Dvir* | r1 | 0.003224 | 0.003375 | 0.002637 | 0.001279 | -188.766 | -189.09 | 0.648 | 0.420829 | 0.227721 |
| UGT | *Dgri* | r1 | 0.003207 | 0.00363 | 0.002505 | 0.002535 | -189.07 | -189.09 | 0.04 | 0.841481 | 0.42328 |
| UGT | *Shsu* | r1 | 0.003474 | 1.62E-10 | 0.002448 | 0.003085 | -187.035 | -189.09 | 4.11 | 0.04263 | 0.037771 |
| UGT | *Spal* | r3 | 0.003342 | 0.001787 | 0.002622 | 1.76E-09 | -188.127 | -189.09 | 1.926 | 0.165197 | 0.114561 |
| UGT | *Sgra* | r3 | 0.003322 | 0.000314 | 0.002568 | 7.44E-10 | -188.056 | -189.09 | 2.068 | 0.150419 | 0.106378 |
| UGT | *Smon* | r2 | 0.003253 | 2.73E-10 | 0.002537 | 0.013236 | -189.584 | -189.09 | -0.988 | 1 | 0.42517 |
| UGT | *Sfla* | r1 | 0.003308 | 1.82E-11 | 0.002588 | 0.000627 | -188.759 | -189.09 | 0.662 | 0.415855 | 0.226747 |

| | | | | | | | | | | | |
|---|---|---|---|---|---|---|---|---|---|---|---|
| UGT | AncH | r1 | 0.003329 | 1.19E-09 | 0.002161 | 0.01371 | -184.839 | -189.09 | 8.502 | 0.003548 | 0.008447 |
| UGT | CladeH | r1 | 0.00352 | 1.04E-10 | 0.002299 | 0.005023 | -185.864 | -189.09 | 6.452 | 0.011083 | 0.01612 |

**Table S6. Orthologous genes identified as significantly expanding or contracting across the drosophilid tree.** Family-wide *P*-value indicates whether significant shifts in turnover rate were detected within a given orthology group. Viterbi *P*-values indicate whether turnover rate was significantly different at individual branches within the phylogeny. Branch IDs (0-22) correspond to the labels shown in Fig. S14. Viterbi *P*-values <0.05 are highlighted in bold black text/pale green cells for all branches, except for the ancestral herbivore branch in which these values are highlighted in white bold text/dark green cells. For genes from the random gene set, the genes are listed by cluster IDs and gene names in parentheses.

| | | **Viterbi *P*-values for individual branches** | | | | | | | | | | | | | | | | | | | | | |
|---|---|---|---|---|---|---|---|---|---|---|---|---|---|---|---|---|---|---|---|---|---|---|---|
| **Gene / Cluster ID** | **Family-wide *P*-value** | **0** | **3** | **2** | **4** | **1** | **6** | **5** | **19** | **8** | **17** | **10** | **12** | **11** | **14** | **13** | **16** | **15** | **18** | **9** | **21** | **20** | **22** |
| *G36abc* | 0.0329 | **0.019** | 0.588 | **0.019** | 0.071 | 0.051 | 0.606 | 0.547 | **0.018** | 0.5 | 0.5 | 0.5 | 0.5 | 0.5 | 0.5 | 0.5 | 0.5 | 0.5 | 0.5 | 0.5 | 0.5 | 0.5 | 0.5 |
| *Gr28b* | 0.0003 | 0.572 | 0.547 | 0.528 | **0** | 0.539 | 0.606 | **0.036** | **0.001** | 0.802 | 0.618 | 0.587 | 0.587 | 0.234 | 0.732 | 0.785 | 0.744 | 0.534 | 0.762 | 0.133 | 0.59 | 0.776 | 0.776 |
| *Gr39aA* | 0 | 0.572 | 0.547 | **0.009** | 0.528 | **0.028** | 0.259 | 0.057 | **0.018** | 0.111 | **0.01** | **0.008** | 0.602 | 0.265 | 0.156 | 0.807 | 0.096 | 0.539 | 0.225 | 0.601 | 0.59 | 0.776 | 0.413 |
| *Gr59cd* | 0.028 | 0.63 | 0.588 | 0.554 | 0.554 | 0.574 | 0.682 | 0.588 | 0.123 | 0.22 | 0.133 | 0.571 | 0.571 | 0.629 | 0.232 | 0.186 | **0.024** | 0.534 | 0.386 | **0.004** | 0.557 | 0.703 | 0.291 |
| *Gr85a* | 0 | 0.096 | 0.547 | 0.528 | 0.528 | 0.539 | 0.606 | **0.036** | **0.009** | **0** | 0.601 | 0.111 | **0.029** | 0.654 | **0.003** | 0.759 | 0.31 | 0.528 | 0.161 | 0.109 | 0.574 | 0.744 | 0.357 |
| ***Gr92a93bcd*** | 0.0001 | **0.001** | 0.624 | **0** | 0.103 | 0.605 | 0.737 | 0.117 | 0.592 | 0.652 | 0.544 | 0.555 | 0.069 | 0.602 | 0.642 | **0.046** | 0.609 | 0.512 | 0.622 | 0.544 | 0.539 | **0.028** | 0.083 |
| *Gr98a* | 0.04965 | 0.572 | 0.547 | 0.528 | 0.528 | 0.539 | 0.053 | 0.547 | 0.065 | 0.652 | **0.002** | 0.571 | 0.091 | 0.629 | 0.676 | 0.726 | 0.687 | 0.523 | 0.705 | 0.544 | 0.539 | 0.652 | 0.212 |
| *Ir47abc94abc* | 0.00545 | **0.023** | 0.611 | **0.039** | 0.57 | 0.594 | 0.334 | 0.611 | 0.616 | 0.608 | 0.53 | 0.519 | 0.519 | 0.537 | 0.554 | 0.172 | **0.043** | 0.508 | **0.049** | **0.001** | 0.551 | 0.683 | 0.124 |
| *IR52* | 0.0185 | 0.659 | 0.611 | 0.106 | 0.57 | 0.111 | 0.274 | 0.587 | 0.591 | 0.649 | 0.544 | 0.554 | 0.068 | 0.601 | 0.18 | 0.685 | 0.064 | 0.512 | 0.62 | 0.544 | 0.539 | 0.649 | **0.007** |
| *IR56a* | 0.0001 | 0.549 | 0.532 | 0.519 | 0.519 | 0.527 | 0.574 | 0.532 | 0.534 | 0.559 | **0** | 0.519 | 0.519 | **0.006** | **0.002** | 0.685 | **0.01** | 0.512 | 0.62 | 0.515 | 0.514 | 0.559 | 0.559 |
| *IR60bcdfo* | 0.04565 | 0.078 | 0.587 | 0.082 | **0.029** | 0.574 | 0.677 | 0.092 | **0.035** | 0.559 | 0.515 | 0.519 | 0.519 | 0.537 | 0.554 | 0.576 | 0.062 | 0.504 | 0.546 | 0.515 | 0.514 | 0.559 | 0.559 |
| ***IR67a*** | 0.0693 | 0.549 | 0.532 | 0.519 | **0.01** | 0.527 | 0.574 | 0.532 | 0.534 | 0.559 | 0.515 | 0.537 | 0.537 | 0.571 | **0.03** | **0.017** | 0.541 | 0.504 | 0.546 | 0.515 | 0.514 | 0.559 | 0.559 |
| *IR76a* | 0.0083 | 0.549 | 0.532 | 0.519 | 0.519 | 0.527 | 0.574 | 0.532 | 0.534 | 0.559 | **0.023** | **0.01** | **0.046** | 0.571 | 0.601 | 0.637 | 0.117 | 0.508 | **0.049** | 0.515 | 0.514 | 0.559 | 0.559 |
| *IR7d* | 0.01955 | 0.549 | 0.532 | 0.519 | 0.519 | 0.527 | 0.574 | 0.532 | 0.534 | **0.002** | 0.515 | 0.519 | 0.519 | 0.537 | 0.554 | 0.576 | 0.541 | 0.504 | **0** | 0.515 | 0.514 | 0.559 | 0.559 |
| *IR7e* | 0.0017 | 0.549 | 0.532 | 0.519 | 0.519 | 0.527 | 0.574 | 0.532 | 0.534 | **0** | 0.515 | 0.519 | 0.519 | 0.537 | 0.554 | 0.576 | 0.062 | 0.504 | 0.069 | 0.515 | **0.007** | 0.5 | 0.5 |
| *IR94d* | 0.0478 | 0.549 | 0.532 | 0.519 | 0.519 | 0.527 | 0.574 | 0.532 | 0.534 | 0.089 | 0.515 | 0.519 | **0.023** | 0.537 | 0.066 | 0.576 | 0.541 | 0.504 | 0.546 | 0.515 | 0.514 | 0.089 | 0.559 |
| ***Obp18a*** | 0.0144 | 0.543 | 0.528 | **0.025** | 0.517 | 0.523 | 0.096 | 0.528 | 0.529 | 0.552 | 0.513 | 0.5 | 0.5 | 0.5 | 0.5 | **0.032** | 0.536 | 0.503 | 0.54 | 0.513 | 0.512 | 0.552 | 0.077 |
| *Obp22a* | 0.0204 | 0.064 | 0.528 | **0.008** | 0.517 | 0.523 | 0.096 | 0.528 | **0.044** | 0.5 | 0.5 | 0.5 | 0.5 | 0.5 | 0.5 | 0.5 | 0.5 | 0.5 | 0.5 | 0.5 | 0.5 | 0.5 | 0.5 |
| *Obp51a56fi* | 0.0086 | **0.044** | 0.553 | 0.533 | **0.017** | **0.012** | 0.096 | 0.528 | **0.044** | 0.5 | 0.5 | 0.5 | 0.5 | 0.5 | 0.5 | 0.5 | 0.5 | 0.5 | 0.5 | 0.5 | 0.5 | 0.5 | 0.5 |
| *Obp57ab* | 0.0886 | 0.543 | **0.014** | 0.533 | 0.533 | 0.523 | 0.096 | 0.528 | **0.044** | 0.5 | 0.5 | 0.5 | 0.5 | 0.5 | 0.5 | 0.5 | 0.5 | 0.5 | 0.5 | 0.5 | 0.5 | 0.5 | 0.5 |

| | | | | | | | | | | | | | | | | | | | | | | | |
|---|---|---|---|---|---|---|---|---|---|---|---|---|---|---|---|---|---|---|---|---|---|---|---|
| ***Obp58b*** | 0.0192 | 0.543 | 0.528 | 0.517 | 0.517 | 0.523 | 0.565 | 0.528 | 0.529 | **0** | 0.513 | 0.5 | 0.5 | 0.5 | 0.5 | **0.032** | 0.536 | 0.503 | 0.54 | 0.513 | 0.512 | 0.552 | 0.552 |
| ***Obp58c*** | 0.00555 | 0.543 | 0.528 | 0.517 | 0.517 | 0.523 | 0.565 | 0.528 | 0.529 | **0** | 0.513 | 0.5 | 0.5 | 0.5 | 0.5 | **0.032** | 0.536 | 0.503 | 0.54 | 0.513 | 0.512 | 0.552 | 0.552 |
| *Obp99b* | 0.04745 | **0.001** | 0.528 | **0.025** | 0.517 | 0.523 | 0.565 | 0.528 | 0.529 | 0.552 | 0.513 | 0.505 | 0.505 | 0.51 | 0.515 | 0.522 | 0.536 | 0.503 | 0.54 | 0.513 | 0.512 | 0.552 | 0.552 |
| ***Or22ab*** | 0.01125 | **0.003** | 0.579 | **0.018** | 0.549 | 0.567 | 0.666 | 0.056 | 0.544 | 0.107 | 0.52 | 0.5 | 0.5 | 0.5 | 0.5 | **0.043** | 0.072 | 0.505 | 0.56 | 0.52 | 0.518 | 0.577 | 0.577 |
| *Or42b* | 0.02145 | 0.565 | 0.542 | 0.525 | 0.525 | 0.535 | 0.595 | 0.542 | 0.544 | **0** | **0.007** | 0.5 | 0.5 | 0.5 | 0.5 | 0.5 | 0.5 | 0.5 | 0.5 | 0.52 | 0.518 | 0.577 | 0.577 |
| *Or59a* | 0.0316 | 0.565 | 0.542 | 0.525 | 0.525 | 0.535 | 0.595 | 0.542 | 0.06 | 0.687 | 0.558 | 0.521 | 0.521 | **0.003** | 0.651 | 0.696 | 0.192 | 0.515 | 0.652 | 0.053 | 0.535 | 0.074 | 0.195 |
| *Or59bORN2* | 0.00065 | 0.565 | 0.542 | 0.525 | 0.525 | 0.535 | 0.595 | **0.034** | 0.113 | 0.687 | **0.003** | **0.045** | 0.594 | 0.663 | 0.322 | 0.765 | **0.045** | 0.525 | **0.018** | 0.558 | 0.551 | **0.012** | 0.687 |
| ***Or65abc*** | 0.01435 | 0.661 | 0.612 | 0.571 | 0.571 | 0.595 | 0.06 | 0.107 | **0.036** | **0.035** | 0.52 | 0.5 | 0.5 | 0.5 | 0.5 | **0.043** | 0.554 | 0.505 | **0.025** | 0.52 | 0.518 | 0.577 | **0.035** |
| *Or98a85a* | 0.012 | 0.661 | 0.612 | 0.571 | **0.028** | 0.595 | 0.719 | 0.612 | 0.618 | 0.114 | 0.077 | 0.61 | 0.156 | **0.017** | 0.686 | 0.734 | 0.671 | 0.52 | **0.039** | 0.558 | 0.551 | **0.04** | 0.687 |
| *Or98b47a* | 0.08915 | 0.618 | 0.579 | 0.549 | 0.549 | 0.567 | 0.666 | 0.056 | 0.544 | **0.035** | 0.52 | 0.541 | 0.541 | 0.058 | 0.56 | 0.583 | 0.072 | 0.505 | 0.56 | 0.52 | 0.518 | **0.035** | 0.577 |
| *ppk10* | 0.0844 | 0.511 | 0.507 | 0.504 | 0.504 | 0.506 | 0.517 | 0.507 | 0.507 | 0.513 | 0.503 | 0.509 | 0.509 | 0.518 | 0.527 | 0.538 | **0.002** | 0.501 | 0.51 | 0.503 | 0.503 | 0.513 | 0.513 |
| *ppk29* | 0.0563 | 0.511 | 0.507 | 0.504 | 0.504 | 0.506 | **0.021** | 0.507 | 0.507 | 0.513 | 0.503 | 0.509 | 0.509 | 0.518 | 0.527 | 0.538 | 0.509 | 0.501 | 0.51 | 0.503 | 0.503 | 0.513 | **0.016** |
| *ppk8* | 0.00815 | 0.511 | 0.507 | 0.504 | 0.504 | 0.506 | 0.517 | 0.507 | 0.507 | **0.016** | 0.503 | 0.5 | 0.5 | 0.5 | 0.5 | 0.056 | 0.509 | 0.501 | 0.51 | 0.503 | 0.503 | 0.513 | **0.016** |
| *Trp01* | 0.01 | 0.517 | 0.511 | 0.506 | 0.506 | 0.509 | 0.526 | 0.511 | 0.511 | 0.521 | 0.505 | **0.003** | 0.502 | 0.504 | **0.009** | 0.508 | 0.514 | 0.501 | 0.516 | 0.505 | 0.504 | 0.521 | 0.521 |
| *Trp02* | 0.0746 | 0.517 | 0.511 | 0.506 | 0.506 | 0.509 | 0.526 | 0.511 | 0.511 | 0.521 | 0.505 | **0.003** | 0.502 | 0.504 | 0.506 | 0.508 | 0.514 | 0.501 | 0.516 | 0.505 | 0.504 | 0.521 | 0.521 |
| *Trp06* | 0.071375 | 0.517 | 0.511 | 0.506 | 0.506 | 0.509 | **0** | 0.511 | 0.511 | 0.521 | 0.505 | 0.502 | 0.502 | 0.504 | 0.506 | 0.508 | 0.514 | 0.501 | 0.516 | 0.505 | 0.504 | 0.521 | 0.521 |
| Cluster7 (*Hsp70*) | 0.00015 | 0.758 | 0.697 | 0.635 | 0.083 | 0.095 | **0.008** | 0.674 | 0.681 | 0.204 | **0.003** | 0.525 | 0.525 | 0.548 | **0.001** | 0.595 | 0.645 | 0.517 | 0.66 | 0.597 | **0.049** | 0.784 | **0.018** |
| Cluster8 (*AOX*) | 0.0354 | 0.705 | 0.649 | 0.597 | 0.597 | 0.628 | 0.086 | 0.649 | 0.237 | **0** | 0.562 | 0.525 | 0.525 | 0.548 | 0.569 | 0.595 | 0.645 | 0.517 | 0.66 | 0.562 | 0.555 | 0.296 | 0.694 |
| Cluster16 (*FASN*) | 0.01185 | 0.669 | 0.619 | 0.576 | 0.576 | 0.601 | **0.023** | 0.619 | 0.624 | **0** | 0.094 | 0.517 | 0.517 | 0.533 | 0.548 | 0.567 | 0.605 | 0.511 | 0.617 | 0.562 | 0.555 | 0.694 | 0.694 |
| Cluster30 (CG15270) | 0.0169 | 0.569 | 0.545 | 0.527 | 0.527 | 0.538 | 0.602 | 0.545 | 0.548 | 0.583 | **0** | 0.533 | **0.017** | 0.562 | 0.589 | 0.621 | 0.679 | 0.522 | 0.695 | 0.522 | 0.519 | 0.583 | 0.583 |
| Cluster32 (CG1943) | 0 | **0** | 0.545 | 0.527 | 0.527 | 0.538 | 0.602 | 0.545 | 0.548 | 0.583 | 0.522 | 0.508 | 0.508 | 0.517 | 0.525 | 0.535 | 0.558 | 0.506 | 0.565 | 0.522 | 0.519 | 0.583 | 0.583 |
| Cluster34 (*retn*) | 0.07575 | 0.569 | 0.545 | 0.527 | 0.527 | 0.538 | 0.602 | 0.545 | 0.548 | 0.583 | **0** | 0.533 | 0.533 | 0.562 | 0.589 | 0.621 | 0.679 | 0.522 | 0.695 | 0.522 | 0.519 | 0.583 | 0.583 |
| Cluster36 (*ATP8A*) | 0.07575 | 0.569 | 0.545 | 0.527 | 0.527 | 0.538 | 0.602 | 0.545 | 0.548 | 0.583 | **0** | 0.533 | 0.533 | 0.562 | 0.589 | 0.621 | 0.679 | 0.522 | 0.695 | 0.522 | 0.519 | 0.583 | 0.583 |
| Cluster58 (*Shrm*) | 0.07435 | 0.569 | 0.545 | 0.527 | 0.527 | 0.538 | 0.602 | 0.545 | 0.548 | 0.583 | **0** | 0.525 | **0.013** | 0.548 | 0.569 | 0.595 | 0.645 | 0.517 | 0.66 | 0.522 | 0.519 | 0.583 | 0.583 |
| Cluster87 (CG12896) | 0.06 | 0.072 | 0.585 | 0.553 | **0.028** | 0.571 | 0.261 | 0.585 | 0.589 | 0.086 | 0.543 | 0.508 | 0.508 | **0.049** | 0.548 | 0.567 | 0.605 | 0.511 | 0.617 | 0.543 | 0.057 | 0.583 | 0.583 |

| | | | | | | | | | | | | | | | | | | | | | | | |
|---|---|---|---|---|---|---|---|---|---|---|---|---|---|---|---|---|---|---|---|---|---|---|---|
| Cluster104 (*CecC*) | 0.0063 | 0.187 | 0.585 | 0.553 | **0.028** | 0.571 | 0.261 | 0.585 | 0.589 | **0.017** | 0.064 | 0.508 | 0.508 | 0.517 | 0.525 | 0.535 | 0.558 | 0.506 | 0.565 | 0.543 | 0.538 | 0.219 | **0.017** |
| Cluster105 (*Or42a*) | 0.06555 | 0.569 | 0.545 | 0.527 | 0.527 | 0.107 | 0.674 | 0.585 | 0.589 | 0.219 | 0.543 | 0.517 | **0.009** | 0.533 | 0.072 | 0.567 | 0.059 | 0.511 | 0.617 | 0.543 | 0.538 | 0.646 | 0.646 |
| Cluster122 (*Cyp4p1*) | 0.0771 | 0.625 | 0.585 | 0.079 | 0.553 | 0.571 | 0.107 | 0.585 | 0.589 | **0.017** | 0.064 | 0.508 | 0.508 | 0.517 | 0.525 | 0.535 | 0.558 | 0.506 | 0.565 | 0.543 | 0.538 | 0.219 | 0.646 |
| Cluster140 (*qless*) | 0.0201 | 0.569 | 0.545 | 0.527 | 0.527 | 0.538 | 0.602 | 0.545 | 0.548 | 0.583 | **0** | **0.025** | 0.517 | 0.072 | 0.569 | 0.595 | 0.645 | 0.517 | 0.66 | 0.522 | 0.519 | 0.583 | 0.583 |
| Cluster161 (FBgn0052473) | 0.03065 | 0.569 | 0.545 | 0.527 | **0** | 0.538 | 0.602 | 0.127 | 0.589 | **0.017** | 0.064 | 0.508 | 0.508 | 0.517 | 0.525 | 0.535 | 0.558 | 0.506 | 0.565 | 0.543 | 0.538 | 0.646 | 0.646 |
| Cluster171 (*BTBD9*) | 0.06885 | 0.569 | 0.545 | 0.527 | 0.527 | 0.538 | **0** | 0.545 | 0.548 | 0.583 | 0.522 | 0.508 | 0.508 | 0.517 | 0.525 | 0.535 | 0.558 | 0.506 | 0.565 | 0.522 | 0.519 | 0.583 | 0.583 |

**Table S7. Counts of genes that have been duplicated or lost in all herbivorous *Scaptomyza*.** Counts are also indicated by cell shading. Herbivores are indicated by green font.

| | Gene | *Dmel* | *Dere* | *Dana* | *Dpse* | *Dvir* | *Dmoj* | *Dgri* | *Spal* | *Shsu* | *Sgra* | *Smon* | *Sfla* | |
|---|---|---|---|---|---|---|---|---|---|---|---|---|---|---|
| Gr | *Gr39aA* | 1 | 0 | 1 | 3 | 6 | 5 | 4 | 9 | 9 | 7 | 4 | 6 | loss |
| | *Gr39aE* | 1 | 1 | 1 | 2 | 0 | 1 | 1 | 1 | 1 | 0 | 0 | 0 | loss |
| | *Gr59ab* | 2 | 3 | 4 | 4 | 2 | 1 | 4 | 4 | 3 | 2 | 2 | 2 | loss |
| | *Gr59cd* | 2 | 2 | 2 | 2 | 4 | 3 | 7 | 9 | 7 | 3 | 4 | 4 | loss |
| | *Gr68a* | 1 | 1 | 1 | 1 | 1 | 1 | 1 | 1 | 1 | 0 | 0 | 0 | loss |
| Gst | *GstS1* | 1 | 1 | 1 | 1 | 1 | 1 | 1 | 1 | 1 | 2 | 2 | 2 | gain |
| Ir | *Ir47abc94abc* | 4 | 3 | 6 | 3 | 5 | 4 | 2 | 1 | 3 | 1 | 1 | 1 | loss |
| | *Ir51be* | 1 | 2 | 2 | 2 | 1 | 1 | 1 | 1 | 1 | 0 | 0 | 0 | loss |
| | *Ir56e* | 0 | 0 | 0 | 0 | 1 | 1 | 1 | 2 | 2 | 1 | 1 | 1 | loss |
| | *Ir60e* | 1 | 1 | 1 | 1 | 0 | 1 | 1 | 1 | 1 | 0 | 0 | 0 | loss |
| | *Ir67a* | 0 | 1 | 1 | 1 | 1 | 1 | 1 | 1 | 1 | 2 | 2 | 3 | gain |
| | *Ir7f* | 1 | 1 | 1 | 1 | 1 | 1 | 1 | 1 | 1 | 0 | 0 | 0 | loss |
| | *Ir94f* | 1 | 1 | 1 | 0 | 1 | 1 | 1 | 1 | 1 | 0 | 0 | 0 | loss |
| Obp | *Obp18a* | 1 | 0 | 1 | 0 | 0 | 1 | 1 | 1 | 1 | 0 | 0 | 0 | loss |
| | *Obp46a* | 1 | 1 | 1 | 1 | 1 | 1 | 1 | 1 | 1 | 0 | 0 | 0 | loss |
| | *Obp50cd* | 2 | 2 | 2 | 2 | 1 | 1 | 1 | 1 | 1 | 0 | 0 | 0 | loss |
| | *Obp56b* | 1 | 1 | 1 | 1 | 1 | 1 | 1 | 1 | 1 | 0 | 0 | 0 | loss |
| | *Obp58b* | 1 | 1 | 1 | 1 | 1 | 1 | 4 | 1 | 1 | 0 | 0 | 0 | loss |
| | *Obp58c* | 1 | 1 | 1 | 1 | 1 | 1 | 5 | 1 | 1 | 0 | 0 | 0 | loss |
| | *Obp58d* | 1 | 1 | 1 | 1 | 1 | 1 | 1 | 1 | 1 | 0 | 0 | 0 | loss |
| | *Obp93a* | 1 | 1 | 1 | 1 | 1 | 1 | 1 | 1 | 1 | 0 | 0 | 0 | loss |
| Or | *Or22a* | 2 | 1 | 5 | 2 | 1 | 1 | 2 | 2 | 1 | 0 | 0 | 0 | loss |
| P450 | *Cyp4ad1* | 1 | 1 | 1 | 1 | 1 | 1 | 1 | 1 | 1 | 0 | 0 | 0 | loss |
| | *Cyp4d1* | 1 | 1 | 2 | 1 | 1 | 2 | 1 | 1 | 1 | 0 | 0 | 0 | loss |
| Ppk | *ppk8* | 1 | 1 | 1 | 1 | 0 | 1 | 0 | 1 | 1 | 0 | 0 | 0 | loss |
| Ugt | *Ugt302e1* | 1 | 1 | 0 | 1 | 1 | 1 | 2 | 1 | 1 | 0 | 0 | 0 | loss |

**Table S8. PAML analyses under branch and branch-site models.** Shown are models in which there was a significant difference in selective constraint between the ancestral herbivore branch and background branches. Parentheses surrounding genes indicate that the branch at the base of these genes was evaluated (i.e. genes with multiple paralogs wherein only some paralogs experienced significant shifts in selection). Values in parenthesis to the right of dN/dS values indicate proportion of sites corresponding to the given dN/dS rate.

| Gene | Model | κ | tree length | dN/dS | | | | | | | | lnL | LRT | *q*-value *(FDR 5%)* |
|---|---|---|---|---|---|---|---|---|---|---|---|---|---|---|
| Csp2 | M0 (one ratio) | 1.4 | 5.3 | ω= | 0.07 | | | | | | | -1897.9 | 7.14 | 0.05 |
| | Branch (two-ratios) | 1.4 | 5.5 | ω0= | 0.06 | ω1= | 0.24 | | | | | -1894.3 | | |
| Gr59e | M0 (one ratio) | 2.1 | 6.7 | ω= | 0.26 | | | | | | | -6683.9 | 18.26 | <0.001 |
| | Branch (two-ratios) | 2.1 | 7 | ω0= | 0.23 | ω1= | 999 | | | | | -6674.7 | | |
| Gr63a | Branch-site Model A | 1.8 | 3.6 | ω0= | 0.03 (81%) | ω1= | 1 (7%) | ω2a= | 1 (11%) | ω2b= | 1 (1%) | -6110.5 | 16.52 | <0.01 |
| | Branch-site Model A (ω = 1) | 1.8 | 3.6 | ω0= | 0.03 (0%) | ω1= | 1 (0%) | ω2a= | 1 (92%) | ω2b= | 1 (8%) | -6118.7 | | |
| Gr98a | M0 (one ratio) | 1.9 | 16.3 | ω= | 0.29 | | | | | | | -19802.4 | 7.74 | 0.04 |
| (SmonGr98a1, SflaGr98a1, SgraGr98a2) | Branch (two-ratios) | 1.9 | 16.3 | ω0= | 0.29 | ω1= | 0.87 | | | | | -19798.5 | | |
| | Branch-site Model A | 2 | 17 | ω0= | 0.22 (71%) | ω1= | 1 (26%) | ω2a= | 32.44 (2%) | ω2b= | 32.44 (1%) | -19593 | 11.4 | 0.03 |
| | Branch-site Model A (ω = 1) | 2 | 16.9 | ω0= | 0.22 (65%) | ω1= | 1 (24%) | ω2a= | 1 (8%) | ω2b= | 1 (3%) | -19598.7 | | |
| Gr98bcd | Branch-site Model A | 2.1 | 23.6 | ω0= | 0.16 (73%) | ω1= | 1 (27%) | ω2a | 179.55 (0.004%) | ω2b= | 179.55 (0.001%) | -12238.2 | 10.2 | 0.05 |
| | Branch-site Model A (ω = 1) | 2.1 | 23.5 | ω0= | 0.16 (73%) | ω1= | 1 (27%) | ω2a | 1 (0%) | ω2b= | 1 (0%) | -12243.3 | | |
| GstE9 | M0 (one ratio) | 1.6 | 4.1 | ω= | 0.14 | | | | | | | -3640.7 | 9.35 | 0.02 |
| | Branch (two-ratios) | 1.6 | 4.1 | ω0= | 0.15 | ω1= | 0.01 | | | | | -3636 | | |
| GstO2 | M0 (one ratio) | 1.8 | 5.5 | ω= | 0.11 | | | | | | | -4112.2 | 10.3 | 0.01 |
| | Branch (two-ratios) | 1.8 | 5.4 | ω0= | 0.12 | ω1= | 0 | | | | | -4107.1 | | |
| GstS1 | M0 (one ratio) | 2 | 1.9 | ω= | 0.11 | | | | | | | -2833 | 50.43 | <0.0001 |

| | | | | | | | | | | | | | | |
|---|---|---|---|---|---|---|---|---|---|---|---|---|---|---|
| | Branch (two-ratios) | 2 | 1.9 | ω0= | 0.09 | ω1= | 3.27 | | | | | -2807.8 | | |
| (SflaGstS1b, SmonGstS1b, SgraGstS1b) | Branch-site Model A | 2.1 | 1.9 | ω0= | 0.08 (84%) | ω1= | 1 (2%) | ω2a= | 38.47 (13%) | ω2b= | 38.47 (0.004%) | -2800.91 | 14.04 | 0.01 |
| | Branch-site Model A (ω = 1) | 2 | 1.9 | ω0= | 0.07 (0%) | ω1= | 1 (0%) | ω2a= | 1 (97%) | ω2b= | 1 (3%) | -2807.93 | | |
| Ir21a | M0 (one ratio) | 1.5 | 4.7 | ω= | 0.11 | | | | | | | -13380.2 | 9.2 | 0.02 |
| | Branch (two-ratios) | 1.5 | 4.8 | ω0= | 0.11 | ω1= | 0.26 | | | | | -13375.6 | | |
| Ir48d | M0 (one ratio) | 1.9 | 3.5 | ω= | 0.19 | | | | | | | -8419.1 | 7.96 | 0.04 |
| | Branch (two-ratios) | 1.9 | 3.5 | ω0= | 0.18 | ω1= | 0.38 | | | | | -8415.2 | | |
| Ir56a | M0 (one ratio) | 1.8 | 15 | ω= | 0.34 | | | | | | | -23071.9 | 18.8 | <0.001 |
| (SflaIr56a, SmonIr56a) | Branch (two-ratios) | 1.8 | 15.1 | ω0= | 0.33 | ω1= | 0.98 | | | | | -23062.5 | | |
| (SgraIr56a1, SgraIr56a2, SgraIr56a4, SgraIr56a5) | Branch (two-ratios) | 1.8 | 15 | ω0= | 0.33 | ω1= | 3.66 | | | | | -23067.9 | 7.98 | 0.03 |
| | Branch (two-ratios, fixed ω1 = 1) | 1.8 | 15 | ω0= | 0.34 | ω1= | 1 | | | | | -23068.7 | 1.56 | 0.31 |
| Ir60a | Branch-site Model A | 1.8 | 7.1 | ω0= | 0.06 (83%) | ω1= | 1 (16%) | ω2a= | 999 (0.005%) | ω2b= | 999 (0.001%) | -10580.9 | 10 | 0.05 |
| | Branch-site Model A (ω = 1) | 1.8 | 5.4 | ω0= | 0.06 (82%) | ω1= | 1 (16%) | ω2a= | 1 (2%) | ω2b= | 1 (0.003%) | -10585.9 | | |
| Ir67a | M0 (one ratio) | 1.8 | 4 | ω= | 0.31 | | | | | | | -10318.5 | 10.6 | 0.01 |
| | Branch (two-ratios) | 1.8 | 4 | ω0= | 0.3 | ω1= | 0.6 | | | | | -10313.2 | | |
| Obp57cL1 | Branch-site Model A | 2.1 | 9.3 | ω0= | 0.21 (59%) | ω1= | 1 (36%) | ω2a= | 148.51 (3%) | ω2b= | 148.51 (2%) | -2539.7 | 15.68 | <0.01 |
| | Branch-site Model A (ω = 1) | 2 | 7.3 | ω0= | 0.21 (55%) | ω1= | 1 (30%) | ω2a= | 1 (10%) | ω2b= | 1 (5%) | -2547.5 | | |
| Or19a | M0 (one ratio) | 1.7 | 7.6 | ω= | 0.27 | | | | | | | -7966.9 | 13.7 | <0.01 |
| | Branch (two-ratios) | 1.7 | 7.5 | ω0= | 0.3 | ω1= | 0.09 | | | | | -7960.1 | | |
| Or22c | M0 (one ratio) | 1.6 | 3.7 | ω= | 0.17 | | | | | | | -5823.7 | 7.24 | 0.05 |
| | Branch (two-ratios) | 1.6 | 3.7 | ω0= | 0.18 | ω1= | 0.05 | | | | | -5820.1 | | |

| | | | | | | | | | | | | | | |
|---|---|---|---|---|---|---|---|---|---|---|---|---|---|---|
| Or42a | M0 (one ratio) | 1.7 | 7.5 | ω= | 0.14 | | | | | | | -10294.1 | 11.46 | <0.01 |
| (SflaOr42a1, SmonOr42a1, SgraOr42a1) | Branch (two-ratios) | 1.7 | 7.6 | ω0= | 0.13 | ω1= | 0.45 | | | | | -10288.4 | | |
| Or56a | M0 (one ratio) | 1.5 | 4.3 | ω= | 0.1 | | | | | | | -6120.1 | 8.98 | 0.02 |
| | Branch (two-ratios) | 1.5 | 4.3 | ω0= | 0.11 | ω1= | 0.03 | | | | | -6115.6 | | |
| Or63a | M0 (one ratio) | 1.6 | 4.7 | ω= | 0.22 | | | | | | | -7544.9 | 29.5 | <0.0001 |
| | Branch (two-ratios) | 1.6 | 4.8 | ω0= | 0.2 | ω1= | 0.76 | | | | | -7530.2 | | |
| Or67d | M0 (one ratio) | 1.9 | 4.7 | ω= | 0.16 | | | | | | | -4995.1 | 15.18 | <0.01 |
| | Branch (two-ratios) | 1.9 | 4.5 | ω0= | 0.18 | ω1= | 0.03 | | | | | -4987.5 | | |
| Or85aLike | M0 (one ratio) | 1.5 | 3.9 | ω= | 0.14 | | | | | | | -5773.6 | 6.98 | 0.05 |
| | Branch (two-ratios) | 1.5 | 4 | ω0= | 0.13 | ω1= | 0.29 | | | | | -5770.1 | | |
| Or85f | Branch-site Model A | 2.1 | 5.5 | ω0= | 0.13 (88%) | ω1= | 1 (11%) | ω2a= | 46.61 (1%) | ω2b= | 46.61 (0.001%) | -5725.5 | 12.54 | 0.03 |
| | Branch-site Model A (ω = 1) | 2.1 | 5.4 | ω0= | 0.13 (85%) | ω1= | 1 (10%) | ω2a= | 1 (5%) | ω2b= | 1 (1%) | -5731.8 | | |
| Or98aLike1 | M0 (one ratio) | 1.6 | 2.9 | ω= | 0.18 | | | | | | | -5564.4 | 7.58 | 0.04 |
| | Branch (two-ratios) | 1.6 | 2.9 | ω0= | 0.19 | ω1= | 0.05 | | | | | -5560.6 | | |
| Or98aLike2 | M0 (one ratio) | 1.9 | 4.3 | ω= | 0.18 | | | | | | | -5970.6 | 35.92 | <0.0001 |
| | Branch (two-ratios) | 1.9 | 4.2 | ω0= | 0.2 | ω1= | 0 | | | | | -5952.7 | | |
| OrN2.3prime | Branch-site Model A | 1.9 | 9.2 | ω0= | 0.16 (73%) | ω1= | 1 (20%) | ω2a= | 15.3 (5%) | ω2b= | 15.3 (1%) | -11146.2 | 15.6 | <0.01 |
| | Branch-site Model A (ω = 1) | 1.9 | 9.2 | ω0= | 0.16 (60%) | ω1= | 1 (17%) | ω2a= | 1 (18%) | ω2b= | 1 (5%) | -11154 | | |
| Cyp28a5 | M0 (one ratio) | 1.6 | 4.6 | ω= | 0.12 | | | | | | | -8110 | 9.95 | 0.02 |
| | Branch (two-ratios) | 1.6 | 4.5 | ω0= | 0.13 | ω1= | 0.05 | | | | | -8105.1 | | |
| Cyp310a1 | M0 (one ratio) | 1.9 | 4.4 | ω= | 0.21 | | | | | | | -8063.8 | 18.43 | <0.001 |
| | Branch (two-ratios) | 1.9 | 4.4 | ω0= | 0.19 | ω1= | 0.57 | | | | | -8054.6 | | |
| Cyp4d14 | Branch-site Model A | 1.8 | 7.2 | ω0= | 0.06 (84%) | ω1= | 1 (15%) | ω2a= | 809.22 (1%) | ω2b= | 809.22 (0.001%) | -7537.6 | 11.44 | 0.03 |

| | | | | | | | | | | | | | | |
|---|---|---|---|---|---|---|---|---|---|---|---|---|---|---|
| | Branch-site Model A (ω = 1) | 1.8 | 5.3 | ω0= | 0.06 (81%) | ω1= | 1 (15%) | ω2a= | 1 (4%) | ω2b= | 1 (1%) | -7543.3 | | |
| Cyp6a16 | M0 (one ratio) | 1.7 | 3.5 | ω= | 0.2 | | | | | | | -8081 | 7.57 | 0.04 |
| | Branch (two-ratios) | 1.7 | 3.5 | ω0= | 0.21 | ω1= | 0.1 | | | | | -8077.2 | | |
| Cyp6a22 | M0 (one ratio) | 1.6 | 3.4 | ω= | 0.09 | | | | | | | -5558.1 | 21.17 | <0.001 |
| | Branch (two-ratios) | 1.6 | 3.5 | ω0= | 0.07 | ω1= | 0.26 | | | | | -5547.5 | | |
| Cyp6u1 | M0 (one ratio) | 1.5 | 5.2 | ω= | 0.15 | | | | | | | -8378.5 | 20.06 | <0.001 |
| | Branch (two-ratios) | 1.6 | 5.3 | ω0= | 0.14 | ω1= | 0.61 | | | | | -8368.5 | | |
| Ppk10 | M0 (one ratio) | 1.5 | 5.4 | ω= | 0.1 | | | | | | | -7191.5 | 12.12 | <0.01 |
| | Branch (two-ratios) | 1.5 | 5.3 | ω0= | 0.1 | ω1= | 0.01 | | | | | -7185.4 | | |
| Ppk12 | M0 (one ratio) | 1.5 | 5.6 | ω= | 0.19 | | | | | | | -10244.3 | 10.07 | 0.02 |
| | Branch (two-ratios) | 1.5 | 5.6 | ω0= | 0.18 | ω1= | 0.38 | | | | | -10239.3 | | |
| Ppk13 | M0 (one ratio) | 1.5 | 3.2 | ω= | 0.06 | | | | | | | -5756.3 | 16.12 | <0.01 |
| | Branch (two-ratios) | 1.5 | 3.3 | ω0= | 0.05 | ω1= | 0.19 | | | | | -5748.3 | | |
| Ppk19 | Branch-site Model A | 1.7 | 6.1 | ω0= | 0.11 (75%) | ω1= | 1 (23%) | ω2a= | 15.75 (2%) | ω2b= | 15.75 (1%) | -8909.7 | 13.98 | 0.01 |
| | Branch-site Model A (ω = 1) | 1.7 | 6 | ω0= | 0.11 (72%) | ω1= | 1 (23%) | ω2a= | 1 (4%) | ω2b= | 1 (1%) | -8916.7 | | |
| Ppk22 | M0 (one ratio) | 1.7 | 4.5 | ω= | 0.16 | | | | | | | -9151.1 | 7.7 | 0.04 |
| | Branch (two-ratios) | 1.7 | 4.5 | ω0= | 0.16 | ω1= | 0.37 | | | | | -9147.3 | | |
| Ppk25 | M0 (one ratio) | 1.5 | 6.1 | ω= | 0.2 | | | | | | | -8564.4 | 16.82 | <0.01 |
| | Branch (two-ratios) | 1.5 | 6 | ω0= | 0.21 | ω1= | 0.04 | | | | | -8556 | | |
| Ppk28 | M0 (one ratio) | 1.5 | 3.4 | ω= | 0.1 | | | | | | | -7869.2 | 11.72 | <0.01 |
| | Branch (two-ratios) | 1.5 | 3.3 | ω0= | 0.1 | ω1= | 0.03 | | | | | -7863.4 | | |
| Ppk30 | M0 (one ratio) | 1.5 | 6.1 | ω= | 0.22 | | | | | | | -8365 | 12.2 | <0.01 |
| | Branch (two-ratios) | 1.5 | 6.3 | ω0= | 0.2 | ω1= | 0.44 | | | | | -8358.9 | | |
| Ppk31 | M0 (one ratio) | 1.1 | 4 | ω= | 0.13 | | | | | | | -6663 | 17.23 | <0.01 |
| | Branch (two-ratios) | 1.1 | 4 | ω0= | 0.12 | ω1= | 0.39 | | | | | -6654.4 | | |
| Ppk5 | M0 (one ratio) | 1.5 | 5.4 | ω= | 0.12 | | | | | | | -8397.9 | 15.16 | <0.01 |

| | | | | | | | | | | | | | | |
|---|---|---|---|---|---|---|---|---|---|---|---|---|---|---|
| | Branch (two-ratios) | 1.5 | 5.5 | ω0= | 0.11 | ω1= | 0.36 | | | | | -8390.3 | | |
| Ppk6 | M0 (one ratio) | 1.9 | 3.2 | ω= | 0.1 | | | | | | | -6550 | 10.35 | 0.01 |
| | Branch (two-ratios) | 1.9 | 3.2 | ω0= | 0.09 | ω1= | 0.25 | | | | | -6544.9 | | |
| Ppk7 | M0 (one ratio) | 1.5 | 3.5 | ω= | 0.16 | | | | | | | -8885.9 | 7.75 | 0.04 |
| | Branch (two-ratios) | 1.5 | 3.5 | ω0= | 0.15 | ω1= | 0.27 | | | | | -8882 | | |
| Trp04 | M0 (one ratio) | 1.6 | 2.7 | ω= | 0.03 | | | | | | | -8768.9 | 10.66 | 0.01 |
| | Branch (two-ratios) | 1.6 | 2.7 | ω0= | 0.03 | ω1= | 0.09 | | | | | -8763.6 | | |
| | Branch-site Model A | 1.8 | 3.6 | ω0= | 0.01 (94%) | ω1= | 1 (6%) | ω2a= | 999 (0.004%) | ω2b= | 999 (0.0002%) | -8618 | 12.2 | 0.03 |
| | Branch-site Model A (ω = 1) | 1.8 | 2.9 | ω0= | 0.01 (92%) | ω1= | 1 (6%) | ω2a= | 1 (3%) | ω2b= | 1 (0.002%) | -8624.1 | | |
| Trp12 | Branch-site Model A | 1.9 | 2.1 | ω0= | 0.01 (97%) | ω1= | 1 (3%) | ω2a= | 999 (0.001%) | ω2b= | 999 (0.00003%) | -16143.7 | 11.8 | 0.03 |
| | Branch-site Model A (ω = 1) | 1.9 | 1.9 | ω0= | 0.01 (96%) | ω1= | 1 (3%) | ω2a= | 1 (0.004%) | ω2b= | 1 (0.0001%) | -16149.6 | | |
| Ugt301D1 | M0 (one ratio) | 1.8 | 4.2 | ω= | 0.08 | | | | | | | -7343.2 | 7.22 | 0.05 |
| | Branch (two-ratios) | 1.8 | 4.1 | ω0= | 0.09 | ω1= | 0.03 | | | | | -7339.6 | | |
| Ugt302C1 | M0 (one ratio) | 1.8 | 3.9 | ω= | 0.1 | | | | | | | -7695.6 | 7.42 | 0.05 |
| | Branch (two-ratios) | 1.8 | 3.9 | ω0= | 0.11 | ω1= | 0.04 | | | | | -7691.9 | | |

**Table S9. Repetitive element content (as a proportion of genome size) in *S. flava* falls within the range observed across other *Drosophila* genomes.**

| Species | LINE | SINE | LTR | DNA | Unclassified | SmRNA | Others | Total (%) |
|---|---|---|---|---|---|---|---|---|
| *Drosophila melanogaster* | 8,532 | 39 | 14,673 | 5,964 | 9,873 | 257 | 93,764 | 23.71 |
| *Drosophila simulans* | 8,649 | 25 | 11,204 | 5,413 | 9,369 | 27 | 84,572 | 12.04 |
| *Drosophila pseudoobscura* | 11,054 | 82 | 12,481 | 9,418 | 28,827 | 157 | 191,276 | 21.23 |
| *Drosophila virilis* | 15,786 | 0 | 20,749 | 9,133 | 35,155 | 986 | 231,347 | 31.84 |
| *Drosophila mojavensis* | 16,605 | 856 | 18,226 | 27,355 | 48,632 | 125 | 326,217 | 28.64 |
| *Drosophila yakuba* | 13,533 | 0 | 24,306 | 12,327 | 29,688 | 474 | 108,910 | 27.31 |
| *Drosophila erecta* | 17,007 | 0 | 16,968 | 5,845 | 28,264 | 976 | 91,589 | 23.78 |
| *Drosophila anannasse* | 20,405 | 35 | 44,019 | 36,293 | 67,229 | 768 | 124,103 | 44.14 |
| *Drosophila sechellia* | 21,266 | 64 | 19,213 | 6,222 | 21,766 | 1,063 | 97,717 | 28.7 |
| *Drosophila persimilis* | 16,818 | 88 | 24,630 | 16,653 | 48,215 | 271 | 188,702 | 33.06 |
| *Drosophila grimshawi* | 15,538 | 0 | 36,347 | 8,867 | 36,262 | 105 | 281,598 | 30.56 |
| *Scaptomyza flava* (sfla_v1) | 29,063 | 2,208 | 19,457 | 26,651 | 123,393 | 2,241 | 213,342 | 33.82 |

**Table S10. Gene functions of chemosensory and detoxification genes that were duplicated, lost, or experienced a change in selection regime in the ancestral lineage of herbivorous *Scaptomyza*.** *Indicates that the ancestral herbivore lineage experienced a gene loss or gain, positive selection (dN/dS>1), relaxed purifying selection (foreground branch showed significantly higher dN/dS), or stronger purifying selection (foreground branch showed significantly lower dN/dS) relative to the background branches.

| Gene | Herbivore-specific change* | Gene Function in *D. melanogaster* | References |
|---|---|---|---|
| **Bitter Detection** | | | |
| *Gr39aA* | Loss | Involved in the detection of many bitter compounds. | (Sung *et al.* 2017; Dweck and Carlson 2020) |
| *Gr59a* | Loss | Putative bitter reception: expressed in labellar S-a and S-b type bitter gustatory neurons. | (Weiss *et al.* 2011) |
| *Gr59d* | Loss | Bitter detection in larvae. Expressed in labellar S-a and I-a type bitter gustatory neurons. | (Weiss *et al.* 2011; Kim *et al.* 2016) |
| *Gr98a* | Positive selection | Detects the toxic amino acid histamine, found in high amounts in fermented foods. | (Aryal and Lee 2022) |
| *Gr98bcd* | Positive selection | Gr98b is required for detection of the toxic plant-derived amino acid L-canavanine. | (Shim *et al.* 2015) |
| *Ir56a* | Relaxed purifying | Coexpressed with bitter neurons. | (Koh et al. 2014) |
| **Yeast/fruit volatile detection** | | | |
| *Obp18a* | Loss | Involved in detecting diverse yeast/fruit volatiles (propanol, benzaldehyde, citral, 2-heptanone, isoamylacetate, methyl salicylate, phenyl-ethanol, d-carvone). Down-regulated in *D. sechellia.* | (Dworkin and Jones 2009; Swarup *et al.* 2011) |
| *Obp58b* | Loss | Involved in detecting diverse yeast/fruit volatiles (1-hexanol, 2-ethylpyrazine, d-carvone, isoamylacetate, methyl salicylate). | (Swarup *et al.* 2011) |

| | | | |
|---|---|---|---|
| *Obp58c* | Loss | Involved in detecting diverse yeast/fruit volatiles (1-hexanol, geraniol, 2-heptanone). | (Swarup *et al.* 2011) |
| *Obp93a* | Loss | Involved in detecting diverse yeast/fruit volatiles (1-hexanol, geraniol, ethylpyrazine, benzaldehyde). | (Swarup *et al.* 2011) |
| *Or19a* | Stronger purifying | Valencene (citrus volatile) detection. Exclusively mediates preference for citrus substrates for egg-laying. | (Dweck et al. 2013) |
| *Or22a* | Loss | Detection of esters and alcohols produced during fermentation by yeasts. | (Stensmyr *et al.* 2003) |
| *Or22c* | Stronger purifying | Detection of fruit volatiles in larvae. | (Dweck et al. 2018) |
| *Or85aLike* | Relaxed purifying | *Or85a* detects fatty alcohols produced during fermentation by yeasts. | (Ramasamy et al. 2016) |
| **Other sensory detection** | | | |
| *Gr39aA* | Loss | Mutant males show reduced courtship. | (Watanabe *et al.* 2011) |
| *Gr68a* | Loss | Detection of female sex pheromone. | (Bray and Amrein 2003) |
| *Gr63a* | Positive selection | Carbon dioxide detection. | (Kwon *et al.* 2007) |
| *Ir21a* | Relaxed purifying | Cooling detection and avoidance. | (Ni et al. 2016) |
| *Ppk19* | Positive selection | Detection of high salt concentrations in both larvae and adults. | (Liu *et al.* 2003; Alves *et al.* 2014) |
| *Ppk25* | Stronger purifying | Involved in the detection of female pheromone 7,11-heptacosadiene. | (Liu et al. 2018) |
| *Or42a* | Relaxed purifying | Required for the detection of diverse odors in larvae. | (Fishilevich et al. 2005) |
| *Or56a* | Stronger purifying | Detection of geosmin, emitted by harmful microbes (the only known odorant of *Or56a*). | (Chin et al. 2018) |
| *Or67d* | Stronger purifying | Detection of cVA pheromone. | (Kurtovic et al. 2007) |
| *Or85f* | Positive selection | Detects parasitoid wasp (*Leptopilina*) volatiles. Best ligand is the ketone acetophenone. | (Ebrahim *et al.* 2015) |
| *Or98aLike1* | Stronger purifying | *Or98a* detects a broad range of yeast/fruit volatiles, the pyrethrum component (E)-β-farnesene, and mediates copulation. | (Hallem & Carlson 2006; Wang et al. 2021; Sakurai et al. 2013) |
| *Or98aLike2* | Stronger purifying | *Or98a* detects a broad range of yeast/fruit volatiles, the pyrethrum component (E)-β-farnesene, and mediates copulation. | (Hallem & Carlson 2006; Wang et al. 2021; Sakurai et al. 2013) |
| *Ppk28* | Stronger purifying | Sensing pure water or low osmolarity in taste neurons. | (Cameron et al. 2010) |
| *Ppk30* | Relaxed purifying | Mechanosensing and acid sensing. | (Jang et al. 2019) |
| *nan (Trp4)* | Relaxed purifying | Involved in mechanosensation, hearing, and humidity sensing | (Liu et al. 2007) |
| **Detoxification** | | | |
| *GstE9* | Stronger purifying | Detoxification of 4-hydroxynonenal, a lipid peroxidation product | |
| *GstS1* | Duplication & positive selection | Detoxification of lipid peroxidation product, 4-HN, oxidation product of adrenalin, adrenochrome, and methylmercury. Expressed in CNS and during development, but highest in flight muscle. | (Clayton *et al.* 1998; Singh *et al.* 2001; Saisawang *et al.* 2012; Vorojeikina *et al.* 2017) |
| *Cyp4ad1* | Loss | Up-regulated in response to insecticide pyrethroid deltamethrin. Down-regulated by ecdysteroid agonists. Expressed in larval gonads. | (Davies *et al.* 2006; Liu *et al.* 2020) |
| *Cyp4d1* | Loss | Up-regulated in response to insecticide pyrethroid deltamethrin. Down-regulated by ecdysteroid agonists. Expressed in midgut and fat body. The only P450 that exhibits alternate splicing. | (Davies *et al.* 2006; Chung *et al.* 2009; Liu *et al.* 2020) |
| *Cyp4d14* | Positive selection | Up-regulated in response to phenobarbital, caffeine, mycotoxins. Expressed in larval midgut. Also lost in *D. sechellia.* | (Sun *et al.* 2006; Willoughby *et al.* 2006; Chung *et al.* 2009; Trienens *et al.* 2017; Rane *et al.* 2019) |
| *Cyp28a5* | Stronger purifying | Up-regulated in response to methanol and the insecticide butene-fipronil. | (Wang et al. 2012; Arain et al. 2018) |
| **Other functions** | | | |
| *Ir67a* | Duplication | Wing vein development. | (George *et al.* 2019) |
| *Ppk8* | Loss | Involved in regulating neuronal excitability through interactions with the gene seizure. Expressed in embryonic trachea and adult fat body. | (Xu *et al.* 2011; Yehuda 2012; Suslak 2015) |
| *EbpIII* | Relaxed purifying | Involved in pheromone production. | (Montagne & |

| | | | |
|---|---|---|---|
| *(CSP2)* | | | Wicker-Thomas 2021) |
| *Cyp6u1* | Relaxed purifying | Ecdysteroid metabolism. | (Christesen et al. 2017) |

**Unknown Function in *Drosophila***

| | | | | | | | | | |
|---|---|---|---|---|---|---|---|---|---|
| *Gr39aE* | Loss | | | *Obp58d* | Loss | *Ppk12* | Relaxed purifying | *Cyp310a1* | Relaxed purifying |
| *GstO2* | Relaxed purifying | *Ir94abc* | Loss | *Or63a* | Relaxed purifying | *Ppk13* | Relaxed purifying | *Ugt301d1* | Stronger purifying |
| *Ir7f* | Loss | *Ir94f* | Loss | *OrN2.3prime* | Positive selection | *Ppk22* | Relaxed purifying | *Ugt302c1* | Stronger purifying |
| *Ir48d* | Relaxed purifying | *Obp46a* | Loss | *Ppk5* | Relaxed purifying | *Ppk31* | Relaxed purifying | *Ugt302e1* | Loss |
| *Ir51e* | Loss | *Obp50cd* | Loss | *Ppk6* | Relaxed purifying | *Cyp6a16* | Stronger purifying | | |
| *Ir56e* | Loss | *Obp56b* | Loss | *Ppk7* | Relaxed purifying | *Cyp6a22* | Relaxed purifying | | |
| *Ir60e* | Loss | *Obp57cL1* | Positive selection | *Ppk10* | Stronger purifying | *Cyp12g1* | Loss | | |

### 3. SUPPLEMENTARY FIGURES

**Figure S1. Benchmarking with core dipteran genes.** High levels of completeness are found in (a) the genome assemblies and (b) gene annotations used to curate the chemosensory and detoxification gene families, estimated using BUSCO v.5.4.2 (diptera_odb10, n=3285 genes).

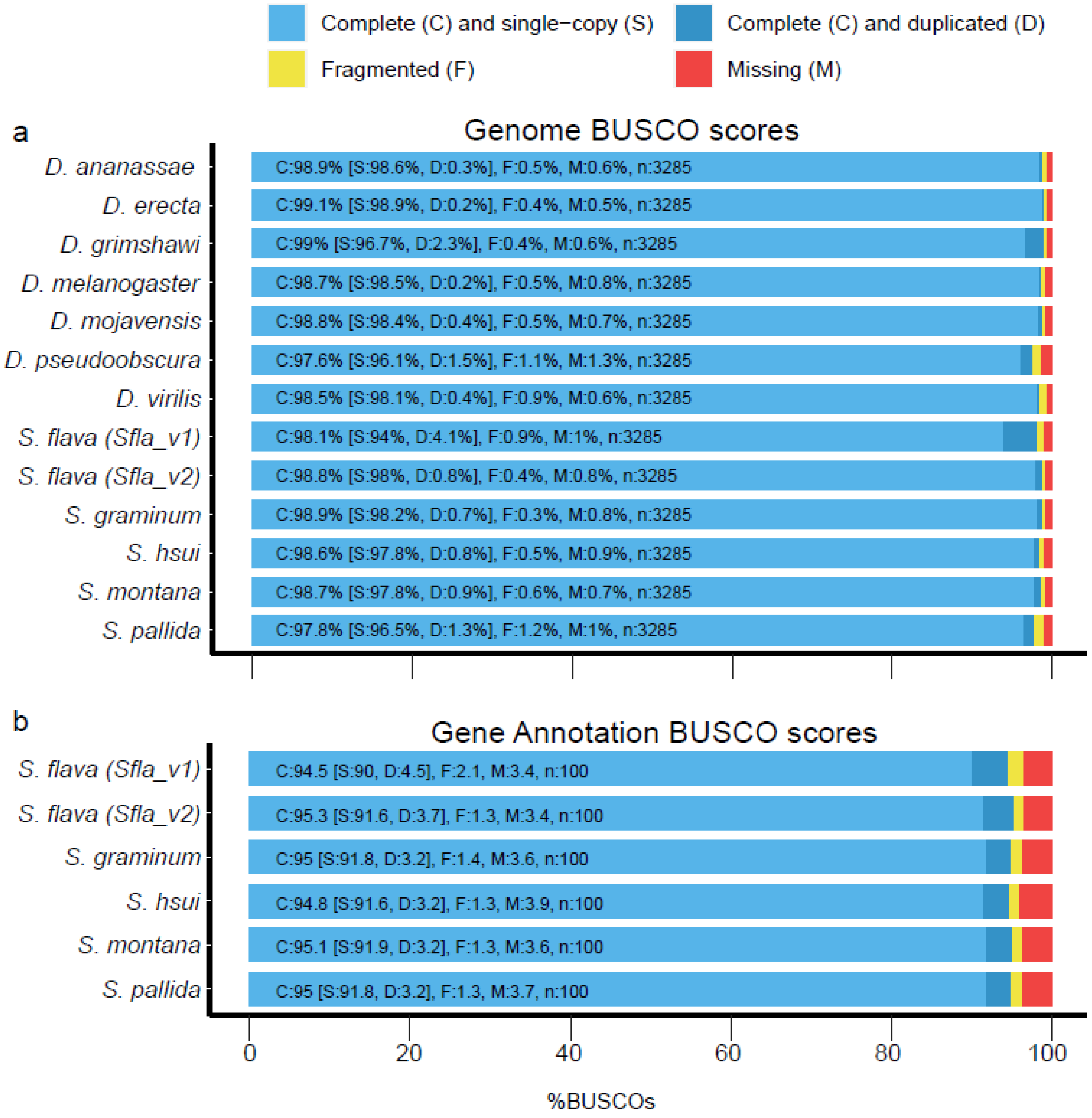

**Figure S2. Phylogeny of gustatory receptors.** A midpoint rooted ML gene tree inferred and visualized using RAxML (Stamatakis 2006) and iTOL (Letunic and Bork 2021). Bootstrap support is given the size of squares at the midpoint of each branch, with only those >70 shown. Species are coded by font color: *D. grimshawi* (red), *S. pallida* (blue), *S. hsui* (magenta), *S. montana* (yellow), *S. flava* (green), *S. graminum* (purple). Gene orthology groups are indicated by alternating branch shading. Outer colored clade labels group genes by known functional classes: sweet/sugar receptors (red), $CO_2$ receptors (turquoise), bitter receptors (green); thermosensitive receptor (purple), and pheromone receptor (orange).

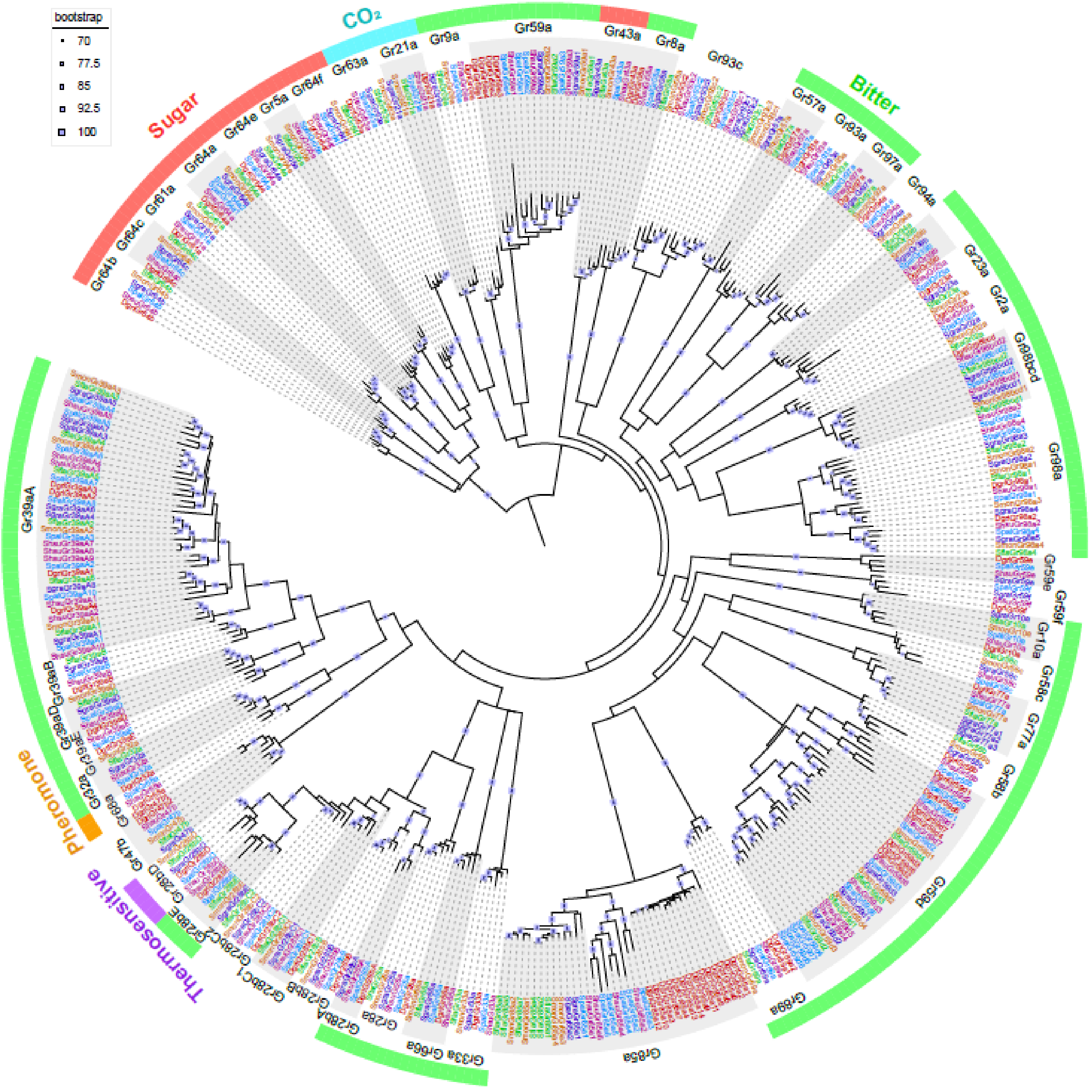

**Figure S3. Phylogeny of ionotropic receptors.** A midpoint rooted ML gene tree inferred and visualized using RAxML (Stamatakis 2006) and iTOL (Letunic and Bork 2021). Bootstrap support is given the size of squares at the midpoint of each branch, with only those >70 shown. Species are coded by font and tip color: *D. grimshawi* (red), *S. pallida* (blue), *S. hsui* (magenta), *S. montana* (yellow), *S. flava* (green), *S. graminum* (purple). Gene orthology groups are indicated by alternating branch shading. Outer colored clade labels group genes by known IR classes: divergent IRs (blue), antennal IRs (yellow), IR co-receptors (orange).

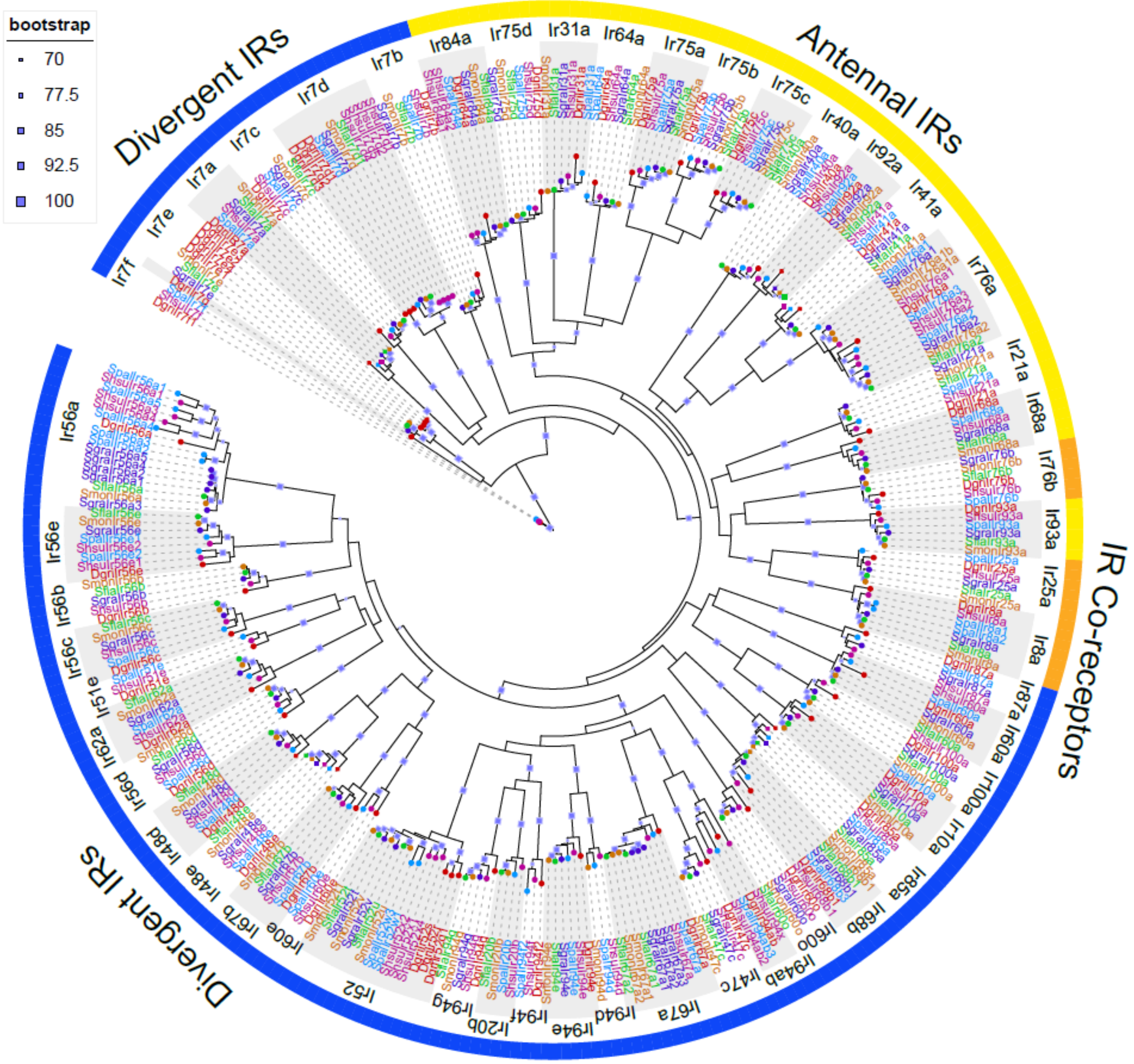

**Figure S4. Phylogeny of odorant binding receptors.** A midpoint rooted ML gene tree inferred and visualized using RAxML (Stamatakis 2006) and iTOL (Letunic and Bork 2021). Bootstrap support is given the size of squares at the midpoint of each branch, with only those >70 shown. Species are coded by font and tip color: *D. grimshawi* (red), *S. pallida* (blue), *S. hsui* (magenta), *S. montana* (yellow), *S. flava* (green), *S. graminum* (purple). Gene orthology groups are indicated by alternating branch shading. Outer colored clade labels group genes by known OBP classes based on the number of conserved cysteine residues: classic (6 cysteines, green), plus-C (>6 cysteines, blue), minus-C (<6 cysteines, purple).

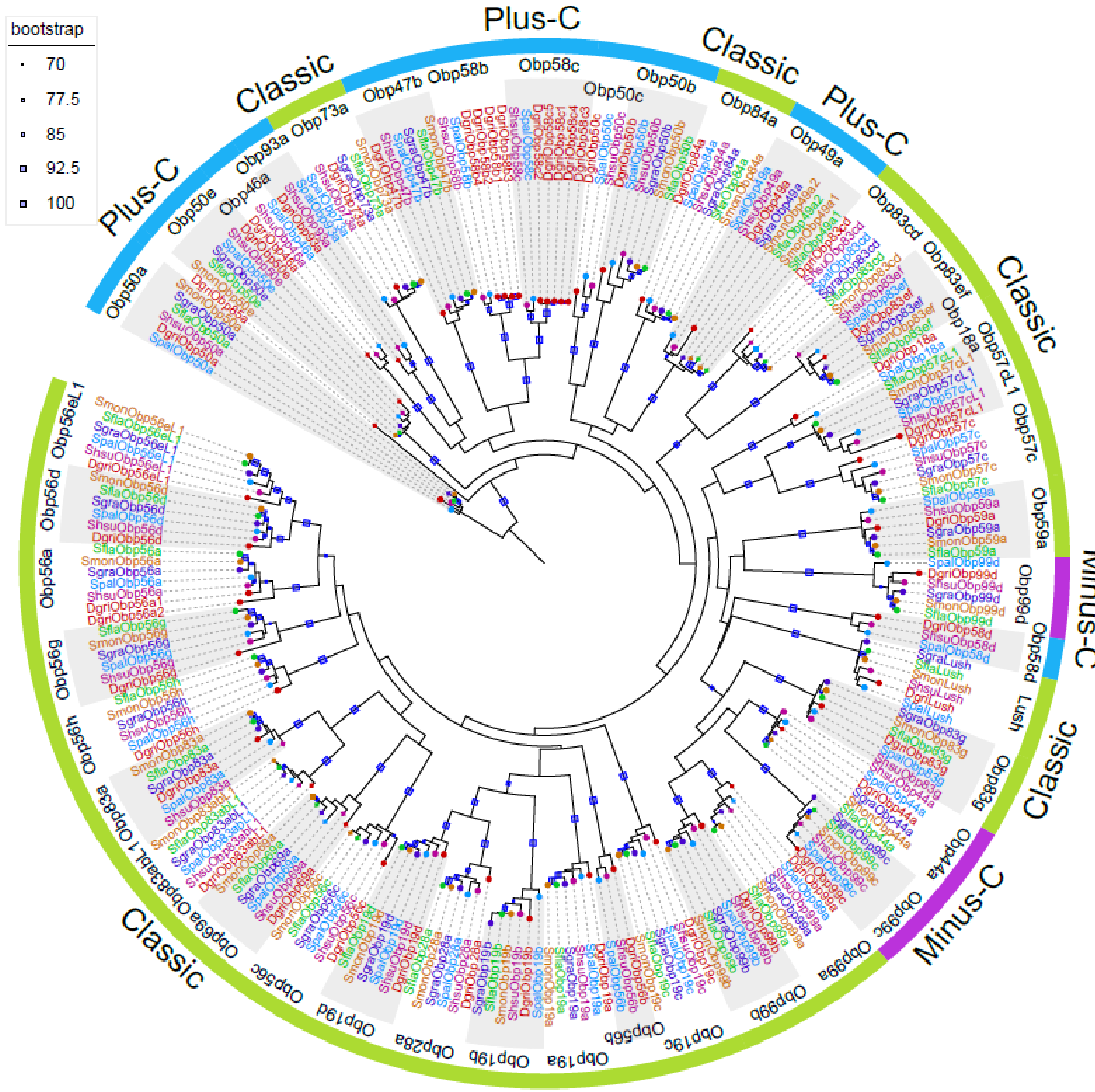

**Figure S5. Phylogeny of olfactory receptors.** A midpoint rooted ML gene tree inferred and visualized using RAxML (Stamatakis 2006) and iTOL (Letunic and Bork 2021). Bootstrap support is given the size of squares at the midpoint of each branch, with only those >70 shown. Species are coded by font and tip color: *D. grimshawi* (red), *S. pallida* (blue), *S. hsui* (magenta), *S. montana* (yellow), *S. flava* (green), *S. graminum* (purple). Gene orthology groups are indicated by alternating branch shading.

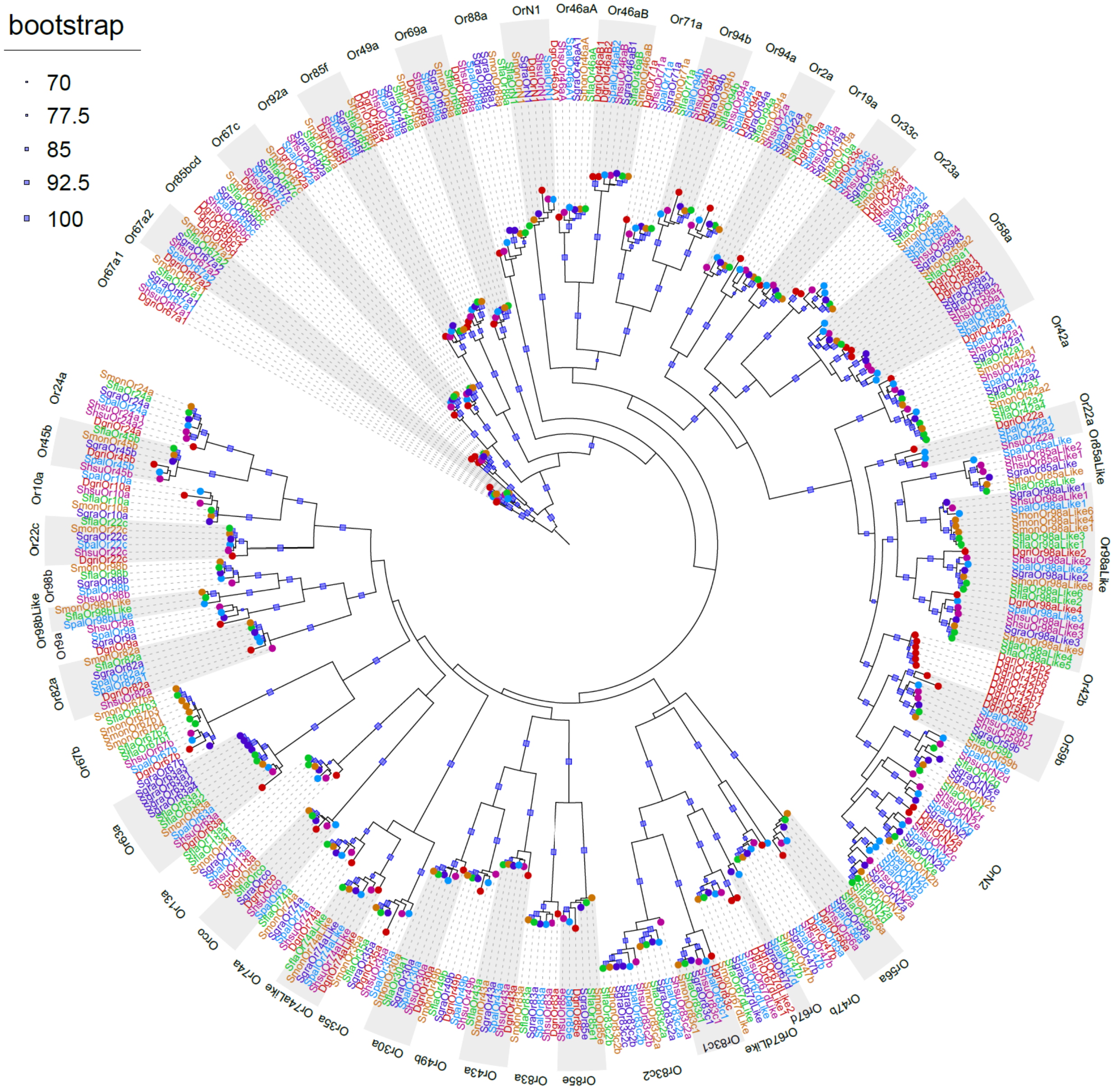

**Figure S6. Phylogeny of degenerin/epithelial sodium channels.** A midpoint rooted ML gene tree inferred and visualized using RAxML (Stamatakis 2006) and iTOL (Letunic and Bork 2021). Bootstrap support is given the size of squares at the midpoint of each branch, with only those >70 shown. Species are coded by font and tip color: *D. grimshawi* (red), *S. pallida* (blue), *S. hsui* (magenta), *S. montana* (yellow), *S. flava* (green), *S. graminum* (purple). Gene orthology groups are indicated by alternating branch shading.

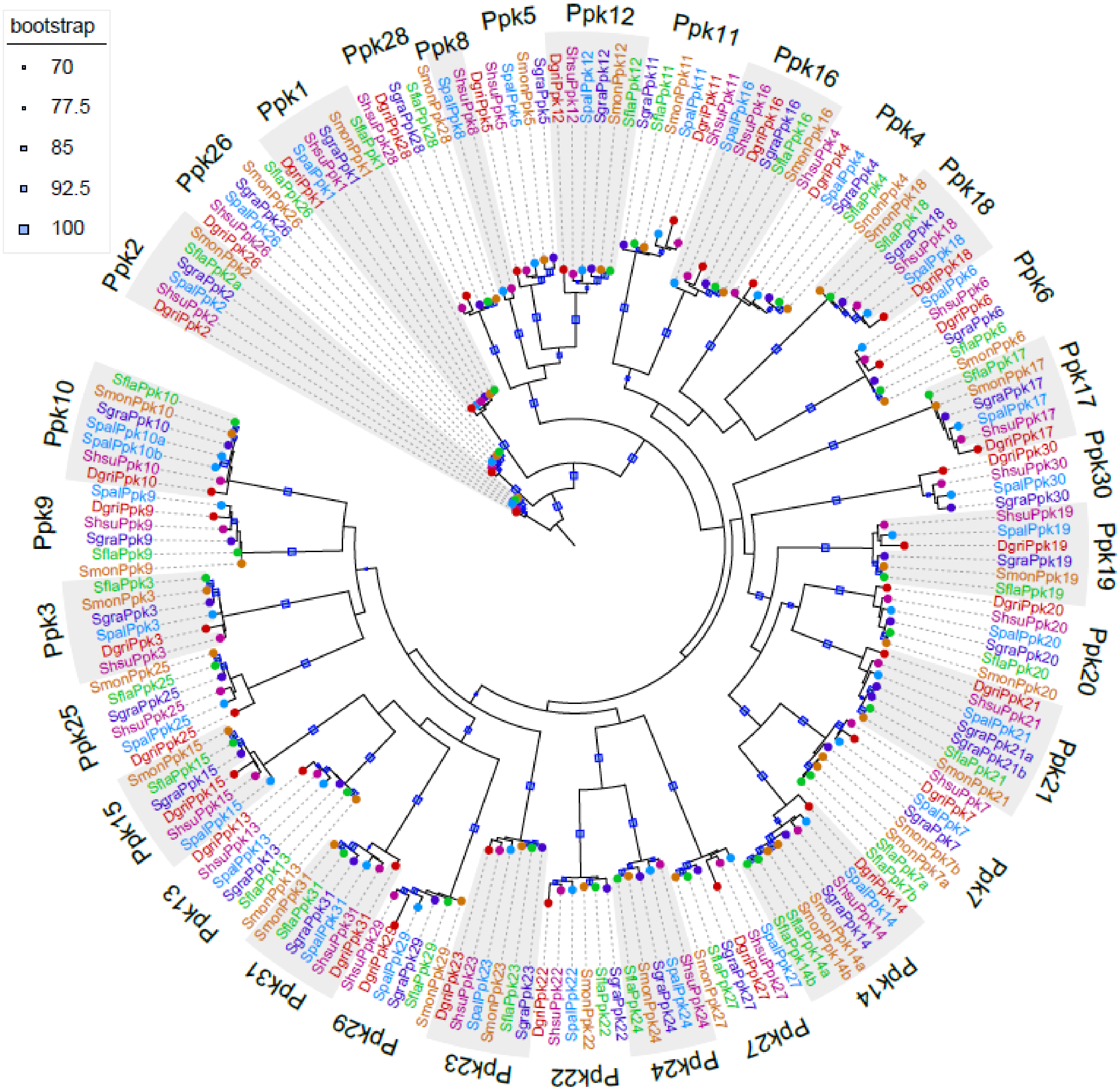

**Figure S7. Phylogeny of transient receptor potential channels.** A midpoint rooted ML gene tree inferred and visualized using RAxML (Stamatakis 2006) and iTOL (Letunic and Bork 2021). Bootstrap support is given the size of squares at the midpoint of each branch, with only those >70 shown. Species are coded by font and tip color: *D. grimshawi* (red), *S. pallida* (blue), *S. hsui* (magenta), *S. montana* (yellow), *S. flava* (green), *S. graminum* (purple). Gene orthology groups are indicated by alternating branch shading.

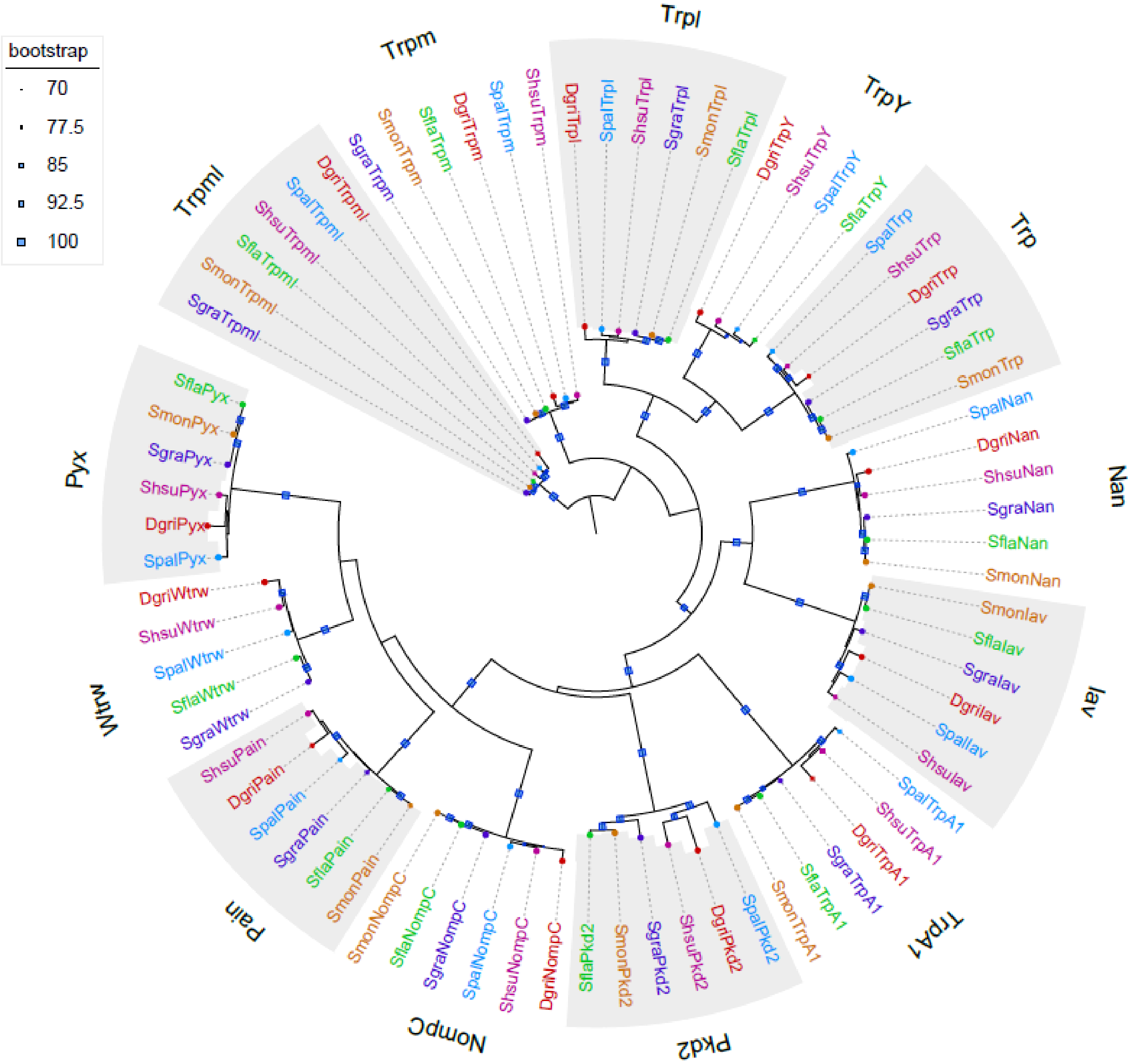

**Figure S8. Phylogeny of glutathione S-transferases.** A midpoint rooted ML gene tree inferred and visualized using RAxML (Stamatakis 2006) and iTOL (Letunic and Bork 2021). Bootstrap support is given the size of squares at the midpoint of each branch, with only those >70 shown. Species are coded by font and tip color: *D. grimshawi* (red), *S. pallida* (blue), *S. hsui* (magenta), *S. montana* (yellow), *S. flava* (green), *S. graminum* (purple). Gene orthology groups are indicated by alternating branch shading. Outer colored clade labels group genes by known GST classes: microsomal (yellow), zeta (purple), omega (green), epsilon (orange), delta (red), sigma (magenta), and theta (blue).

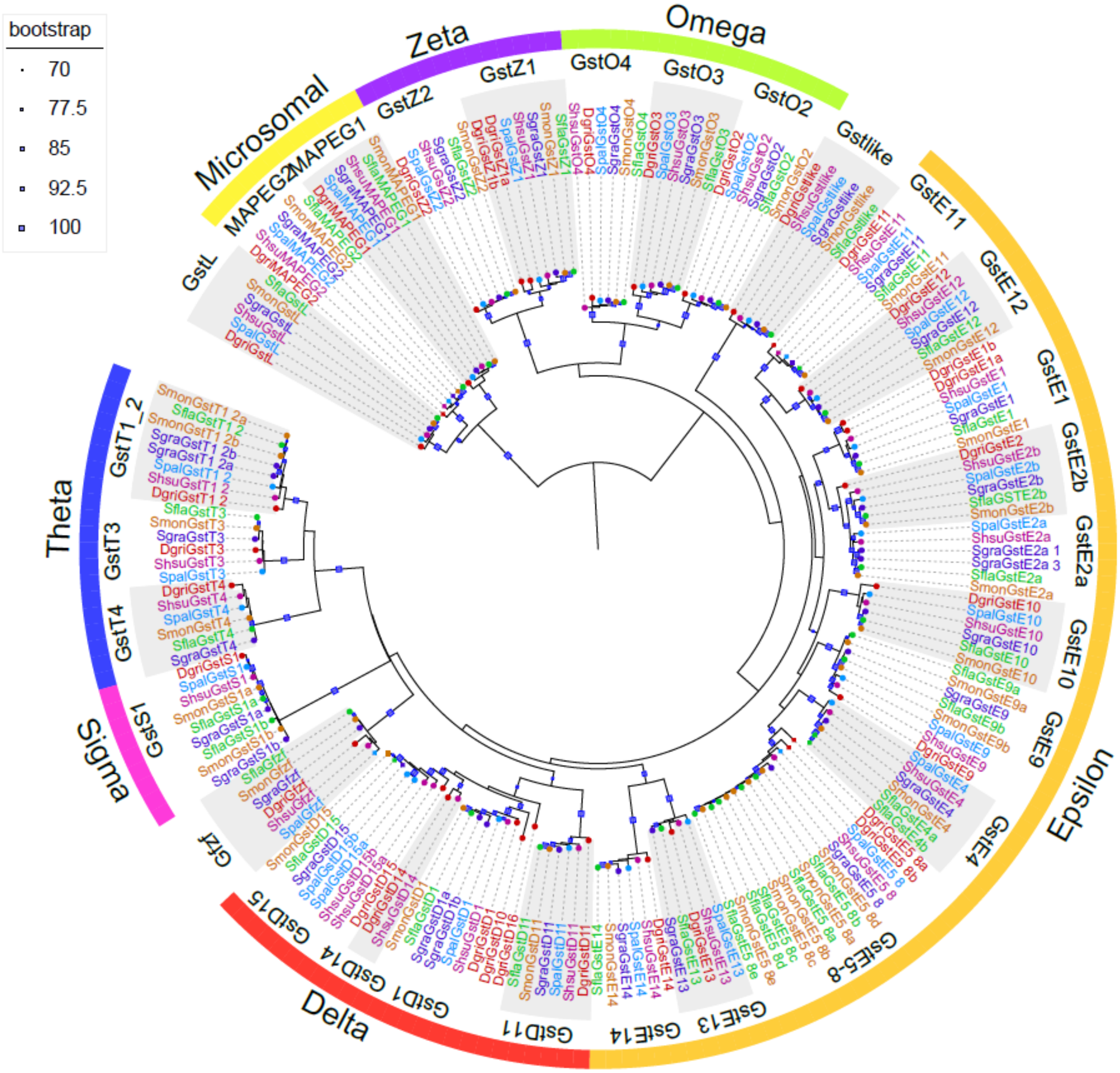

**Figure S9. Phylogeny of cytochrome CYP450s.** A midpoint rooted ML gene tree inferred and visualized using RAxML (Stamatakis 2006) and iTOL (Letunic and Bork 2021). Bootstrap support is given the size of squares at the midpoint of each branch, with only those >70 shown. Species are coded by font and tip color: *D. grimshawi* (red), *S. pallida* (blue), *S. hsui* (magenta), *S. montana* (yellow), *S. flava* (green), *S. graminum* (purple). Gene orthology groups are indicated by alternating branch shading.

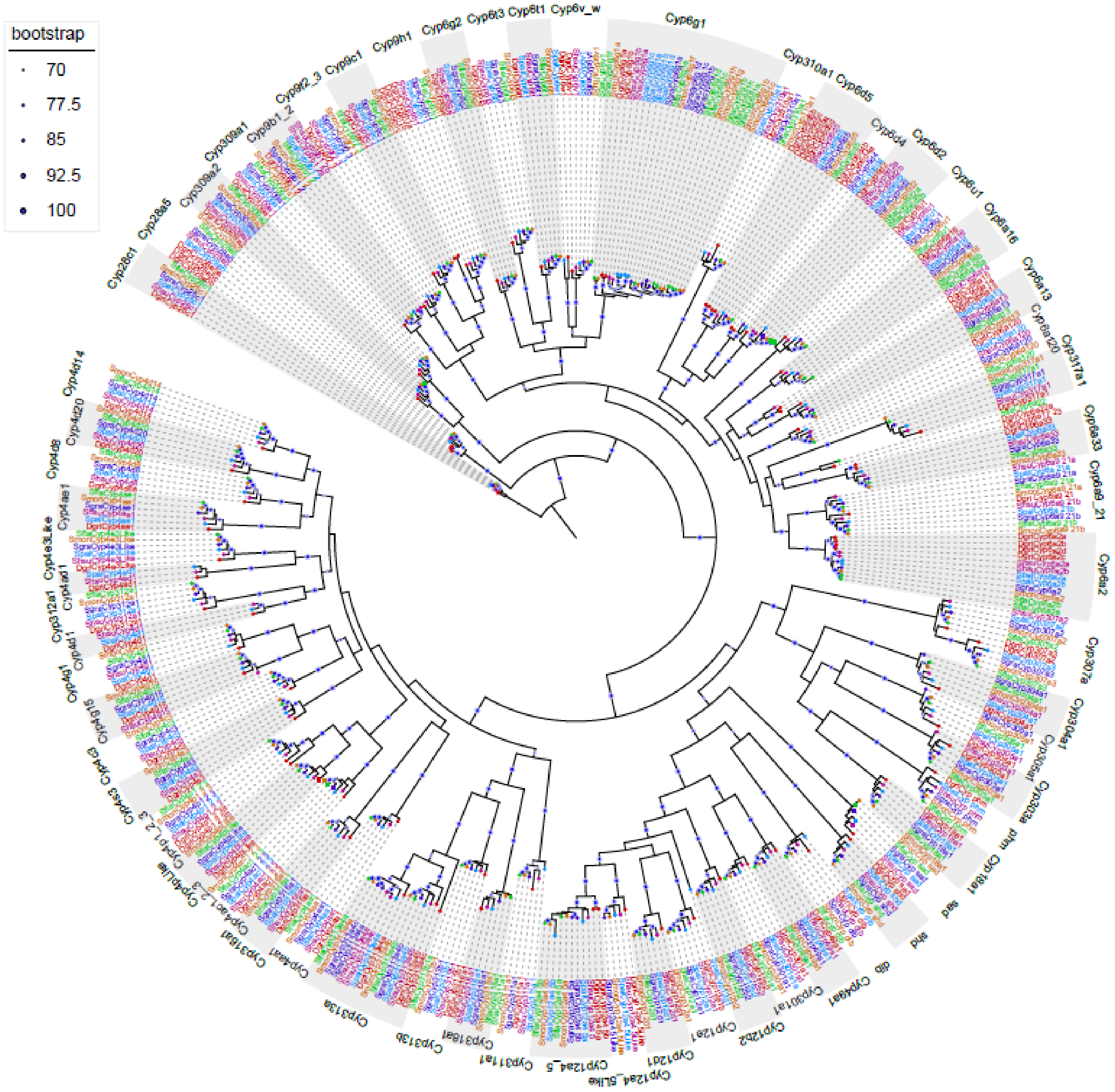

**Figure S10. Phylogeny of UDP glucuronosyltransferases UGTs.** A midpoint rooted ML gene tree inferred and visualized using RAxML (Stamatakis 2006) and iTOL (Letunic and Bork 2021). Bootstrap support is given the size of squares at the midpoint of each branch, with only those >70 shown. Species are coded by font and tip color: *D. grimshawi* (red), *S. pallida* (blue), *S. hsui* (magenta), *S. montana* (yellow), *S. flava* (green), *S. graminum* (purple). Gene orthology groups are indicated by alternating branch shading.

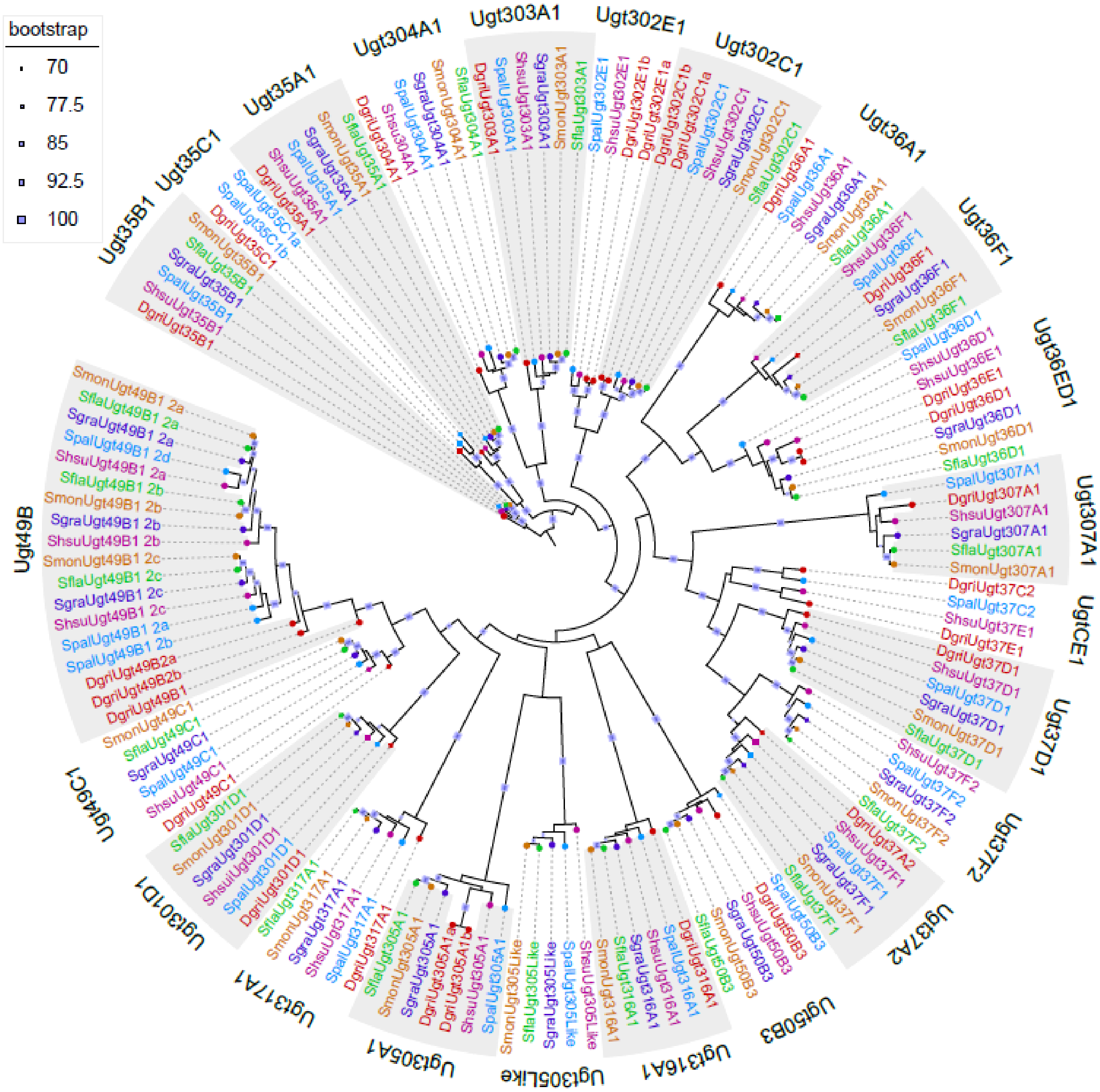

**Figure S11. Higher rates of gene turnover among herbivorous *Scaptomyza* are not due to differences in branch lengths.** (a) Phylogeny (Matsunaga et al. 2022) highlighting branch lengths of *S. flava* and *S. graminum* versus *D. melanogaster* and *D. erecta*, the latter pair being comparable in branch lengths to the former pair. (b) CAFE results when the foreground was either the clade of *D. melanogaster* and *D. erecta* or the clade of *S. flava* and *S. graminum*. All chem = all chemosensory genes. all detox = all detoxification genes. Random = random set of 200 orthology groups.

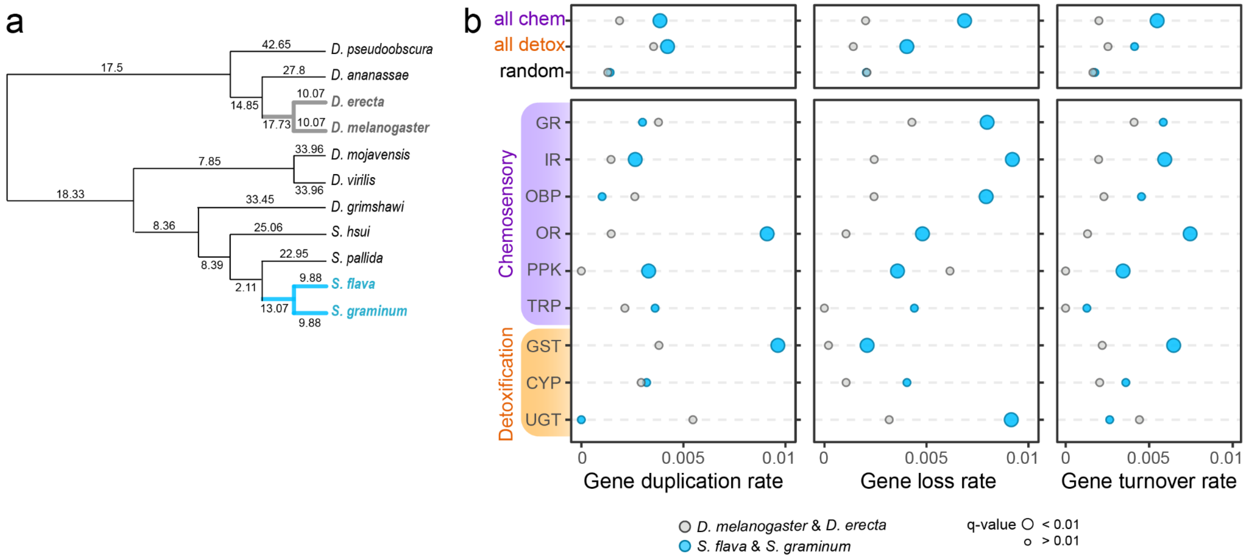

**Figure S12. Dramatic loss of "Plus-C" odorant binding proteins at the base of herbivorous drosophilid clade**. (a) CAFE gene family expansion/contraction analysis for all OBPs – gains are shown above branches in blue, losses are shown below in red. Circles indicate feeding ecology: green = herbivorous, gray = non-herbivorous. (b) Gene counts by OBP class show that the majority of herbivore-specific losses are among the Plus-C OBPs. (c) Phylogeny of Plus-C OBPs. Thick branches indicate lineages with herbivore-specific losses. Node values indicate bootstrap support.

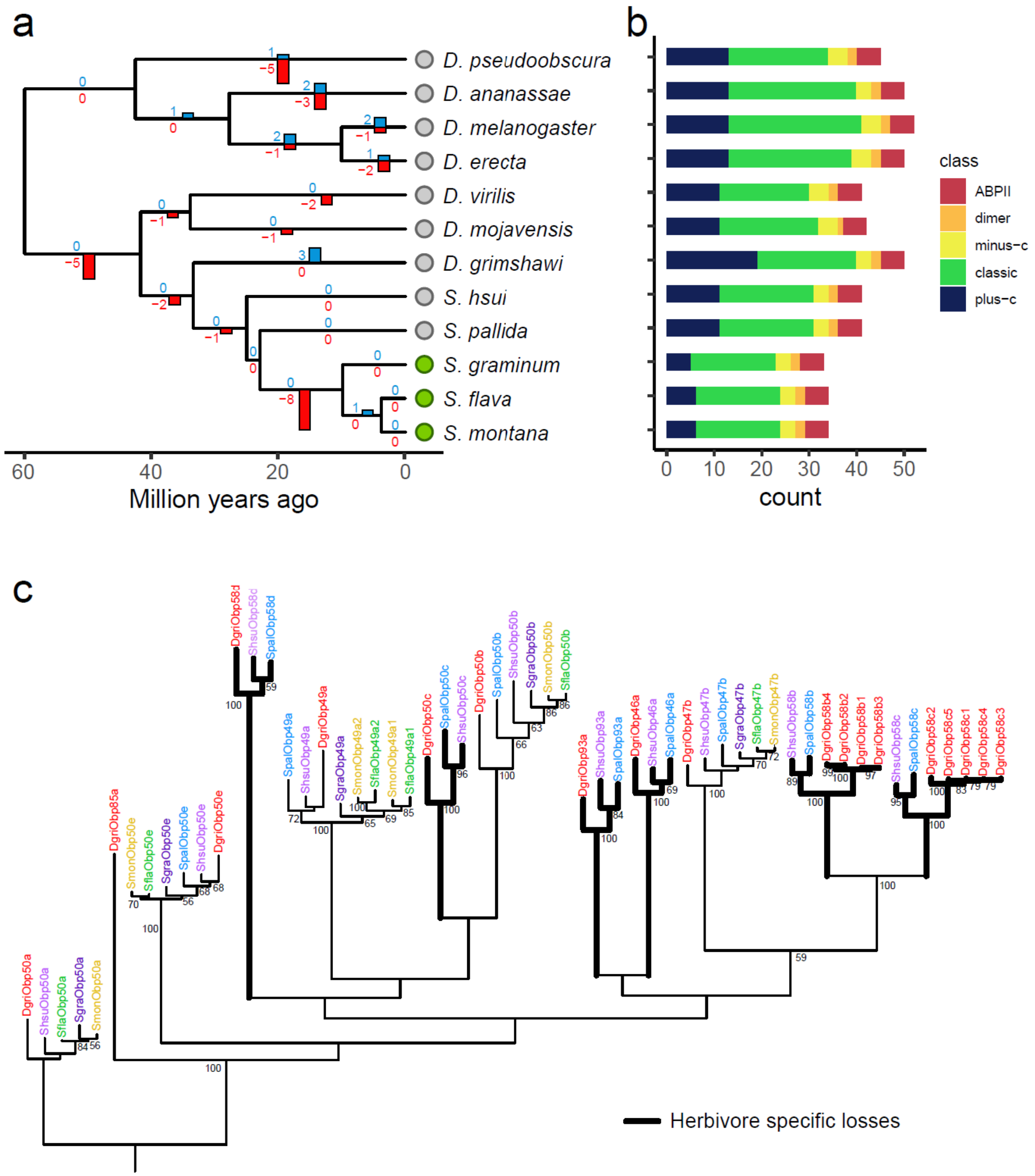

**Figure S13. Chemosensory genes duplicated, lost, or under changing selection regimes in the branch at the base of the herbivorous *Scaptomyza* lineage.** Localization map of chemosensory gene products in *D. melanogaster* adults and larvae. Only shown are those that have been lost in all herbivores (gray), duplicated (underlined), or experienced relaxed purifying selection (orange), stronger purifying selection (blue), or positive selection (red). Dorsal organ (DO); terminal organ (TO);

dorsal, ventral, and posterior pharyngeal sense organ (DPS, VPS, PPS, respectively); labral sense organ (LSO); ventral cibarial sense organ (VCSO).

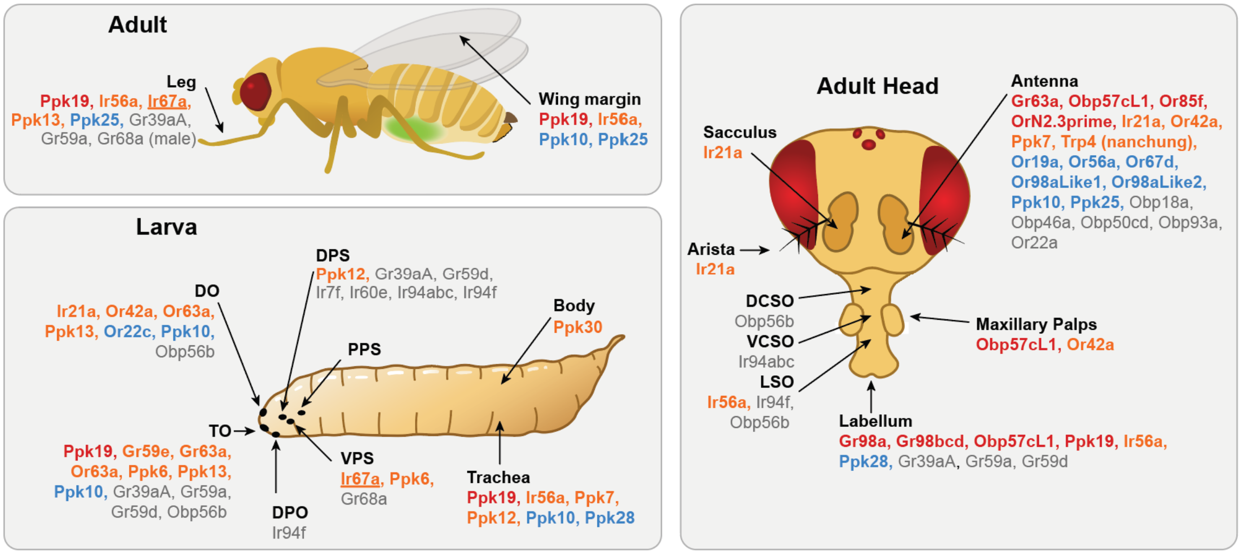

**Figure S14. Output branch labels for CAFE analysis.** Branch IDs correspond to those listed in Table S6. Herbivorous taxa in green font.

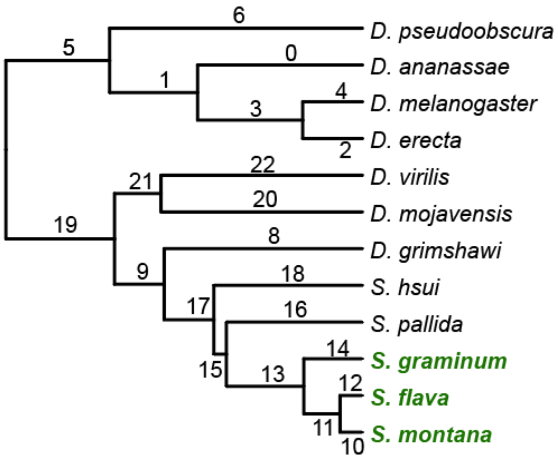

### LIST OF SUPPORTING DATASETS

**Dataset 1.** Gene coordinates and identification numbers of chemosensory and detoxification genes curated from *Drosophila* and *Scaptomyza* genome assemblies.
**Dataset 2.** Data and code used to analyze gene family evolutionary dynamics presented (Figure 2), including curated gene count matrices supplied as input and the raw and parsed output files from the CAFE analysis.
**Dataset 3.** Data, output, and code for rates of molecular evolution (*dN/dS*, PAML) (Table 1).
**Dataset 4.** Sequence alignment and phylogeny used to characterize the taxonomic diversity of herbivorous *Scaptomyza* (Figure 1b).
